# Generalist *Eimeria* species in rodents: multilocus analyses indicate inadequate resolution of established markers

**DOI:** 10.1101/690487

**Authors:** Víctor Hugo Jarquín-Díaz, Alice Balard, Anna Mácová, Jenny Jost, Tabea Roth von Szepesbéla, Karin Berktold, Steffen Tank, Jana Kvičerová, Emanuel Heitlinger

## Abstract

Intracellular parasites of the genus *Eimeria* are described as tissue/host specific. Phylogenetic classification of rodent *Eimeria* suggested that some species have a broader host range than previously assumed. We explore if *Eimeria* spp. infecting house mice are misclassified by the most widely used molecular markers due to a lack of resolution, or if, instead, these parasite species are indeed infecting multiple host species.

With the commonly used markers (18S/COI), we recovered monophyletic clades of *E. falciformis* and *E. vermiformis* from *Mus* that included *E. apionodes* identified in other rodent host species (*Apodemus* spp., *Myodes glareolus*, and *Microtus arvalis*). A lack of internal resolution in these clades could suggest the existence of a species complex with a wide host range infecting murid and cricetid rodents. We question, however, the power of COI and 18S markers to provide adequate resolution for assessing host specificity. In addition to the rarely used marker ORF470 from the apicoplast genome, we present multilocus genotyping as an alternative approach. Phylogenetic analysis of 35 nuclear markers differentiated *E. falciformis* from house mice from isolates from *Apodemus* hosts. Isolates of *E. vermiformis* from *Mus* are still found in clusters interleaved with non-*Mus* isolates, even with this high resolution data.

In conclusion, we show that species-level resolution should not be assumed for COI and 18S markers in Coccidia. Host-parasite co-speciation at shallow phylogenetic nodes, as well as contemporary coccidian host ranges more generally, are still open questions that need to be addressed using novel genetic markers with higher resolution.

## 1. Introduction

Parasites are often categorized as either generalists or specialists depending on their host range. In natural systems, however, the distinction between generalist and specialist should be based not only on the capacity to infect one or more host species, but also on their prevalence and intensity of infection in different host genotypes or species (Combes 2001; Leggett et al. 2013; Schmid-Hempel 2011). Under this framework, the difference between generalist and specialist represents a continuum (Schmid-Hempel 2011). Hypotheses, assumptions and predictions concerning host-parasite interactions from evolutionary (Adamson and Caira 1994; Combes 2001; Poulin, Krasnov, and Mouillot 2011; Schmid-Hempel 2011), ecological (Fenton and Brockhurst 2008; Forbes, Muma, and Smith 2002; Kassen 2002) and mechanistic (Rathore et al. 2003) perspectives depend on the placement of parasite species in this continuum.

Coccidians of the genus *Eimeria* have been described as monoxenous, intracellular parasites (Becker 1934; Long and Joyner 1984; Marquardt 1981). Their assumed high degree of host and tissue specificity is extensively used to delineate species. It is not clear, however, whether host specificity is the same for *Eimeria* species infecting hosts in different clades. *Eimeria* species of rodents show a degree of specificity (Ball and Lewis 1984; Duszynski 2011; De Vos 1970; Wilber et al. 1998) but individual isolates can experimentally infect different species and even genera of rodents (Levine and Ivens 1988; Upton et al. 1992).

Descriptions of *Eimeria* species are based on the size and shape of sporulated oocysts and their internal structures. The life cycles of a few species has additionally been studied and data on their dynamics (e.g. the patent period, the time before oocysts are shed in faeces) are available (Duszynski, Eastham, and Yates 1982; Hnida, Wilson, and Duszynski 1998; Todd and Lepp 1971; Lainson and Shaw 1990; Levine and Ivens 1965; Mesfin and Bellamy 1978; Todd and Hammond 1968; Turner, Penzhorn, and Getz 2016; Wash, Duszynski, and Yates 1985). For field studies, the morphology of sporulated oocysts alone is considered insufficient to infer species identity because of inadequate reference descriptions (MacPherson and Gajadhar 1993; Tenter et al. 2002).

Genetic markers from nuclear (nu) and mitochondrial (mt) genomes, and less frequently also of the and apicoplast (ap) genome, have been used to complement morphological taxonomy with phylogenetic analyses (Hnida and Duszynski, 1999a; Hnida and Duszynski, 1999b; Kvičerová et al., 2011; Ogedengbe et al., 2015; Zhao and Duszynski, 2001a). Based on the assumption of host specificity of individual *Eimeria* species, phylogenetic analysis of nuclear small subunit ribosomal (18S) rDNA and cytochrome c Oxidase I (COI) fragments supports predominant host-parasite co-speciation (Ogedengbe et al. 2018). Species infecting rodents, however, are found in two separate clades, generating marked discrepancy between parasite and host phylogeny at deeper nodes (Kvičerová and Hypša 2013). At shallow nodes of the phylogeny for rodent coccidians, host generalism has been suggested (Mácová et al., 2018). Host specificity of *Eimeria* species infecting rodents is not as undisputed as in other hosts such as poultry (Barta et al. 1997) or rabbits (Kvičerová, Pakandl, and Hypša 2008). Kvičerová and Hypša (2013) suggested that adaptation rather than co-speciation is shaping rodent-*Eimeria* co-phylogenies. Mácová et al. (2018) added that host ecology and distribution may favour host-switches among closely related rodent species. A high specificity of *E. apionodes* naturally infecting *Apodemus flavicollis* was originally suggested based on failed attempts to experimentally infect other rodents: *Myodes* (C*lethrionomys) glareolus, Microtus arvalis*, or *Mus musculus* (Pellérdy 1954). It is, however, unclear if this result holds for the multiple isolates that have been assigned as *E. apionodes*.

We studied wild populations of *Mus musculus* and other rodents to assess the diversity of *Eimeria* isolates at shallow depth of phylogenetic relationships. We test host specificity based on phylogenetic analysis using established markers (nu 18S, mt COI and ap ORF470). We question in how far these markers are polymorphic enough to resolve between genetic clusters with different host usage (and whether a negative result for genetic differentiation therefore suggests generalism). We develop and apply multilocus sequence typing to disentangle relationships unresolved by 18S and COI markers.

## 2. Material and methods

### 2.1. Origin of samples

DNA was extracted from the colon content or gastrointestinal tissue of house mice (*Mus musculus*) infected with *Eimeria*. These samples came from rodents captured in farms and private properties in the German federal states of Mecklenburg-Vorpommern, Bavaria and Brandenburg (capture permit No. 2347/35/2014) and in Bohemia (Czech Republic) between 2014 and 2017 (Jarquín-Díaz et al. 2019) (Fig. 1A). Additionally, DNA from gastrointestinal tract, tissue or faeces of *Apodemus* spp. from different regions in Europe (including areas overlapping with those sampled for house mice) were also included (Mácová et al., 2018) (Fig. 1B) (supplementary data S1).

**Figure 1.**
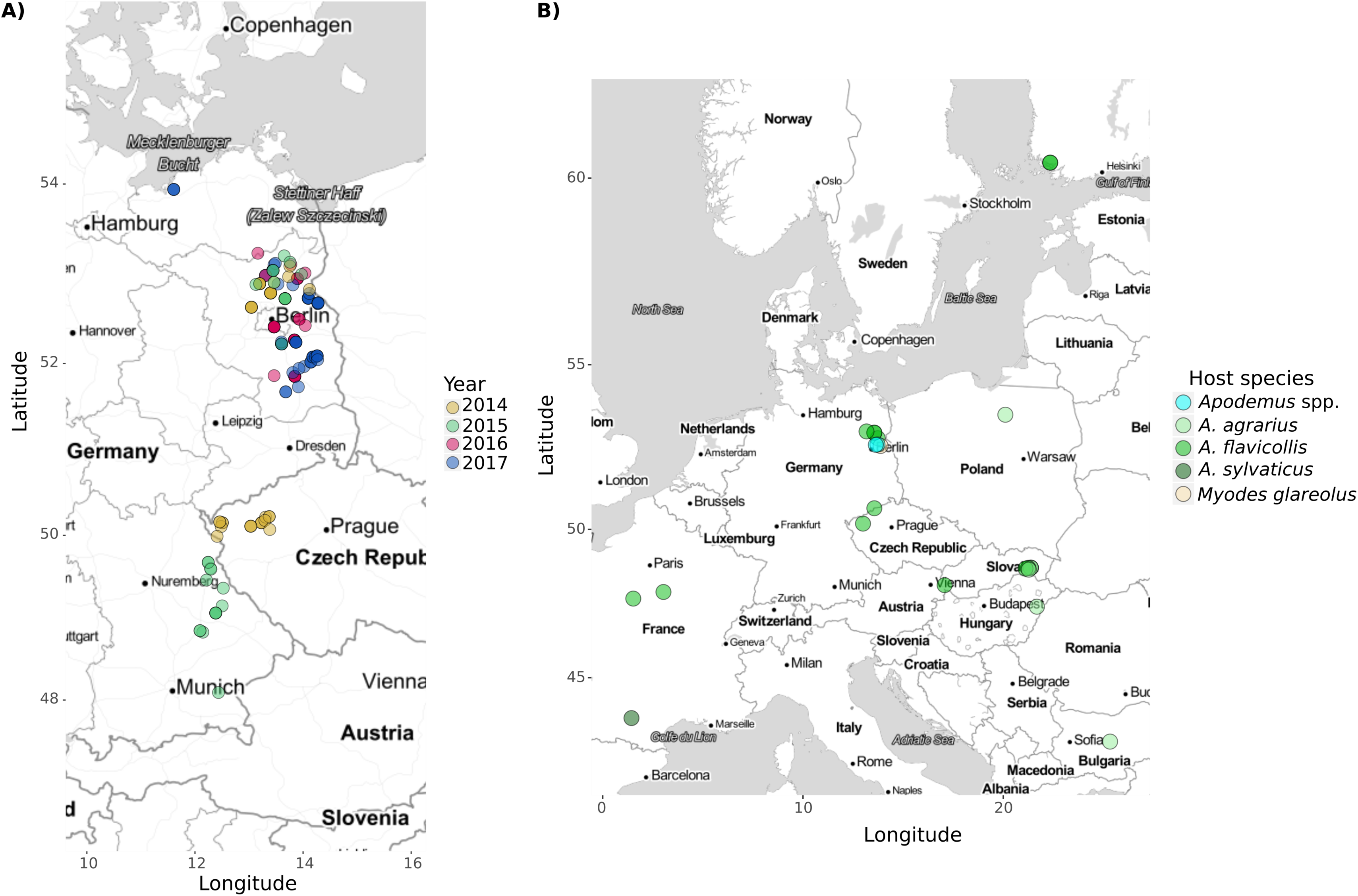
Location of rodent samples. A) *Mus musculus* samples were collected from the German federal states of Mecklenburg-Vorpommern, Bavaria and Brandenburg and in Bohemia (Czech Republic) between 2014 and 2017. Colour code indicate the year of collection. B) Non-*Mus* samples were collected from different countries within Europe. Colour in the points indicate the host species.

### 2.2. Host identification

Rodents were first identified visually based on their morphology. Identification of *Mus musculus* at the sub-species level was confirmed based on a set of previously described markers (Ďureje et al. 2012). In order to confirm the species of non-*Mus* rodents, a fragment of cytochrome *b* (∼ 900 bp) was amplified from host DNA. PCRs were performed according to the protocols described by Reutter et al. (2003) for *Apodemus* spp., Abramson et al. (2009) (primers UCBO_F/LM_R) and Jaarola and Searle (2002) (primers L14641M/H15408M) for rodents belonging to the subfamily Arvicolinae (*Myodes* spp. And *Microtus* spp.).

### 2.3. PCR amplification (ap tRNA, nu 18S rDNA, mt COI and ap ORF470)

For phylogenetic analysis, nuclear small subunit ribosomal DNA (18S; ∼1,500 bp), a fragment of the mitochondrial cytochrome c oxidase subunit I (COI; ∼ 800 bp) gene and apicoplast ORF470 (∼ 800 bp) were amplified using primers previously reported by Kvičerová et al. (2008), Ogedengbe et al. (2011) and Zhao and Duszynski (2001b), respectively.

When COI failed to amplify with this protocol, an alternative pair of primers was used: Eim_COI_M_F (ATGTCACTNTCTCCAACCTCAGT) and Eim_COI_M_R (GAGCAACATCAANAGCAGTGT). These primers amplify a ∼700 bp fragment of COI and were designed based on the mitochondrial genome of *E. falciformis* (CM008276.1) (Heitlinger et al. 2014; Jarquín-Díaz et al. 2019).

PCR reactions were carried out in a Labcycler (SensoQuest GmbH, Göttingen, Germany) using 0.025 U/µL of DreamTaqTM DNA Polymerase (Thermo Scientific, Waltham, USA), 1X DreamTaq Buffer, 0.5 mM dNTP Mix, 0.25 µM from each primer and 1 – 20 ng/µL of DNA template in 25 µL reaction. A concentration of 0.25 mM dNTP Mix and a supplementation with 0.5 mM MgCl_2_ was used for the ap ORF470 amplification. The thermocycling protocol consisted of 95 °C initial denaturation (4 min) followed by 35 cycles of 92 °C denaturation (45 s), annealing at 52°C (30 s/Eim_COI); 53 °C (45 s/18S); 55 °C (30 s/COI); 50 °C (45 s/ORF470); 72 °C extension 90 s (18S/ORF470), 20 s (COI/Eim_COI), and a final extension at 72 °C (10 min). DNA from oocysts of *E. falciformis* BayerHaberkorn1970 and DNA from colon content of a non-infected lab (NMRI) mouse were used as positive and negative controls, respectively.

All PCR products from nu 18S, mt COI and ap ORF470 of the expected size were purified using the SAP-Exo Kit (Jena Bioscience GmbH, Jena, Germany), and sequenced in both directions by LGC Genomics (Berlin, Germany). Quality assessment and sequence assembly was performed in Geneious v6.1.8. All sequences were submitted to the NCBI GenBank database (Accession numbers: nu 18S rDNA [MH751925-MH752036, MK246860-MK246868 and MK625202-MK625210]; mt COI [MH777467-MH777593, MH755302-MH755324, MK257106-MK257114 and MK631866-MK631868] and ap ORF470 [MH755325-MH755450, MK257115-MK257125 and MK631869-MK631884]).

### 2.4. Phylogenetic analysis and inference of intraspecific genetic diversity

Datasets for each gene and a concatenated alignment (nu 18S, mt COI and ap ORF470) were created adding closely related reference sequences available in the GenBank (supplementary data S2).

Protein coding sequences (mt COI and ap ORF470) were aligned by translation using the Multiple Align algorithm and translation frame 1 with the genetic code for “mold protozoan mitochondrial”, 18S sequences were aligned using MUSCLE (Edgar 2004), both through Geneious v6.1.8.

Phylogenetic trees for all datasets were constructed using Maximum Likelihood (ML) and Bayesian inference (BI) methods, implemented in PhyML v3.0 (Guindon et al. 2010) and MrBayes v3.2.6 (Huelsenbeck and Ronquist 2001; Ronquist et al. 2012), respectively. Sequence evolution models most appropriate for each dataset were determined in JModelTest v2.1.10 (Posada 2008). For ML trees, a bootstrap analysis with 1,000 replicates was performed, whereas MCMC for BI was run with two cold and two hot chains for 1,000,000 generations or until the split freq value was below 0.05. The concatenated dataset was analysed using partitions and locus-specific models. Trees were visualized with FigTree v1.4.2 (Rambaut 2012). A haplotype network of mt COI sequences was inferred using a codon-based alignment trimmed to 500 bp available for all isolates. Haplotypes frequencies were calculated and a network was constructed with the R package “pegas” v0.11 (Paradis et al. 2018).

### 2.5. Multimarker genotyping PCR and high throughput sequencing

Samples positive for *E. falciformis* and *E. vermiformis* from *Mus musculus* and *Eimeria* sp. from *Apodemus* with indistinguishable 18S and COI sequences were used for a multimarker amplification using the microfluidics PCR system Fluidigm Access Array 48 × 48 (Fluidigm, San Francisco, California, USA). We used target specific primers (supplementary data S3) that were designed based on the genome of *E. falciformis* (Heitlinger et al. 2014) to amplify exons of nuclear genes (supplementary data S4) and coding and non-coding regions from the apicoplast genome (supplementary data S5). Library preparation was performed according to the protocol Access Array Barcode Library for Illumina Sequencers (single direction indexing) as described by the manufacturer (Fluidigm, San Francisco, California, USA). The library was purified using Agencourt AMPure XP Reagent beads (Beckman Coulter Life Sciences, Krefeld, Germany). Quality and integrity of the library was confirmed using the Agilent 2200 Tape Station with D1000 ScreenTapes (Agilent Technologies, Santa Clara, California, USA). Sequences were generated at the Berlin Center for Genomics in Biodiversity Research (BeGenDiv) on the Illumina MiSeq platform (Illumina, San Diego, California, USA) in two runs, one using “v3 chemistry” with 600 cycles, the other “v2 chemistry” with 500 cycles. All sequencing raw data can be accessed through the BioProject PRJNA548431 in the NCBI Short Read Archive (SRA).

### 2.6. Bioinformatic analysis of multilocus sequence typing

Screening and trimming of sequencing reads was performed using the package dada2 v1.2.1 (Callahan et al. 2016). All reads were trimmed to 245 bases, while allowing a maximum of 4 expected errors (maxEE). Sorting and assignment to amplicons was performed with the package MultiAmplicon v0.1 (Heitlinger, 2019) and the most abundant sequence was recorded for each marker in each sample (recording but disregarding minority sequence in non-clonal infection for further analysis; see supplementary data S6). Sequences were aligned using the function “AlignSeqs” from the package DECIPHER v2.10.0 (Wright 2016) and non-target sequences were excluded from alignments if >20% divergence was observed with other sequences (such as in cases off-target amplification of mostly bacterial sequences). Alignments were controlled for the absence of insertions/deletions (indels) that distort the open reading frame. Prevalent multiple-of-3-mere indels corresponding to homopolymeric amino acid repeats (HAARs; Heitlinger et al. 2014) of diverse length were coded as missing data due to their unclear model of evolution. The function “dudi.pcr” from the packages ade4 v1.7-13 (Dray and Dufour 2015) and adegenet v2.1.1 (Jombart 2008) was used to visualize genetic distances between samples based on all markers. The code for this pipeline is available at https://github.com/VictorHJD/AA_Eimeria_Genotyping.

The alignments of the concatenated sequences were then exported. The number of informative sites was summarized using the tool DIVEIN (Deng et al. 2010) and phylogenetic trees were computed by Bayesian inference in MrBayes v3.2.6 (Huelsenbeck and Ronquist, 2001; Ronquist et al., 2012). A partitioned model was implemented to estimate the tree considering each gene separately. The analysis was performed with two runs, with 1,000,000 generations leading to a split frequency value below 0.05, and 200,000 generations were discarded as burn-in when estimating posterior probability. Additionally, Maximum Likelihood trees were inferred with 1,000 bootstrap replicates in PhyML v3.0 (Guindon et al. 2010).

The topology of ML and BI trees was compared and summarized into a consensus tree with minimum clade frequency threshold of 0.95 using the program SumTrees v4.3.0 (Sukumaran and Holder, 2010).

## 3. Results

### 3.1. Established markers don’t recover clades corresponding to species with different host-usage

We performed phylogenetic analyses using nuclear, mitochondrial, and apicoplast markers to assess the clustering of our sequences into groups of previously described species.

We inferred a phylogenetic tree of nu 18S based on 215 sequences (509 – 1,795 bp). Of these, 111 from parasites in house mice (*M. musculus*) (3 from ileum tissue, 16 from cecum tissue and 92 from colon content) and 18 from parasites in non-*Mus* rodents were generated in the present study (3 from ileum tissue, 3 from cecum tissue, 3 from colon content, and 9 from feces). To test for host specificity of house mouse *Eimeria* we included reference sequences from related *Eimeria* species described in murid and cricetid rodents. *Isospora* sp. sequences identified in *Talpa europaea* moles were used as an outgroup. Both ML and BI rooted trees shared the general topology (Fig. 2A).

**Figure 2.**
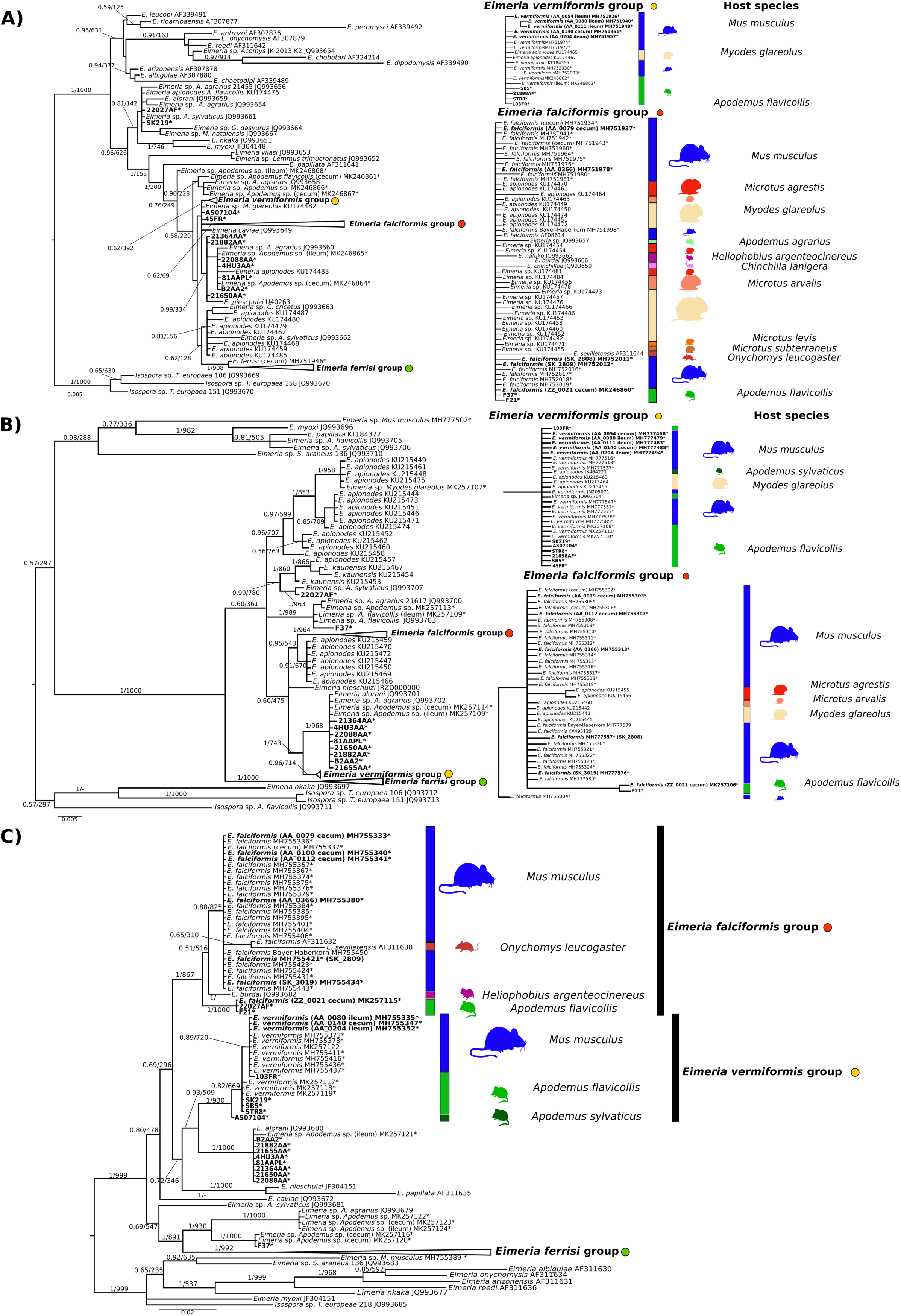
Phylogenetic trees inferred from nuclear (18S rDNA), mitochondrial (COI) and apicoplast (ORF470) sequences. Phylogenetic tree based on a) 18S rDNA, b) COI and c) ORF470 sequences. Numbers in the branches represent Bayesian posterior probability and bootstrap value. The three collapsed groups cluster *Eimeria* sequences from *M. musculus* of this study. Reference sequences from other rodents were included. The scale bar represents sequence divergence. Hosts for closely related sequences of *E. falciformis* and *E. vermiformis* are indicated in the expanded form of the group. * Represent sequences generated in the present study. Tissue of origin is indicated in brackets. Sequences in bold were included in the multi-marker phylogenetic inference.

The sequences derived from *Mus musculus* samples clustered in three well supported monophyletic groups: one comprising reference sequences of *E. falciformis* (*E. falciformis* group), another of *E. ferrisi* (*E. ferrisi* group), and the third of *E. vermiformis* (*E. vermiformis* group). All three groups, however, included sequences of *Eimeria* from other cricetid and murid hosts without showing internal sub-structure reflecting host-usage (Fig. 2A).

The phylogenetic tree of mt COI was based on 233 sequences (381 – 804 bp), 149 of which were obtained from *Eimeria* infecting house mice (3 from ileum, 16 from cecum tissue and 130 from colon content) and 12 from non- *Mus* rodents in our study (2 from ileum, 1 from cecum, 6 from colon content, and 3 from feces) (Fig. 1B). Similar to 18S, COI sequences derived from house mice clustered in three monophyletic groups including reference sequences of *E. falciformis* (*n=* 26), *E. ferrisi* (*n=* 109) and *E. vermiformis* (*n=* 13). Groups of *E. falciformis* and *E. vermiformis* also include sequences derived from *Eimeria* isolates of common voles (*Mi. arvalis*), bank voles (*My. glareolus*), short-tailed voles (*Mi. agrestis*), yellow necked mice (*A. flavicollis*) or wood mice (*A. sylvaticus*). In addition to our isolates from *M. musculus*, the *E. ferrisi* groups contains sequences of *E. burdai* and *E. nafuko*, species described from sub Saharan mole rats (*Heliophobius argenteocinereus*). Again, the clades do not show further sub-structure indicative of host usage (Fig. 2B).

A phylogenetic tree of ORF470 was based on 172 sequences (Fig. 2C) and showed a similar topology to the COI and 18S trees. Sequences derived from *Eimeria* isolates from *Mus musculus* (*n=* 125) also clustered into the same three groups. For this marker, the number of sequences available in databases from other cricetid and murid rodents is very limited, and none of the available sequences clustered within the highly supported “species clusters” of our isolates. In contrast to nu 18S and mt COI, our newly generated sequences from isolates detected in *A. flavicollis* and *A. sylvaticus* formed separate clusters that were basal to the *E. falciformis* group (n = 3), and outside of the *E. vermiformis* group (n = 4) (Fig. 2C).

To combine all available information into a single phylogenetic analysis we used a concatenated alignment. In the tree constructed from this alignment (supplementary data S7), *E. vermiformis* and *E. ferrisi* groups found in the individual marker analyses were confirmed. Sequences from the *E. falciformis* group were found in an unresolved basal position with *E. apionodes* isolates derived from *Myodes* sp. and *Microtus* sp.. This result probably indicates conflicting signals for different markers and missing data.

### 3.2. Low genetic diversity of mt COI in rodent Eimeria isolates

With the aim to estimate the genetic diversity of isolates of *Eimeria* from different rodent hosts, we constructed a haplotype network (Fig. 3) from 161 COI sequences obtained in this study combined with 59 previously published sequences (alignment of 459 bp without gaps). The network comprised 20 different haplotypes with up to 14 polymorphic nucleotide sites among them. *Eimeria* sequences from our study were distributed into five haplotypes (I, II, III, IV, and V). Haplotype I contained the largest number of sequences. It was formed by sequences from parasites of *M. musculus* and included an *E. ferrisi* reference sequence. Haplotype II, with *E. vermiformis* as a reference, was composed of a mixture of sequences from *Eimeria* isolates from *M. musculus, My. glareolus, A. flavicollis* and *A. sylvaticus*. Haplotype III also contained sequences of parasites from multiple hosts: It included *Eimeria* strains identified in *M. musculus, My. glareolus* and *Mi. arvalis* and *E. falciformis* as a reference.

**Figure 3.**
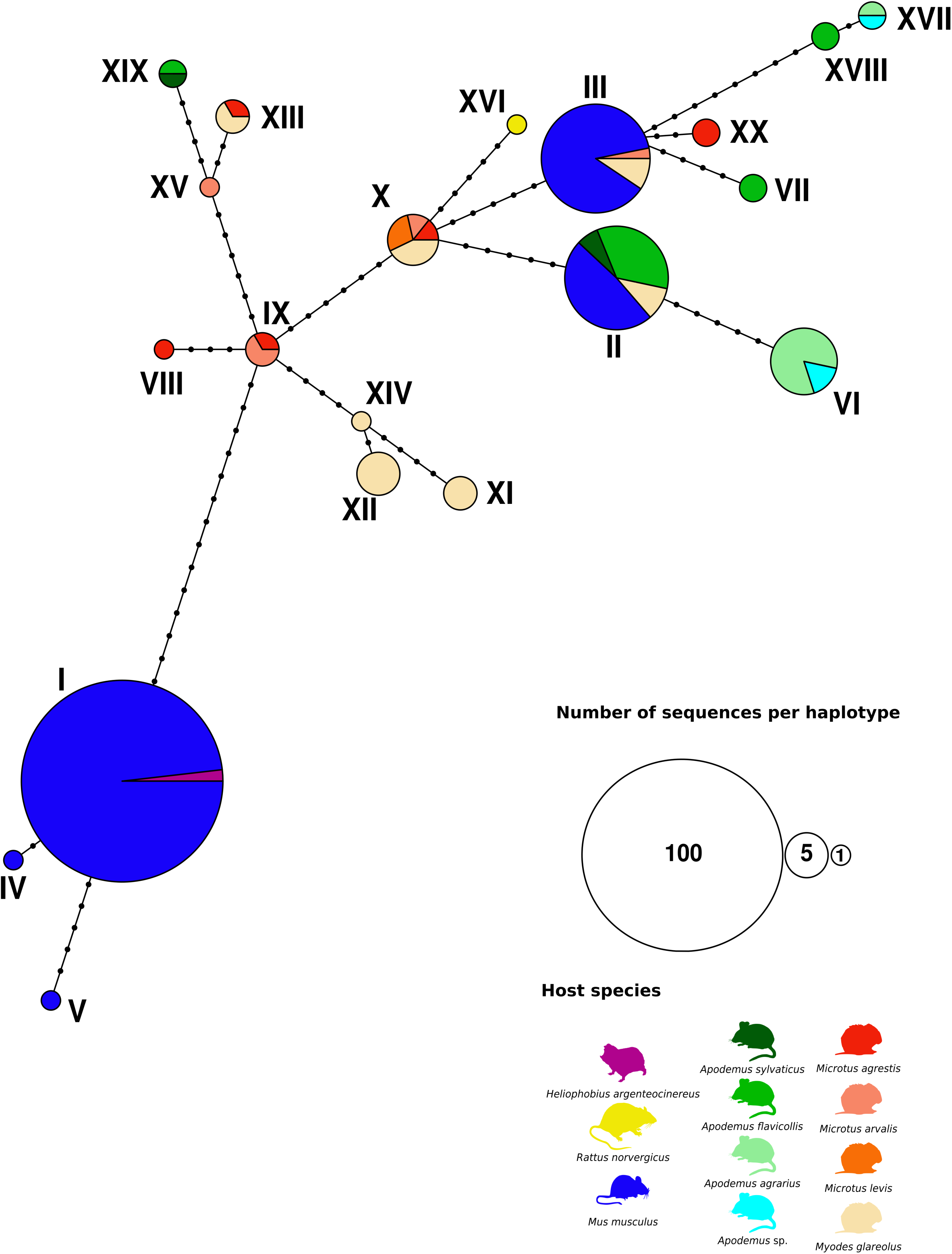
Statistical parsimony network of *Eimeria* spp. haplotypes for COI sequences. Network based on a 459 bp region of the gene coding for the mitochondrial cytochrome c oxidase from *Eimeria* isolates detected in rodents (*Mus musculus, Apodemus flavicollis, A. sylvaticus, A. agrarius*) caught in Europe. Previously published sequences from different species of *Eimeria* infecting cricetid and murid rodents were also included. Colouring of each haplotype is based on the host species from the *Eimeria* isolate. Every haplotype is marked with a consecutive number and its size indicates the number of sequences included on it. Each node represents a mutational step between two haplotypes.

### 3.3. Multilocus genotyping

To determine whether markers with a higher resolution could distinguish host-usage patterns for the “rodent parasite models” *E. falciformis* and *E. vermiformis*, we designed a multilocus sequence typing approach. 35 markers targeting exons in the nuclear genome (supplementary data S4) and 5 regions of the apicoplast genome were amplified for 19 samples from *Apodemus* spp. hosts, 12 samples from house mice and corresponding regions from the reference genome of *E. falciformis* and *E. vermiformis* were included.

A multivariate analysis identified three clusters of isolates for the nuclear markers: one group included the laboratory isolate of *E. vermiformis*, another the isolate of *E. falciformis* and a third group only contained *Eimeria* isolates from *Apodemus agrarius* (Fig. 4B). This result was corroborated by phylogenetic analysis of SNPs (2019 informative alignment columns). We excluded prevalent indels from this analysis. Indels in protein coding genes (all “in-frame” with a length divisible by three) correspond to homopolymeric amino acid repeats (HAARs) and are expected in protein coding genes of Eimeria *spp.* (Heitlinger et al., 2014, Reid et al. 2014) (supplementary data S8). Three clades were recovered in this tree (Fig. 4A): The laboratory isolate of *E. vermiformis* from *Mus musculus* was indistinguishable from a field isolate from *Apodemus flavicollis*. Other isolates from house mouse and *A. flavicollis* and *A. sylvaticus* clustered in an unresovled internal relationship with house mouse *E. vermiformis* isolates (Group II). We note that four of five sequences for *E. vermiformis* from house mice were amplified from ileum tissue, the primary location of infection with this species (in contrast, *E. falciformis* infects primarily the caecum; Jarquín-Díaz et al 2019). A second clade recovered by nuclear multilocus analysis contained *E. falciformis* from house mice. This clade showed a well-supported substructure in which 7 house mouse field isolates grouped with the laboratory isolate BayerHaberkorn1970 but were separated from 4 *Eimeria* isolates from *A. flavicollis* (Group III).

**Figure 4.**
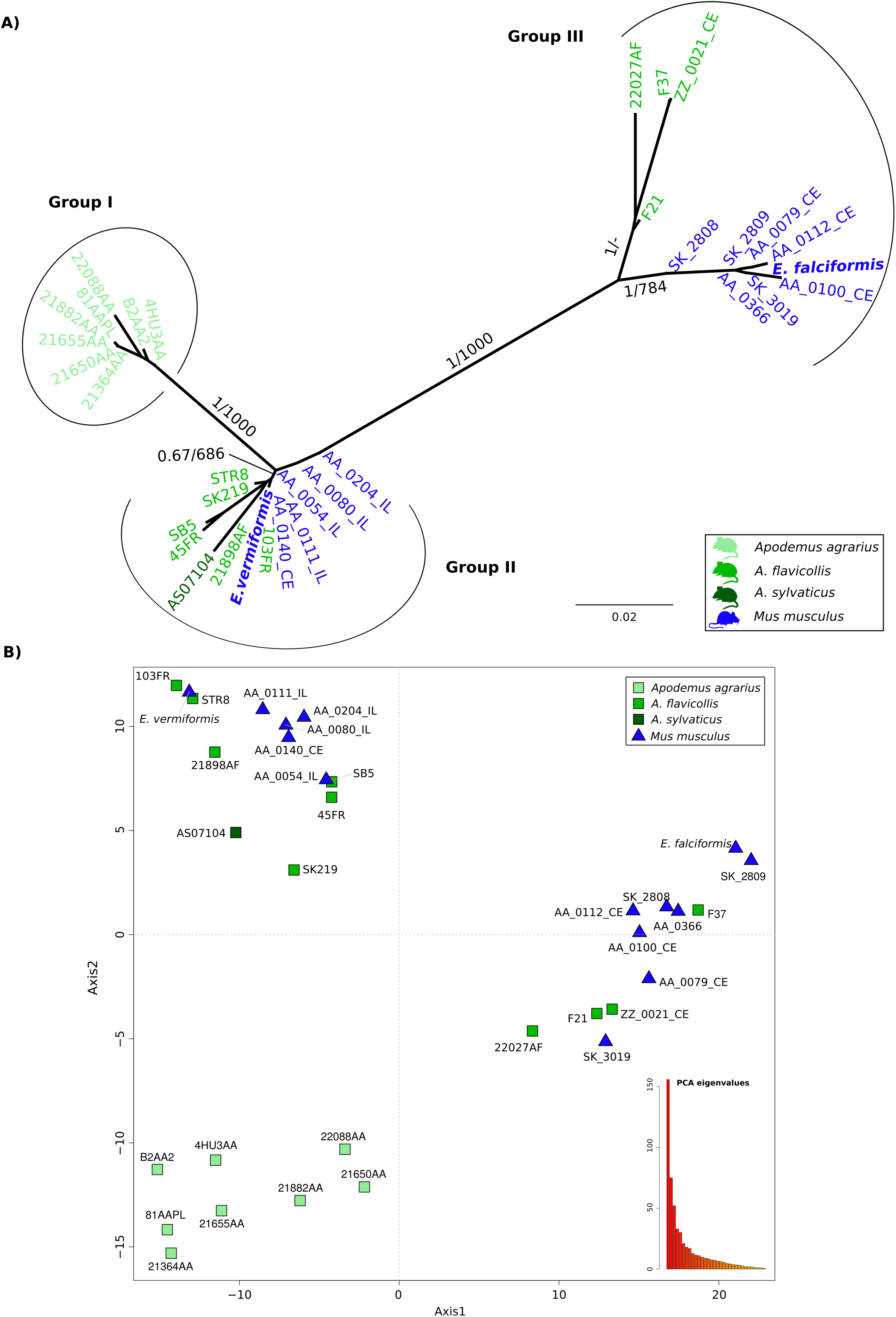
Nuclear multilocus genotyping of *Eimeria* isolates from *Mus musculus* and *Apodemus*. A) The phylogenetic tree was estimated with a multi-marker dataset formed with 35 nuclear markers from 31 *Eimeria* isolates derived from wild *Mus musculus* and three species of *Apodemus* (*A. agrarius, A. sylvaticus, A. flavicollis*). *Eimeria falciformis* and *E. vermiformis* sequences were included as reference. The scale bar represents sequence divergence. Colour represent the host of origin for the isolates. Bootstrap support values and Bayesian posterior probabilities are shown on branches. B) Principal component analysis based on single nucleotide polymorphisms (SNPs) from the same *Eimeria* isolates. Samples form three clusters. Shape indicate the genus of host and color the species. Eigenvalues of the dimensions are shown in an insert to visualize the proportion of variance explained by the axes.

Analyses based on apicoplast markers (both multivariate clustering and phylogenetic analyses; Fig. 5) identified similar groups: a well separated cluster with isolates from *A. agrarius*, a cluster containing *E. vermiformis* and another containing *E. falciformis* isolates. Some differences between the apicoplast and nuclear markers were obvious, though. *Eimeria* isolates from *M. musculus* (AA_0054_IL, AA_0080_IL, AA_0111_IL and AA_0112_CE) were less similar to the *E. vermiformis* group, leading to a multivariate clustering between the *E. falciformis* and *E. vermiformis* groups (Fig. 5B). This was recovered in a phylogenetic tree as isolates appeared at the end of a long branch in the *E. vermiformis* group (Fig. 5A). In an analysis of apicoplast markers, the *E. falciformis* isolates from *Mus* were not differentiated from those from *A. flavicollis*. Inspection of phylogenetic trees for individual markers (supplementary data S10) highlighted problems with the apicoplast dataset: samples that had been previously reported as co-infected with *E. ferrisi* (AA_0080_IL, AA_0111_IL and AA_0112_CE), showed an aberrant clustering for different markers. Samples AA_0080_IL and AA_0111_IL clustered in the group of *E. falciformis* with Ap12, while AA_0112_CE clustered with Ap5, in disagreement with the consensus species trees for other markers. We conclude that for these samples *E. ferrisi* or even *E. falciformis* apicoplast sequences were likely amplified and recovered as the majority sequence.

**Figure 5.**
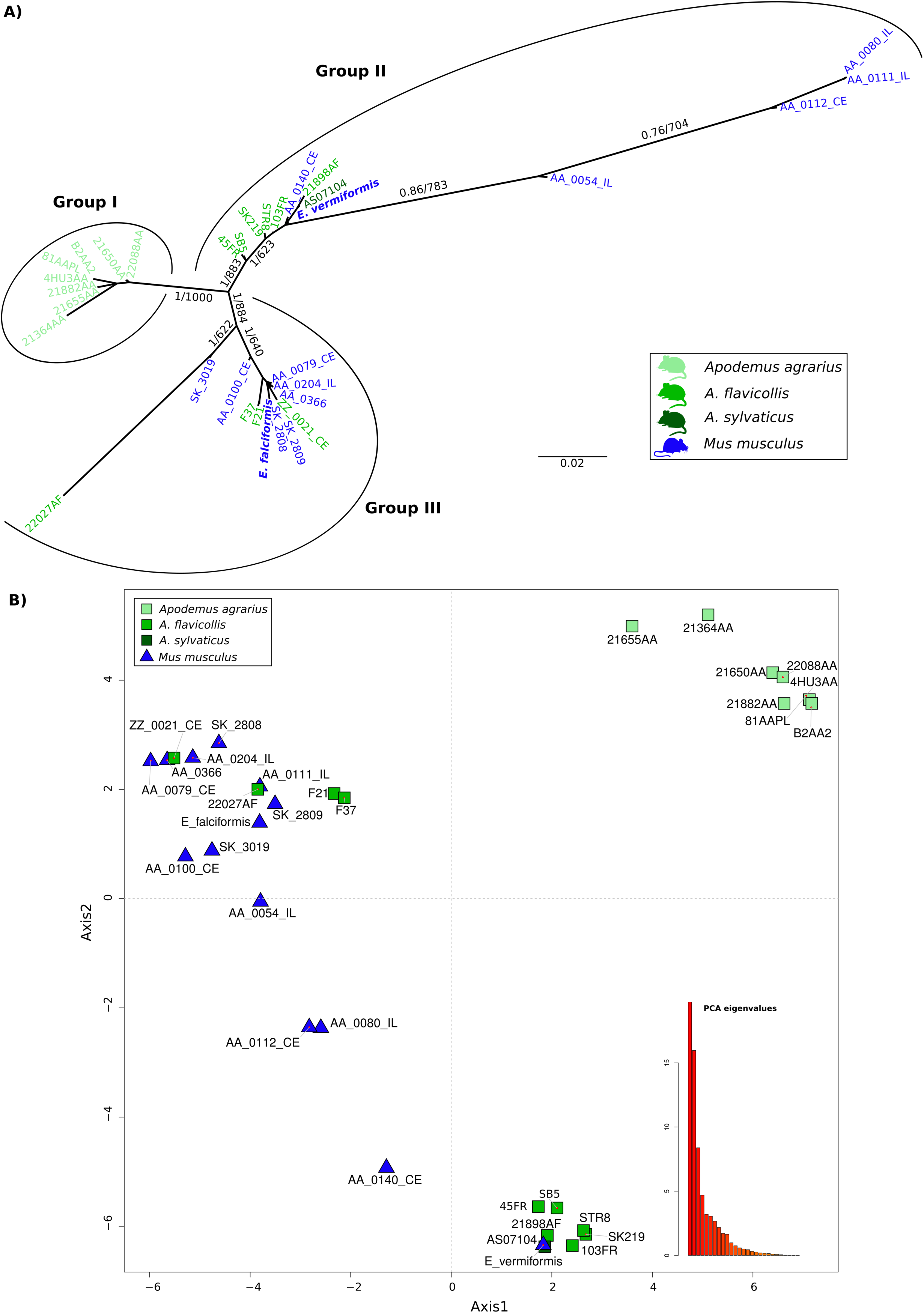
Apicoplast multilocus genotyping of *Eimeria* isolates from *Mus musculus* and *Apodemus*. A) The phylogenetic tree was estimated with a multi-marker dataset formed with 35 nuclear markers from 31 *Eimeria* wild isolates derived from *Mus musculus* and three species of *Apodemus* (*A. agrarius, A. sylvaticus, A. flavicollis*). *Eimeria falciformis* and *E. vermiformis* sequences were included as reference. The scale bar represent the sequence divergence. Colour represent the host of origin for the isolates. Bootstrap support values and Bayesian posterior probabilities are shown on branches. B) Principal component analysis based on single nucleotide polymorphisms (SNPs) from the same *Eimeria* isolates. Samples form three clusters based on the similarities for all the SNPs. Shape indicate the genus of host and color the species. Eigenvalues of the dimensions are shown in an insert to visualize the proportion of variance explained by the axes.

## 4. Discussion

We studied whether host specificity of Coccidia can be assessed with currently used molecular markers, using the example of *Eimeria* species in house mice and related rodents. We found that commonly used phylogenetic markers, nu 18S rDNA and mt COI, are not sufficiently variable to differentiate parasite isolates that would be regarded as separate species based on host usage. The relatively rarely used marker ap ORF470 from the apicoplast genome seems to provide slightly better resolution. We developed a multilocus genotyping approach to show that *E. falciformis* from the house mouse can likely be distinguished from related isolates from other hosts based on nuclear markers. In contrast, even with this high-resolution approach *E. vermiformis* from house mice and isolates from other host species were found in a nested and unresolved cluster.

Phylogenies derived from each of the analysed markers (esp. 18S) confirmed the topology of rodent *Eimeria* species observed before at deeper nodes of the phylogeny (Kvičerová, Pakandl, and Hypša 2008; Ogedengbe et al. 2018; Zhao and Duszynski 2001b). At the tips of the phylogeny, 18S sequences of *E. falciformis* and *E. vermiformis* isolates clustered with isolates from hosts of different genera or even families (Fig. 2A). This result was expected to some extent, as phylogenetic analyses with 18S sequences usually fail to separate closely related parasites isolated from closely related hosts (Ogedengbe et al. 2018).

Previous studies described COI as a universal barcode variable enough to resolve relationships between coccidians, including *Eimeria* (Ogedengbe, Hanner, and Barta 2011; Ogedengbe et al. 2018). We therefore expected to differentiate our house mouse isolates from species found in other hosts using COI. Neither phylogenetic (Fig. 2B) nor haplotype inference (Fig. 3), however, supported differentiation of *E. falciformis* and *E. vermiformis* from some of the isolates described as *E. apionodes.* Many of the COI sequences were even identical for isolates from different hosts. Limited resolution of COI outside of metazoans has been reported before (Meyer and Paulay, 2005). Rodent hosts of *Eimeria*, in the families Muridae (*Mus, Rattus, Apodemus*) and Cricetidae (*Myodes, Microtus*), diverged around 25 Million years ago (Churakov et al. 2010; Steppan, Adkins, and Anderson 2004) and it seems possible that COI of Coccidia evolves at such slow rates that it fails to differentiate *Eimeria* species with similar divergence. We stress that for rodent Coccidia, COI should not be assumed to resolve *bona fide* species with different host usage.

The potential of the apicoplast marker ORF470 to distinguish rodent *Eimeria* species has been highlighted before (Ogedengbe et al. 2015; Zhao and Duszynski 2001b), but few studies have followed the recommendation to use this marker. Consequently, few database sequences are available. Phylogenetic analysis of these sequences (Fig. 2C) separates our three species clusters well and shows hints of internal structure separating *E. apionodes* derived from *A. flavicollis* from house mouse isolates. Our work increases the number of sequences available for ORF470 and supports its use as a marker for discrimination of *Eimeria* species.

To test host specificity for *E. falciformis, E. vermiformis* (from house mice) and *E. apionodes* (from *Apodemus* spp.), we established and used a multilocus sequence typing protocol. Our multilocus approach supports a differentiation of *E. falciformis* (infecting the house mouse; Eimer, 1870; Haberkorn, 1970) from *E. apionodes* (infecting *A. flavicolis;* Pellérdy, 1954). The same approach was unable to distinguish *M. musculus* derived *E. vermiformis* isolates from one “*E. apionodes”* isolate from *A. flavicollis* (Fig. 2, 4 and 5). This suggests a broad host range of genetically indistinguishable *Eimeria* isolates which have been assigned to paraphlyetic species *E. apionodes* and *E. vermiformis*.

Multilocus genotyping using apicoplast markers showed some discrepancies with the nuclear analysis. These discrepancies can be attributed to double infections previously discovered in those particular isolates (Jarquín-Díaz et al. 2019). Compared to the nuclear genome, the apicoplast genome is present in much higher copy numbers (Heitlinger et al. 2014). This, combined with more conserved primer binding sites, can lead to amplification of non-target sequences such as those of the prevalent *E. ferrisi* (Jarquín-Díaz et al., 2019) creating artificial “chimeric” isolates in case of double infections.

We use our system also as a test case whether the commonly used markers (18S, COI) provide enough resolution to assess parasite specificity. We conclude that unresolved genetic clusters and monomorphic haplotypes currently identified via 18S and COI genotyping should not be assumed to indicate parasite species with generalist host usage. Novel nuclear markers are needed in addition to ORF470 to analyse host species specificity of rodent *Eimeria.* Care must be taken to avoid potential artifacts introduced by double infection and mixed amplification.

Whether other *Eimeria* species from different rodent hosts are indeed phylogentically distinguishable species (or whether genetically differentiable clusters show different host usage) is still an open question. This question needs to be addressed more broadly with markers providing higher resolution than 18S or COI.

## Acknowledgements

We thank Jaroslav Piálek and his team (Institute of Vertebrate Biology, AS CR, Brno, Department of Population Biology in Studenec) for help with catching and genotyping of house mice. Deborah Dymke and Julia Murata assisted the processing of samples in the Heitlinger group. We thank Susan Mbedi and Sarah Sparmann from the Berlin Center for Genomics in Biodiversity Research (BeGenDiv) for their technical guidance during the library preparation and Daniel P. Benesh for language revision and proofreading of the manuscript.

## Data Accessibility

- DNA sequences: nu 18S rDNA [MH751925-MH752036, MK246860-MK246868 and MK625202-MK625210]; mt COI [MH777467-MH777593, MH755302-MH755324, MK257106-MK257114 and MK631866-MK631868] and ap ORF470 [MH755325-MH755450, MK257115-MK257125 and MK631869-MK631884]).
- All sequencing raw data can be accessed through the BioProject PRJNA548431 in the NCBI Short Read Archive (SRA).
- The code for the pipeline used in the multi locus type analysis is available on github at https://github.com/VictorHJD/AA_Eimeria_Genotyping.

## Author Contributions

VHJD and EH designed the project and obtained funding. VHJD, AM, TRS, JJ, KB, ST and JK obtained data, VHJD, AB and EH designed the analysis, VHJD, AB and EH performed the analysis. VHJD, EH and JK interpreted the results. VHJD and EH wrote the manuscript with contributions from all other authors. EH supervised the project.

## Supplementary Information

**Supplementary data S1. Origin of samples.**

**Supplementary data S2. Accession number from reference sequences used for phylogenetic analysis.**

**Supplementary data S3. Primer pair list for multimarker genotyping of *Eimeria*.**

**Supplementary data S4. Position of primers for genotyping in *Eimeria falciformis* nuclear genome.** Primer pairs designed to amplify protein coding exons from *E. falciformis* were located across the nuclear genome of *E. falciformis*. Primer pairs are marked with yellow marks and names are highlighted in blue on the right side of the contigs. When more than one primer pair was located in the same contig, names are ordered from left to right.

**Supplementary data S5. Position of primers for genotyping in *Eimeria falciformis* apicoplast genome.** Primer pairs designed to amplify different regions from the apicoplast of *Eimeria* spp. were located across the circular annotated representation of this genome. Primer pairs are marked with green marks and names are highlighted.

**Supplementary data S6. Amplified Sequence Variants (ASVs) recovered for each sample in multilocus amplification.** Each heatmap represents one marker, samples are listed in rows, different ASVs in columns. The number of sequence reads recovered per ASV and sample is log10 transformed and displayed in a colour shading. Pages 1-10: Apicoplast regions, 11 – 75: nuclear makers.

**Supplementary data S7. Concatenated phylogenetic inference with nuclear (18S rDNA), mitochondrial (COI) and apicoplast (ORF470) sequences.**

**Supplementary data S8. Estimation of informative sites within the multilocus alignment of nuclear genes.**

**Supplementary data S9. Estimation of informative sites within the multilocus alignment of apicoplast regions.**

**Supplementary data S10. Individual phylogenetic trees inferred from apicoplast markers.** Trees inferred for the apicoplast markes in an analysis of individual regions. A) Ap7, B) Ap10, C) Ap11, D) Ap12 and E) Ap 5.

