## Supplementary figures and images for "Generalist *Eimeria* species in rodents: multilocus analyses indicate inadequate resolution of established markers"

### Supplementary data S4

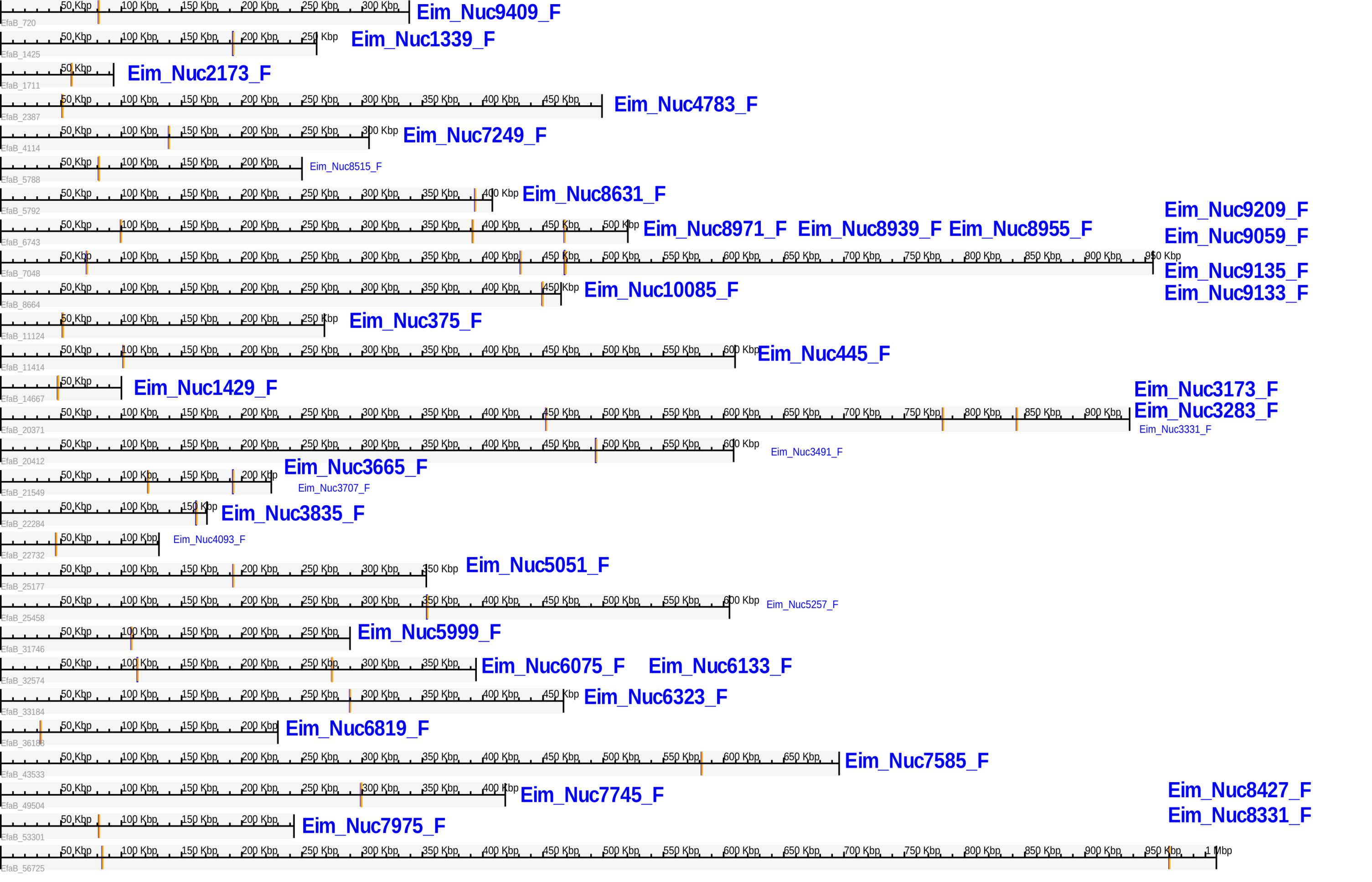

### Supplementary data S5

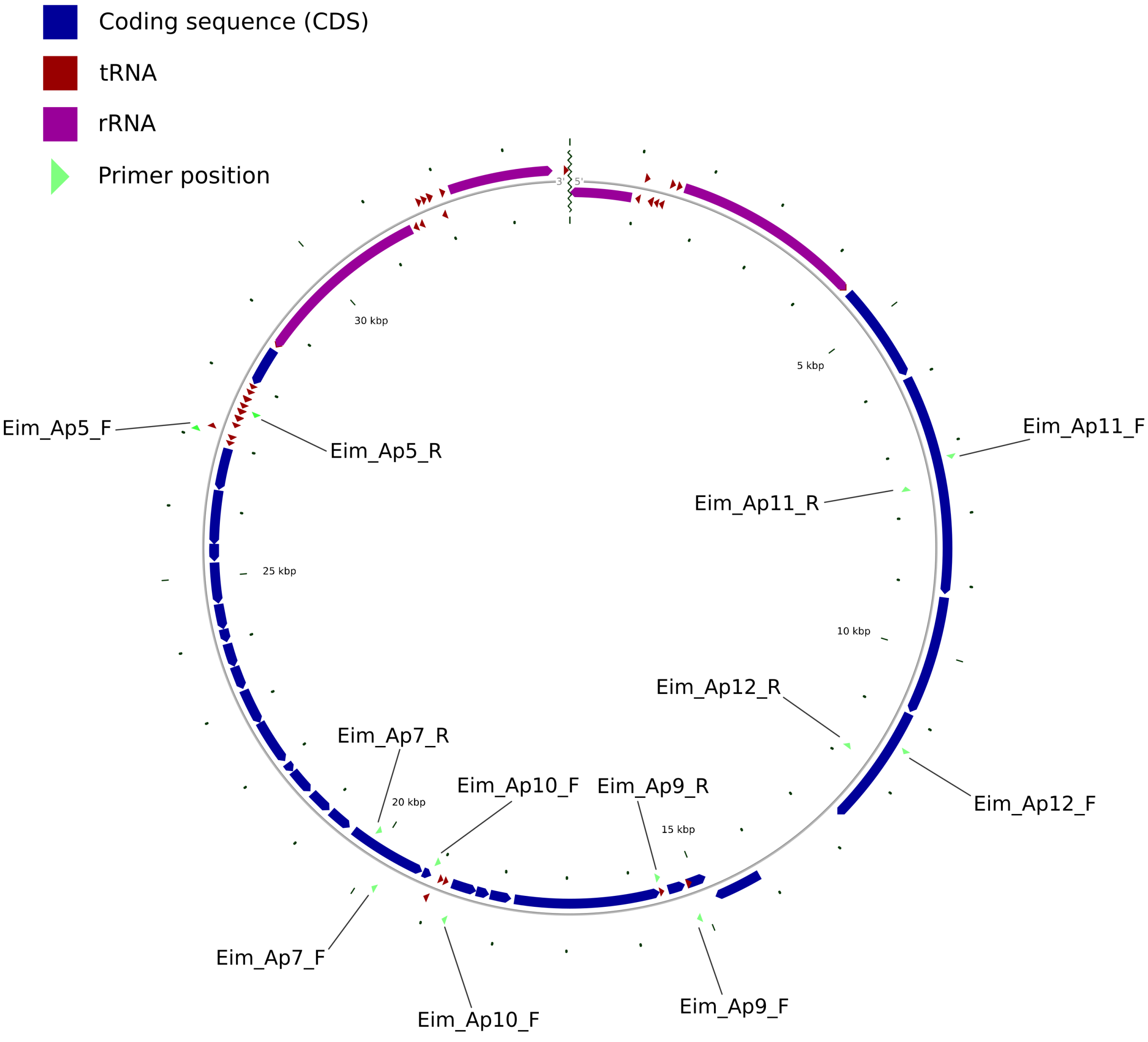

### Supplementary data S7

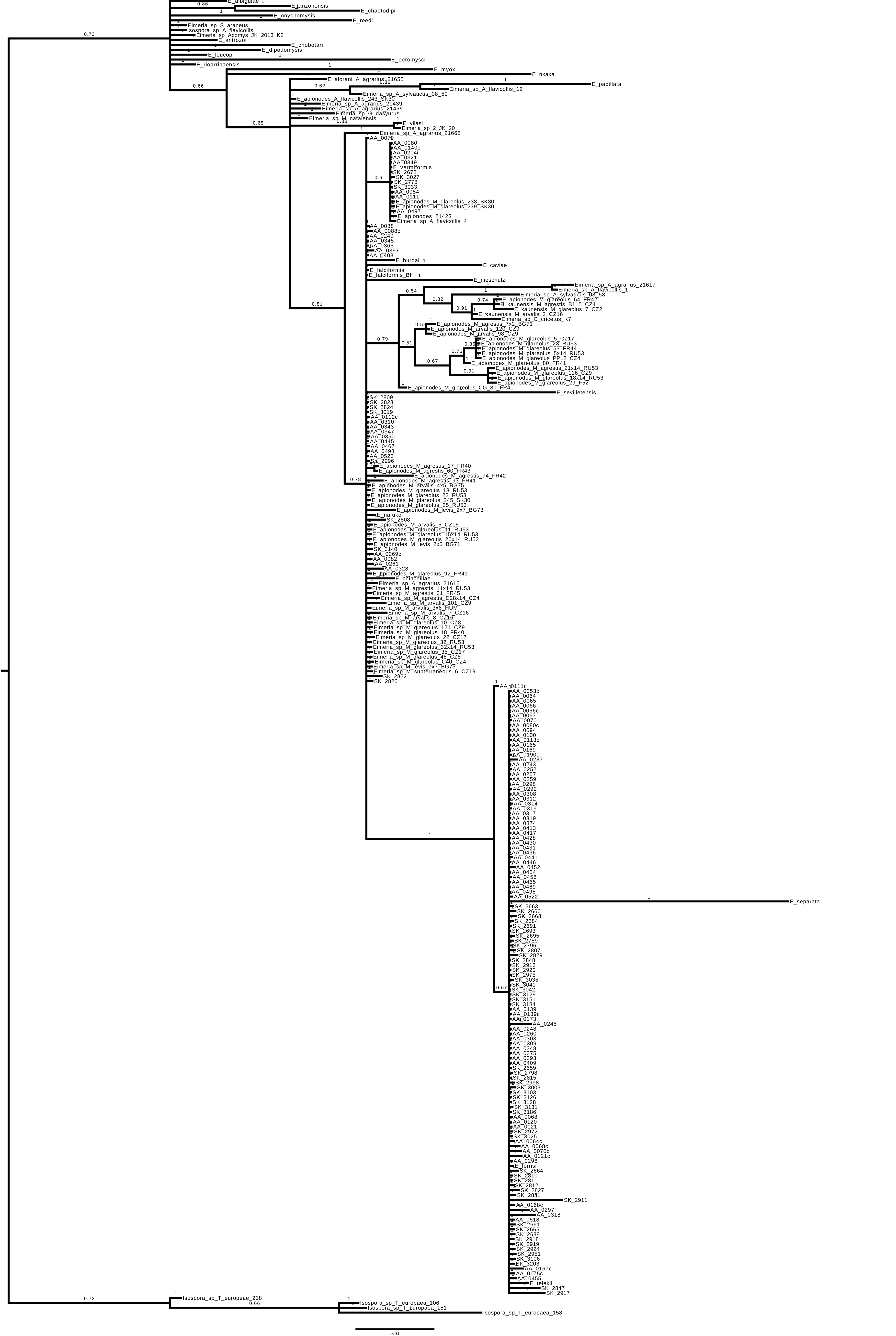

### Supplementary data S10

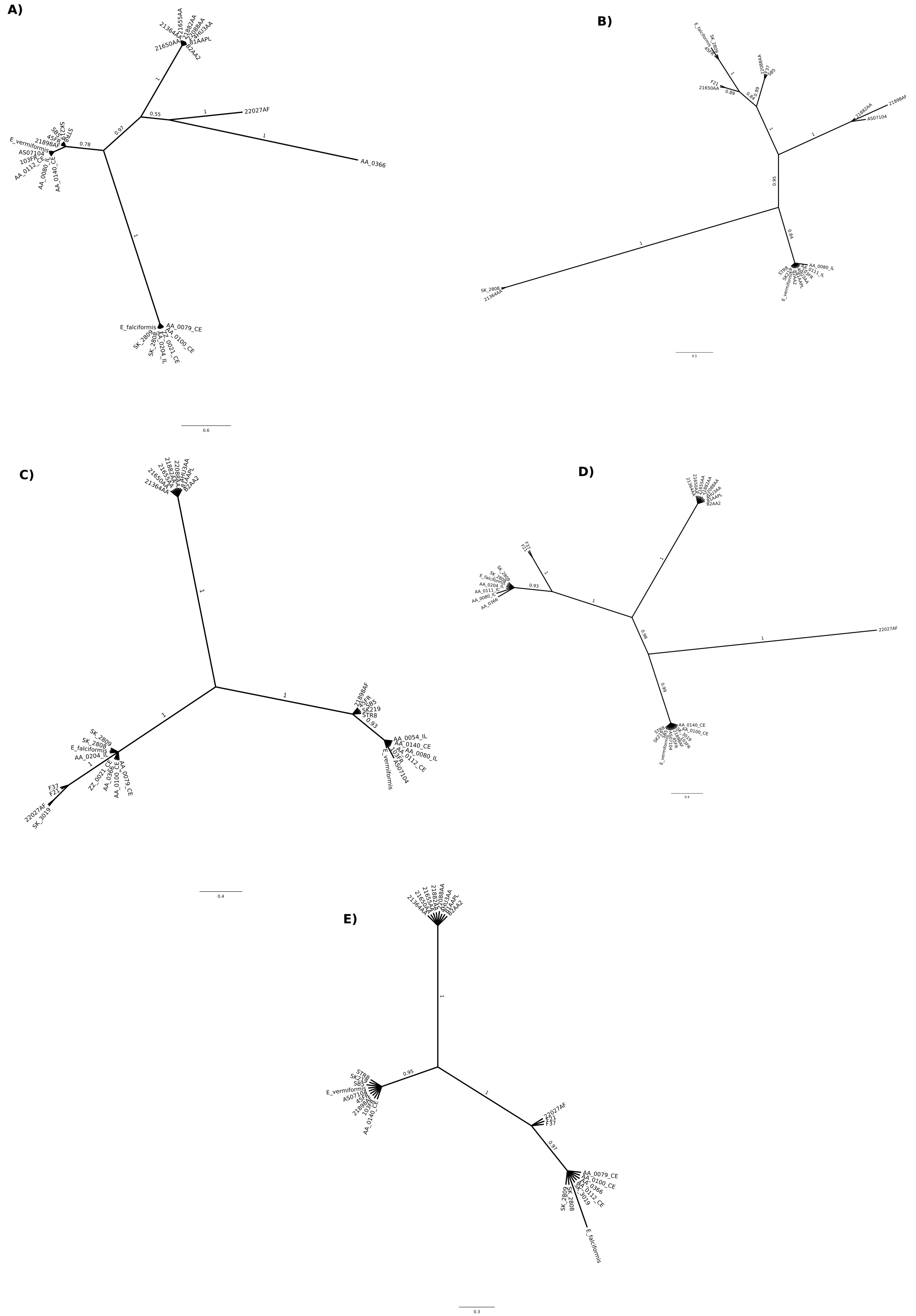
