## Supplementary data S6 for "Generalist *Eimeria* species in rodents: multilocus analyses indicate inadequate resolution of established markers"

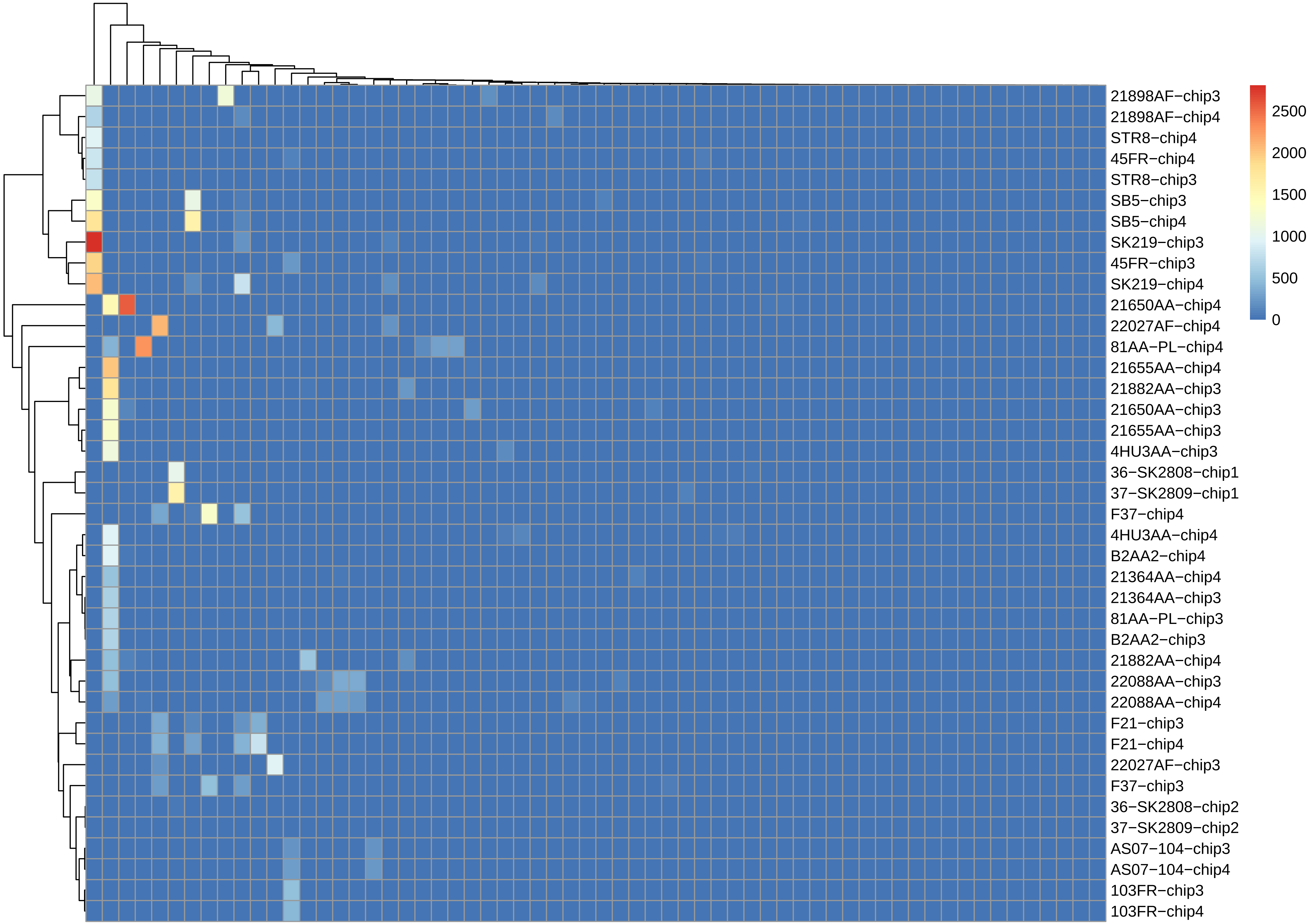
21898AF−chip3
21898AF−chip4
STR8−chip4
45FR−chip4
STR8−chip3
SB5−chip3
SB5−chip4
SK219−chip3
45FR−chip3
SK219−chip4
21650AA−chip4
22027AF−chip4
81AA−PL−chip4
21655AA−chip4
21882AA−chip3
21650AA−chip3
21655AA−chip3
4HU3AA−chip3
36−SK2808−chip1
37−SK2809−chip1
F37−chip4
4HU3AA−chip4
B2AA2−chip4
21364AA−chip4
21364AA−chip3
81AA−PL−chip3
B2AA2−chip3
21882AA−chip4
22088AA−chip3
22088AA−chip4
F21−chip3
F21−chip4
22027AF−chip3
F37−chip3
36−SK2808−chip2
37−SK2809−chip2
AS07−104−chip3
AS07−104−chip4
103FR−chip3
103FR−chip4
2500
2000
1500
1000
500
0

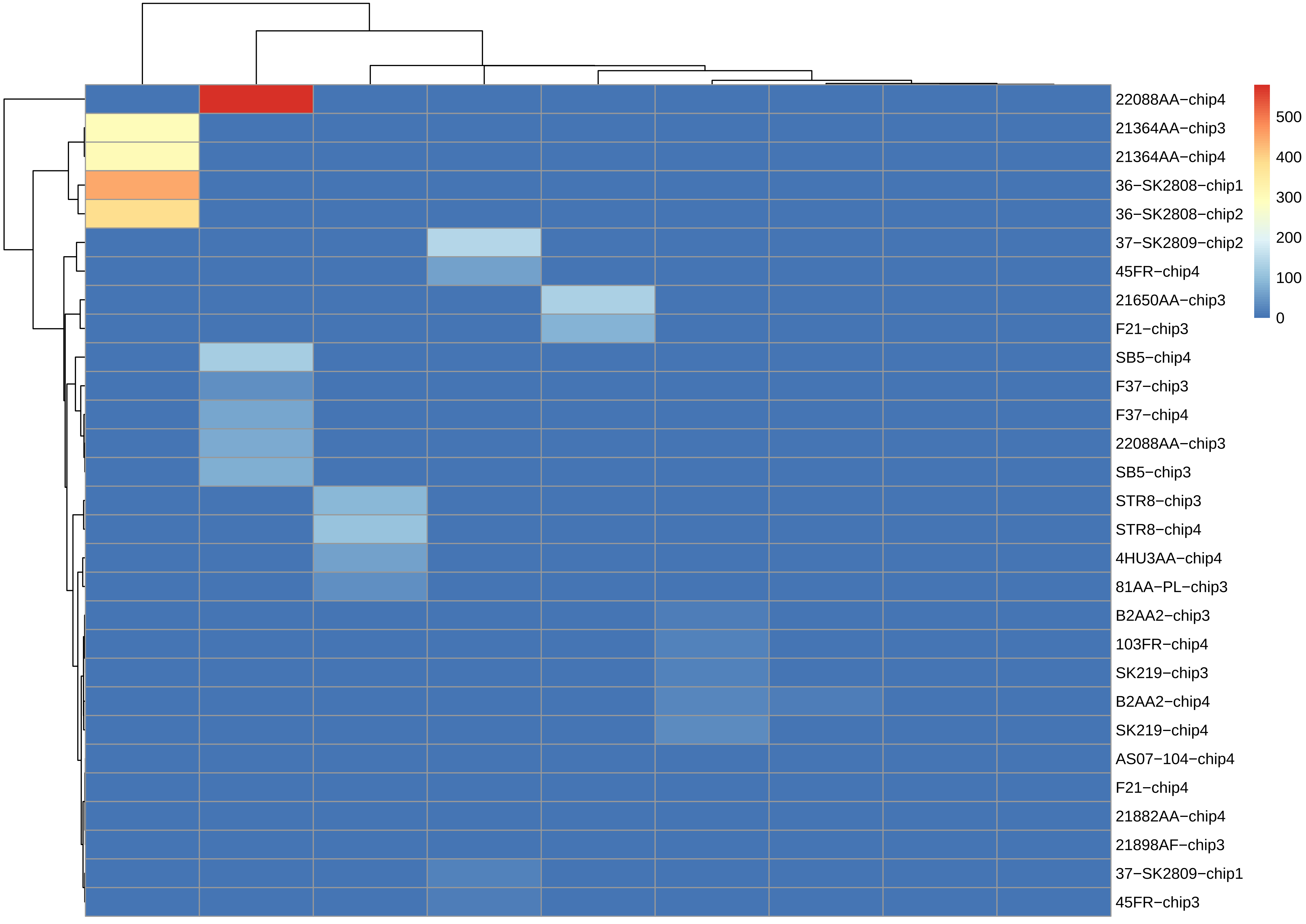

22088AA−chip4
21364AA−chip3
21364AA−chip4
36−SK2808−chip1
36−SK2808−chip2
37−SK2809−chip2
45FR−chip4
21650AA−chip3
F21−chip3
SB5−chip4
F37−chip3
F37−chip4
22088AA−chip3
SB5−chip3
STR8−chip3
STR8−chip4
4HU3AA−chip4
81AA−PL−chip3
B2AA2−chip3
103FR−chip4
SK219−chip3
B2AA2−chip4
SK219−chip4
AS07−104−chip4
F21−chip4
21882AA−chip4
21898AF−chip3
37−SK2809−chip1
45FR−chip3
500
400
300
200
100
0

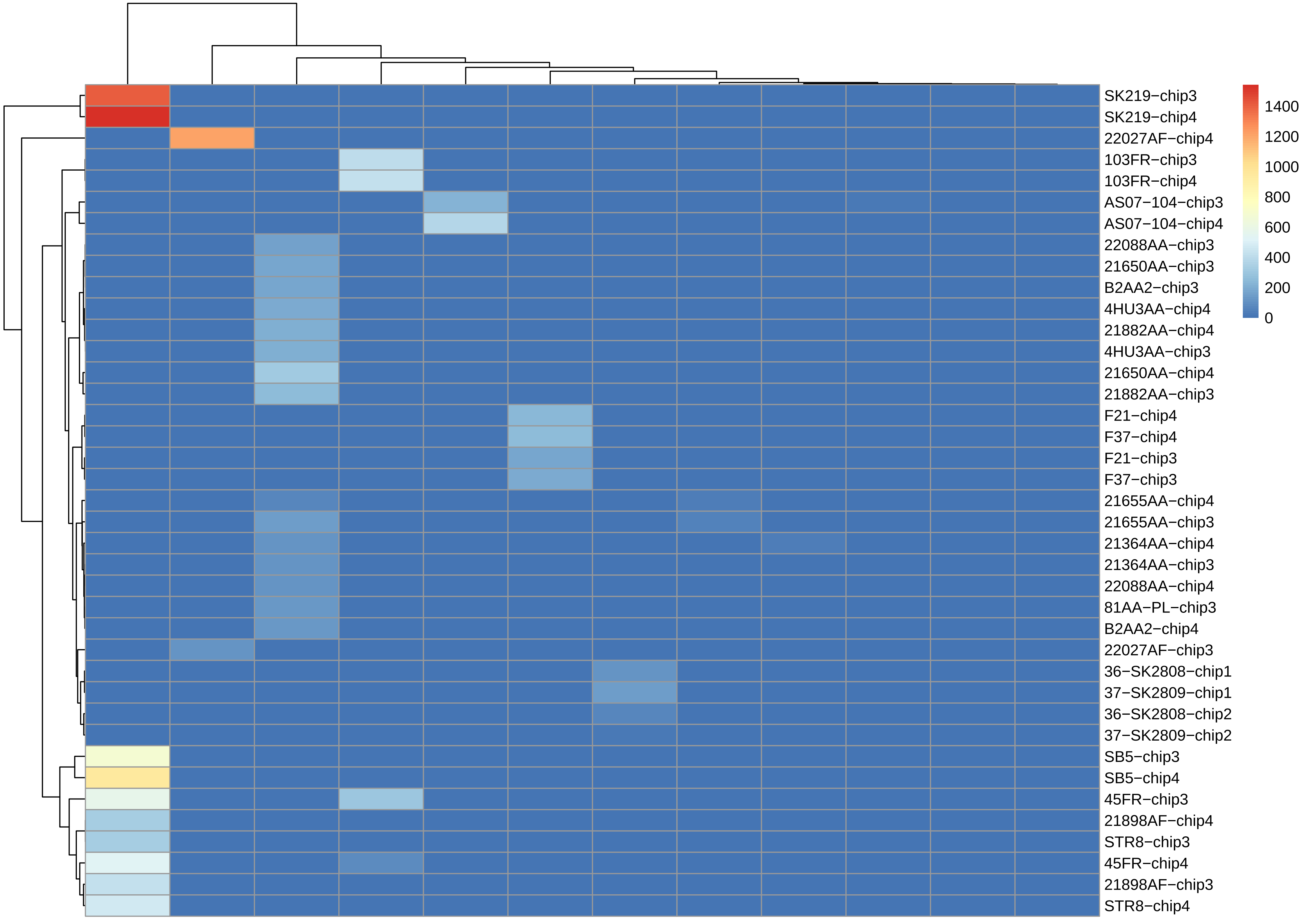

SK219−chip3
SK219−chip4
22027AF−chip4
103FR−chip3
103FR−chip4
AS07−104−chip3
AS07−104−chip4
22088AA−chip3
21650AA−chip3
B2AA2−chip3
4HU3AA−chip4
21882AA−chip4
4HU3AA−chip3
21650AA−chip4
21882AA−chip3
F21−chip4
F37−chip4
F21−chip3
F37−chip3
21655AA−chip4
21655AA−chip3
21364AA−chip4
21364AA−chip3
22088AA−chip4
81AA−PL−chip3
B2AA2−chip4
22027AF−chip3
36−SK2808−chip1
37−SK2809−chip1
36−SK2808−chip2
37−SK2809−chip2
SB5−chip3
SB5−chip4
45FR−chip3
21898AF−chip4
STR8−chip3
45FR−chip4
21898AF−chip3
STR8−chip4
1400
1200
1000
800
600
400
200
0

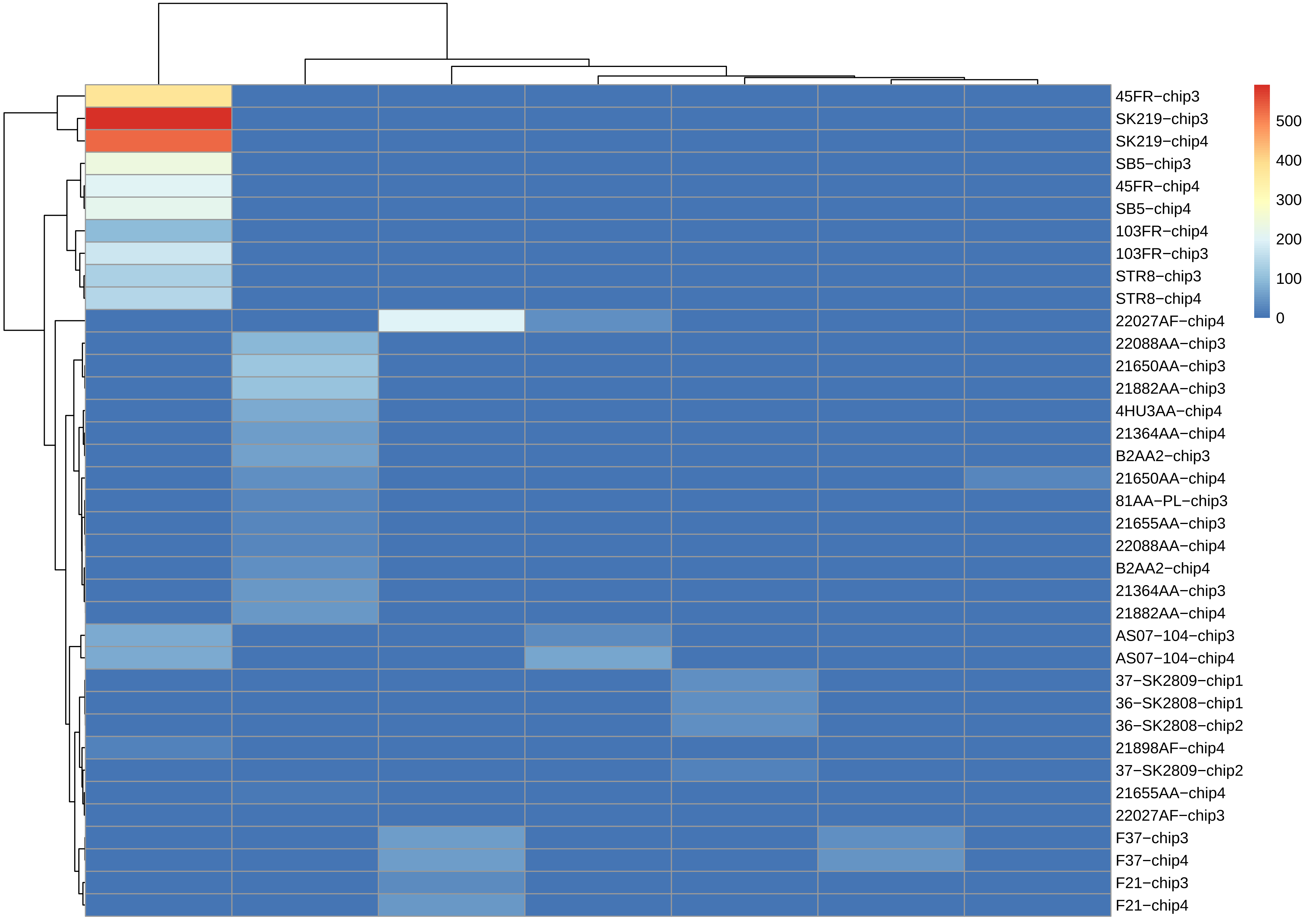
45FR−chip3
SK219−chip3
SK219−chip4
SB5−chip3
45FR−chip4
SB5−chip4
103FR−chip4
103FR−chip3
STR8−chip3
STR8−chip4
22027AF−chip4
22088AA−chip3
21650AA−chip3
21882AA−chip3
4HU3AA−chip4
21364AA−chip4
B2AA2−chip3
21650AA−chip4
81AA−PL−chip3
21655AA−chip3
22088AA−chip4
B2AA2−chip4
21364AA−chip3
21882AA−chip4
AS07−104−chip3
AS07−104−chip4
37−SK2809−chip1
36−SK2808−chip1
36−SK2808−chip2
21898AF−chip4
37−SK2809−chip2
21655AA−chip4
22027AF−chip3
F37−chip3
F37−chip4
F21−chip3
F21−chip4
500
400
300
200
100
0

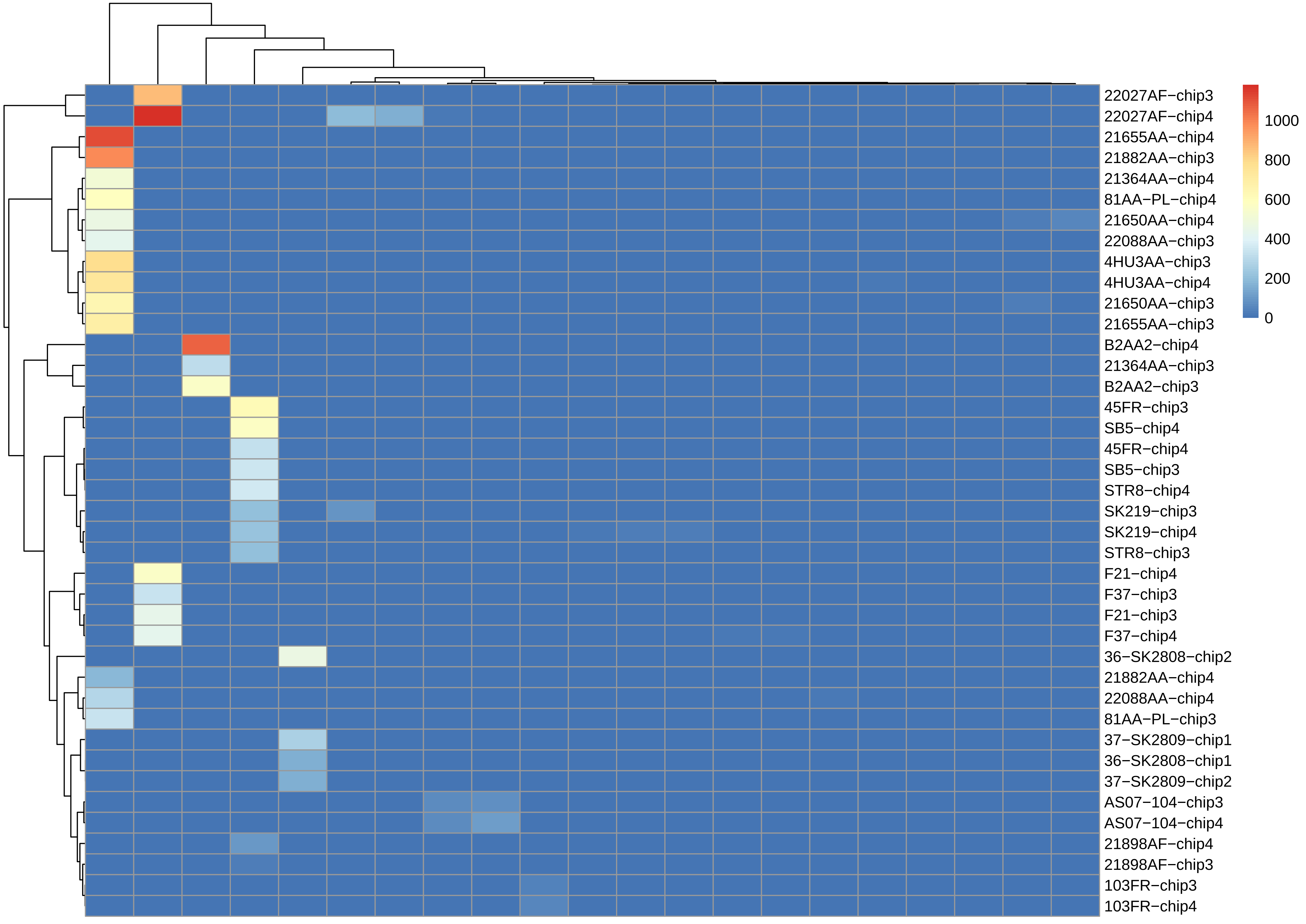

22027AF−chip3
22027AF−chip4
21655AA−chip4
21882AA−chip3
21364AA−chip4
81AA−PL−chip4
21650AA−chip4
22088AA−chip3
4HU3AA−chip3
4HU3AA−chip4
21650AA−chip3
21655AA−chip3
B2AA2−chip4
21364AA−chip3
B2AA2−chip3
45FR−chip3
SB5−chip4
45FR−chip4
SB5−chip3
STR8−chip4
SK219−chip3
SK219−chip4
STR8−chip3
F21−chip4
F37−chip3
F21−chip3
F37−chip4
36−SK2808−chip2
21882AA−chip4
22088AA−chip4
81AA−PL−chip3
37−SK2809−chip1
36−SK2808−chip1
37−SK2809−chip2
AS07−104−chip3
AS07−104−chip4
21898AF−chip4
21898AF−chip3
103FR−chip3
103FR−chip4
1000
800
600
400
200
0

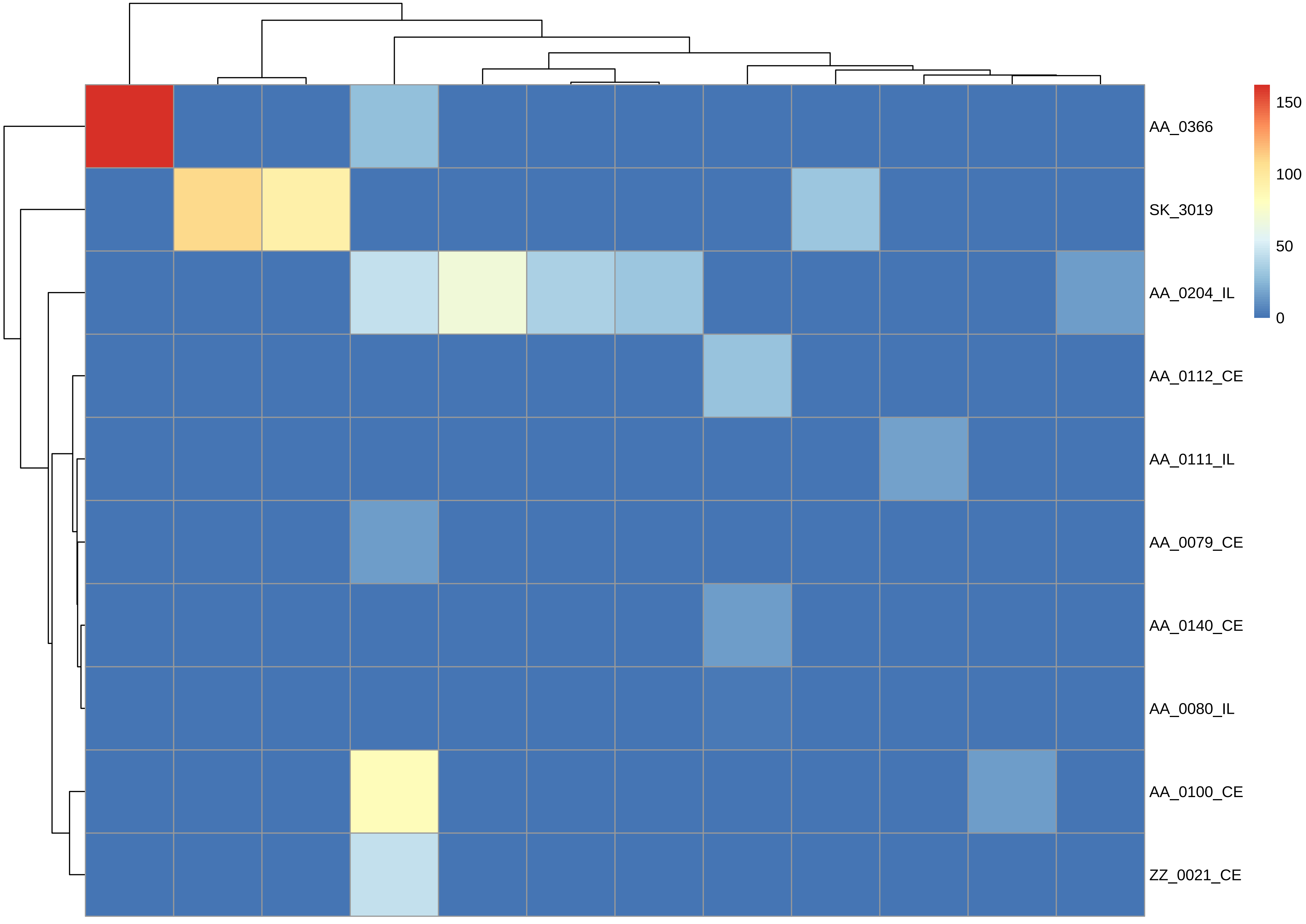

AA_0366
SK_3019
AA_0204_IL
AA_0112_CE
AA_0111_IL
AA_0079_CE
AA_0140_CE
AA_0080_IL
AA_0100_CE
ZZ_0021_CE
150
100
50
0

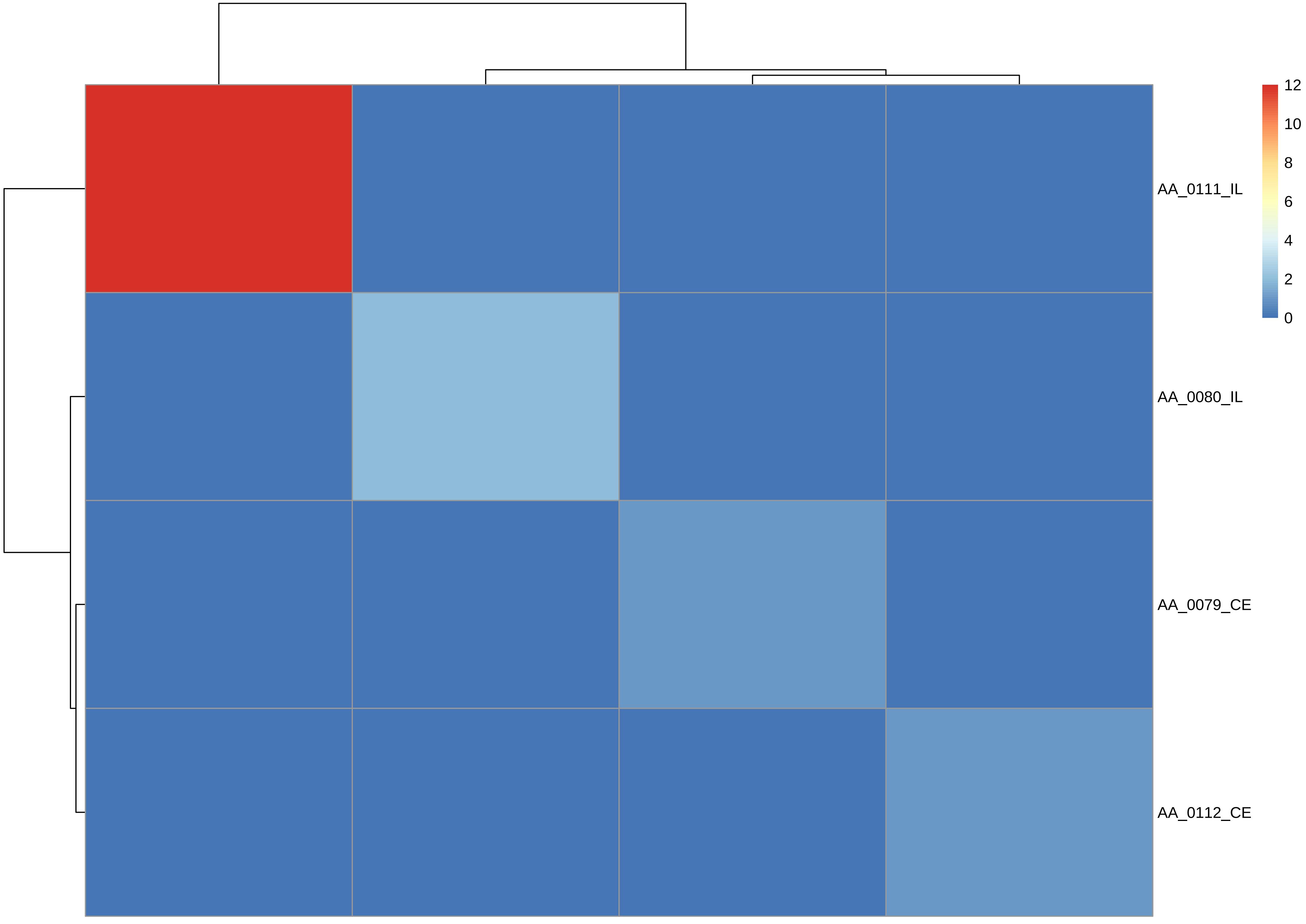

AA_0111_IL
AA_0080_IL
AA_0079_CE
AA_0112_CE
12
10
8
6
4
2
0

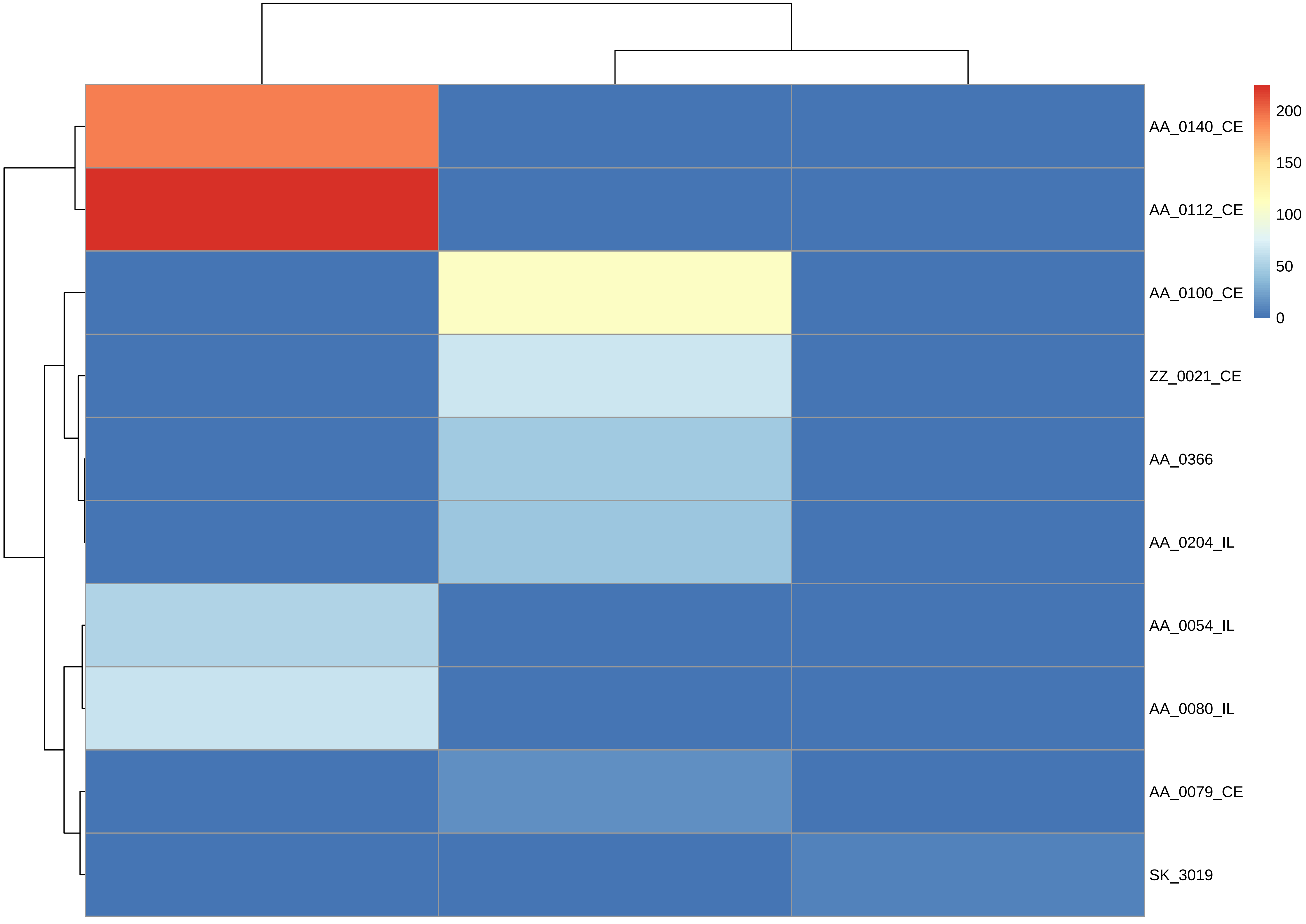
AA_0140_CE
AA_0112_CE
AA_0100_CE
ZZ_0021_CE
AA_0366
AA_0204_IL
AA_0054_IL
AA_0080_IL
AA_0079_CE
SK_3019
200
150
100
50
0

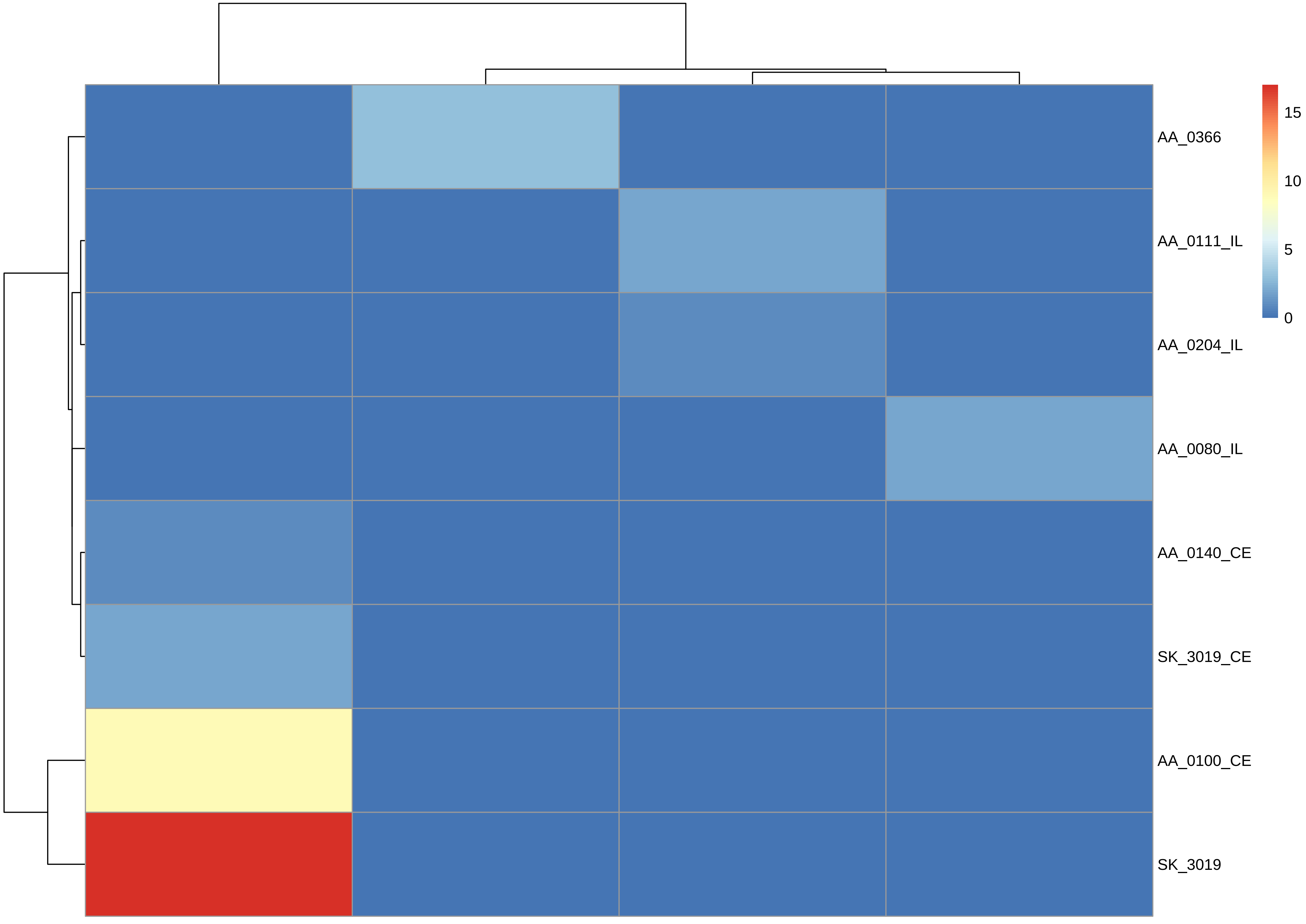

AA_0366
AA_0111_IL
AA_0204_IL
AA_0080_IL
AA_0140_CE
SK_3019_CE
AA_0100_CE
SK_3019
15
10
5
0

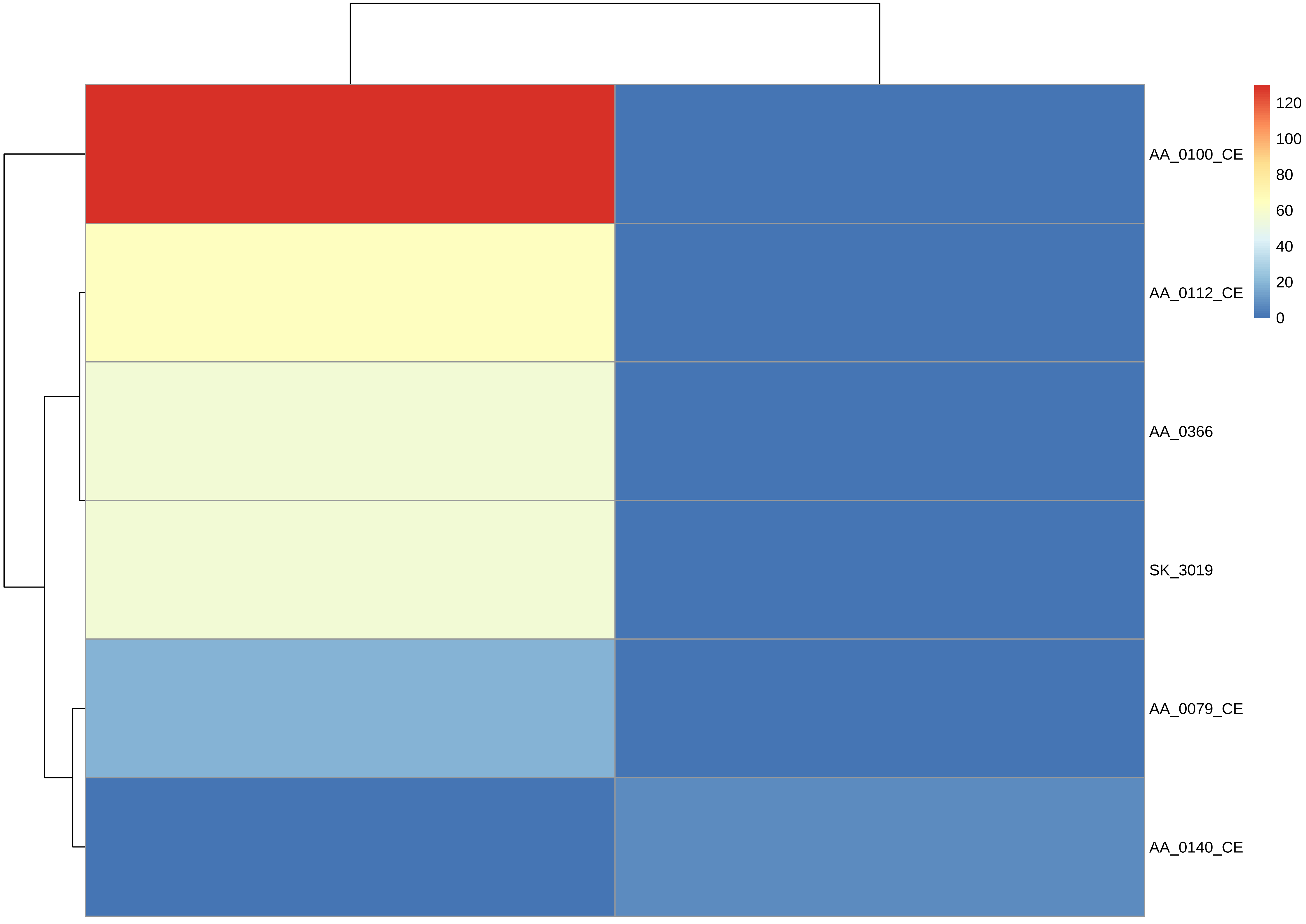

AA_0100_CE
AA_0112_CE
AA_0366
SK_3019
AA_0079_CE
AA_0140_CE
120
100
80
60
40
20
0

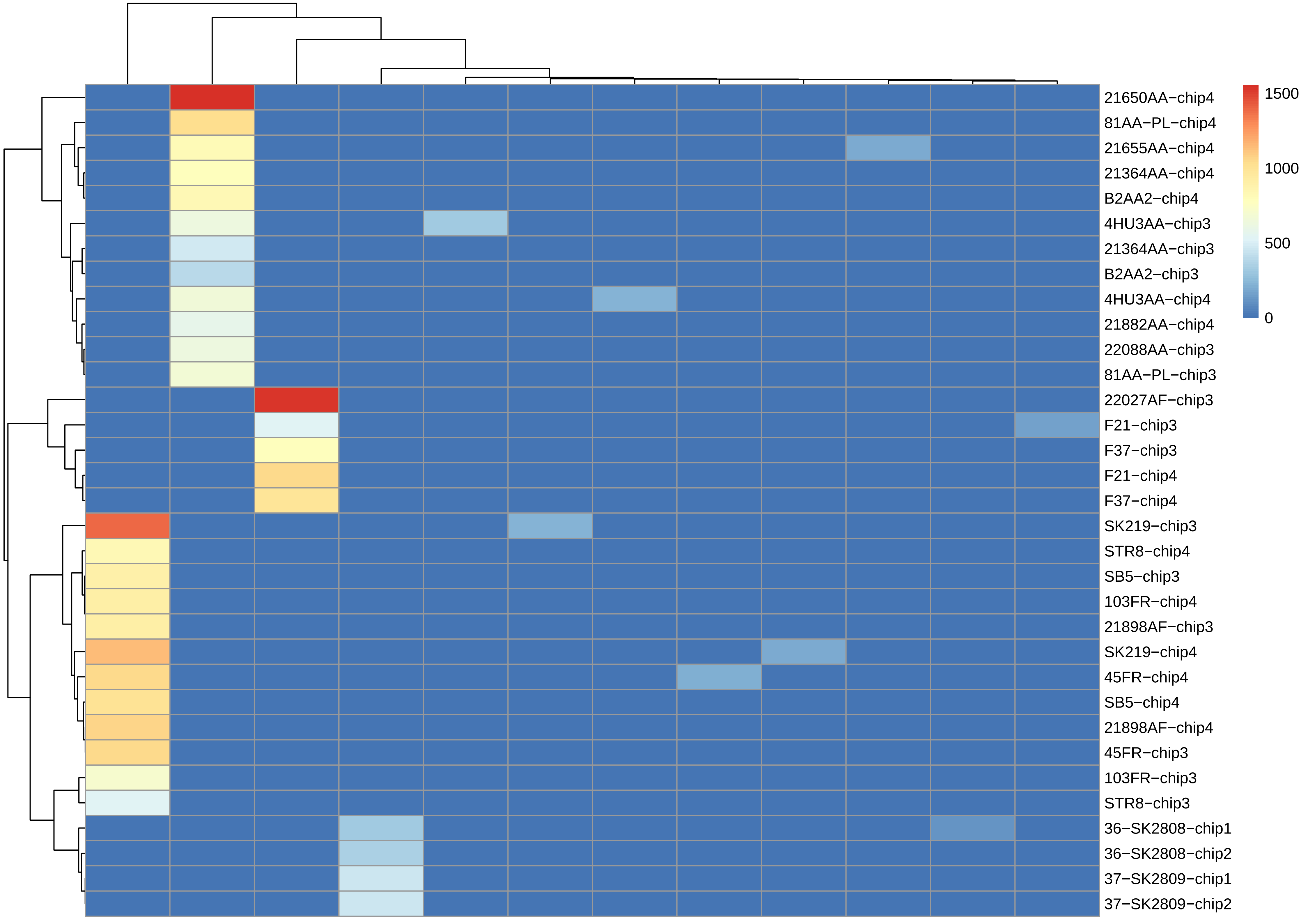

1500
1000
500
0
21650AA−chip4
81AA−PL−chip4
21655AA−chip4
21364AA−chip4
B2AA2−chip4
4HU3AA−chip3
21364AA−chip3
B2AA2−chip3
4HU3AA−chip4
21882AA−chip4
22088AA−chip3
81AA−PL−chip3
22027AF−chip3
F21−chip3
F37−chip3
F21−chip4
F37−chip4
SK219−chip3
STR8−chip4
SB5−chip3
103FR−chip4
21898AF−chip3
SK219−chip4
45FR−chip4
SB5−chip4
21898AF−chip4
45FR−chip3
103FR−chip3
STR8−chip3
36−SK2808−chip1
36−SK2808−chip2
37−SK2809−chip1
37−SK2809−chip2

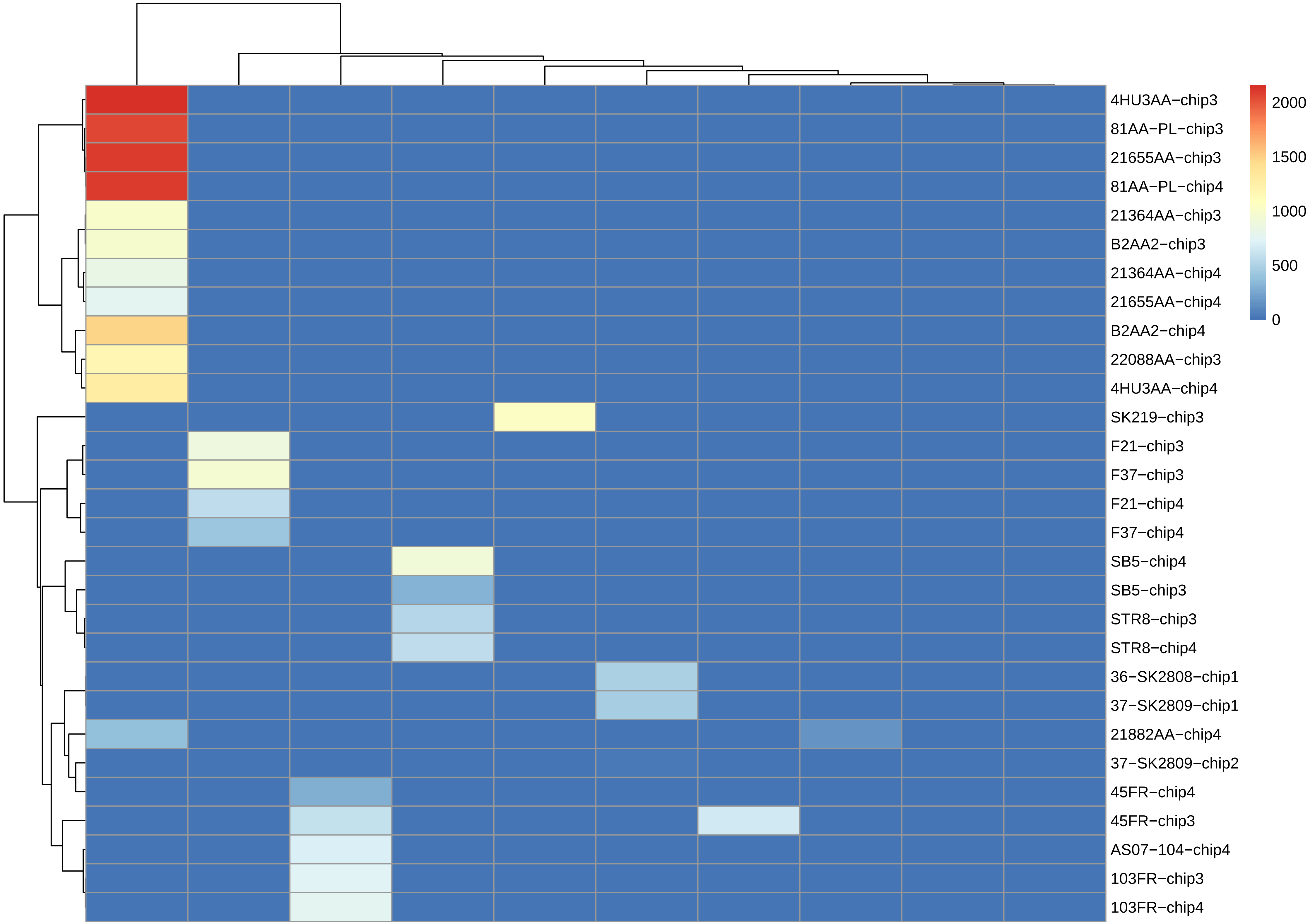
4HU3AA−chip3
81AA−PL−chip3
21655AA−chip3
81AA−PL−chip4
21364AA−chip3
B2AA2−chip3
21364AA−chip4
21655AA−chip4
B2AA2−chip4
22088AA−chip3
4HU3AA−chip4
SK219−chip3
F21−chip3
F37−chip3
F21−chip4
F37−chip4
SB5−chip4
SB5−chip3
STR8−chip3
STR8−chip4
36−SK2808−chip1
37−SK2809−chip1
21882AA−chip4
37−SK2809−chip2
45FR−chip4
45FR−chip3
AS07−104−chip4
103FR−chip3
103FR−chip4
2000
1500
1000
500
0

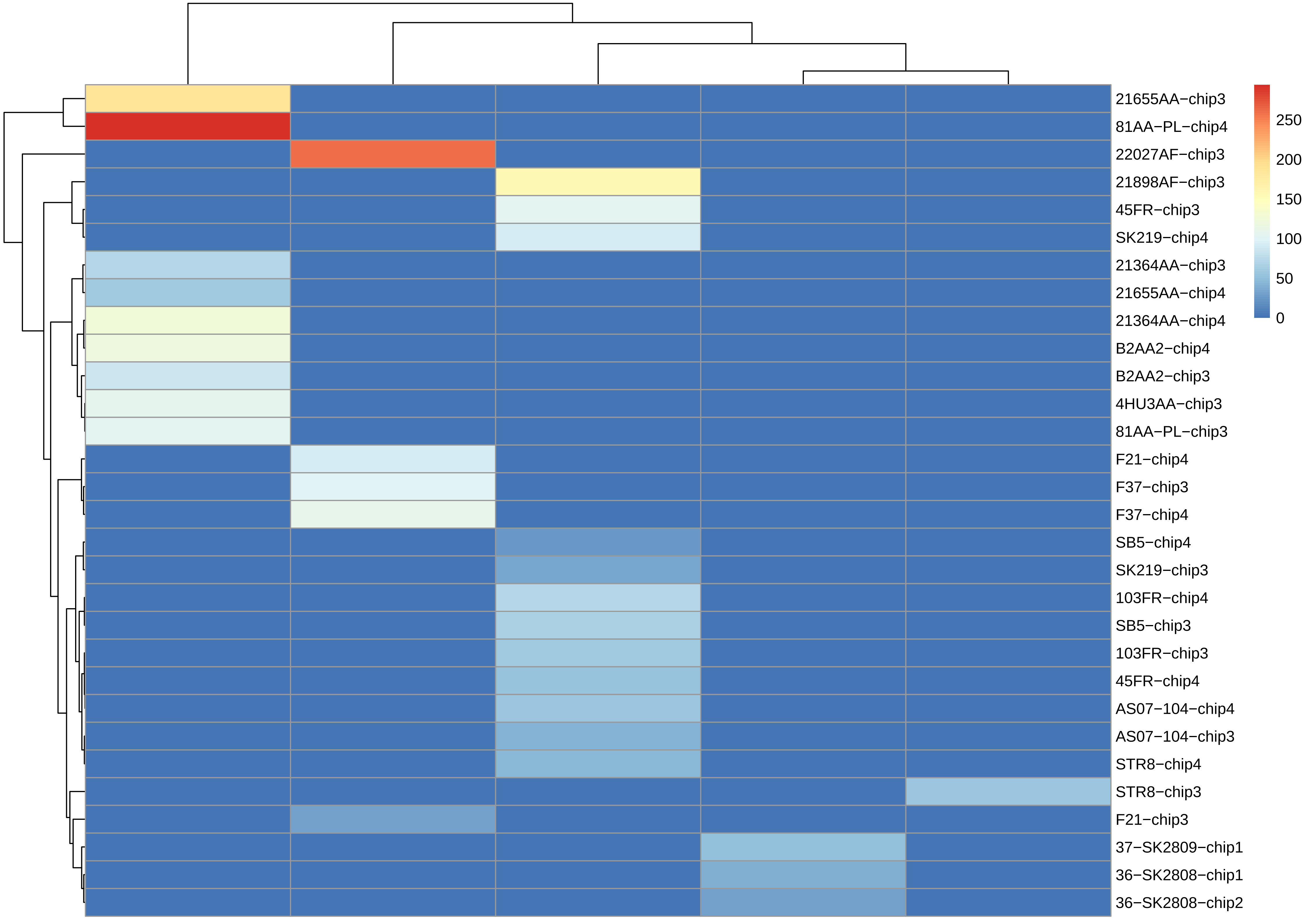
21655AA−chip3
81AA−PL−chip4
22027AF−chip3
21898AF−chip3
45FR−chip3
SK219−chip4
21364AA−chip3
21655AA−chip4
21364AA−chip4
B2AA2−chip4
B2AA2−chip3
4HU3AA−chip3
81AA−PL−chip3
F21−chip4
F37−chip3
F37−chip4
SB5−chip4
SK219−chip3
103FR−chip4
SB5−chip3
103FR−chip3
45FR−chip4
AS07−104−chip4
AS07−104−chip3
STR8−chip4
STR8−chip3
F21−chip3
37−SK2809−chip1
36−SK2808−chip1
36−SK2808−chip2
250
200
150
100
50
0

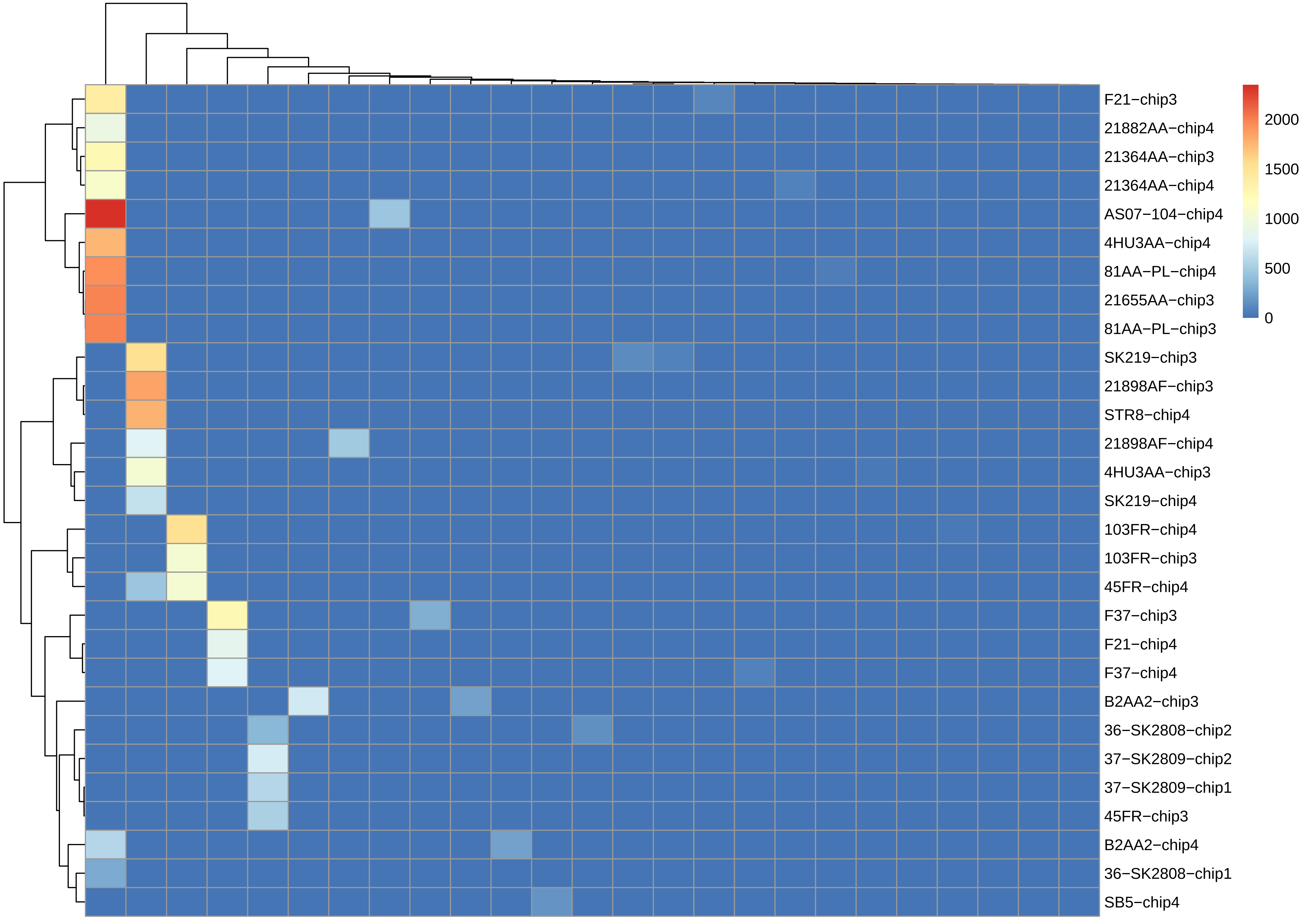

F21−chip3
21882AA−chip4
21364AA−chip3
21364AA−chip4
AS07−104−chip4
4HU3AA−chip4
81AA−PL−chip4
21655AA−chip3
81AA−PL−chip3
SK219−chip3
21898AF−chip3
STR8−chip4
21898AF−chip4
4HU3AA−chip3
SK219−chip4
103FR−chip4
103FR−chip3
45FR−chip4
F37−chip3
F21−chip4
F37−chip4
B2AA2−chip3
36−SK2808−chip2
37−SK2809−chip2
37−SK2809−chip1
45FR−chip3
B2AA2−chip4
36−SK2808−chip1
SB5−chip4
2000
1500
1000
500
0

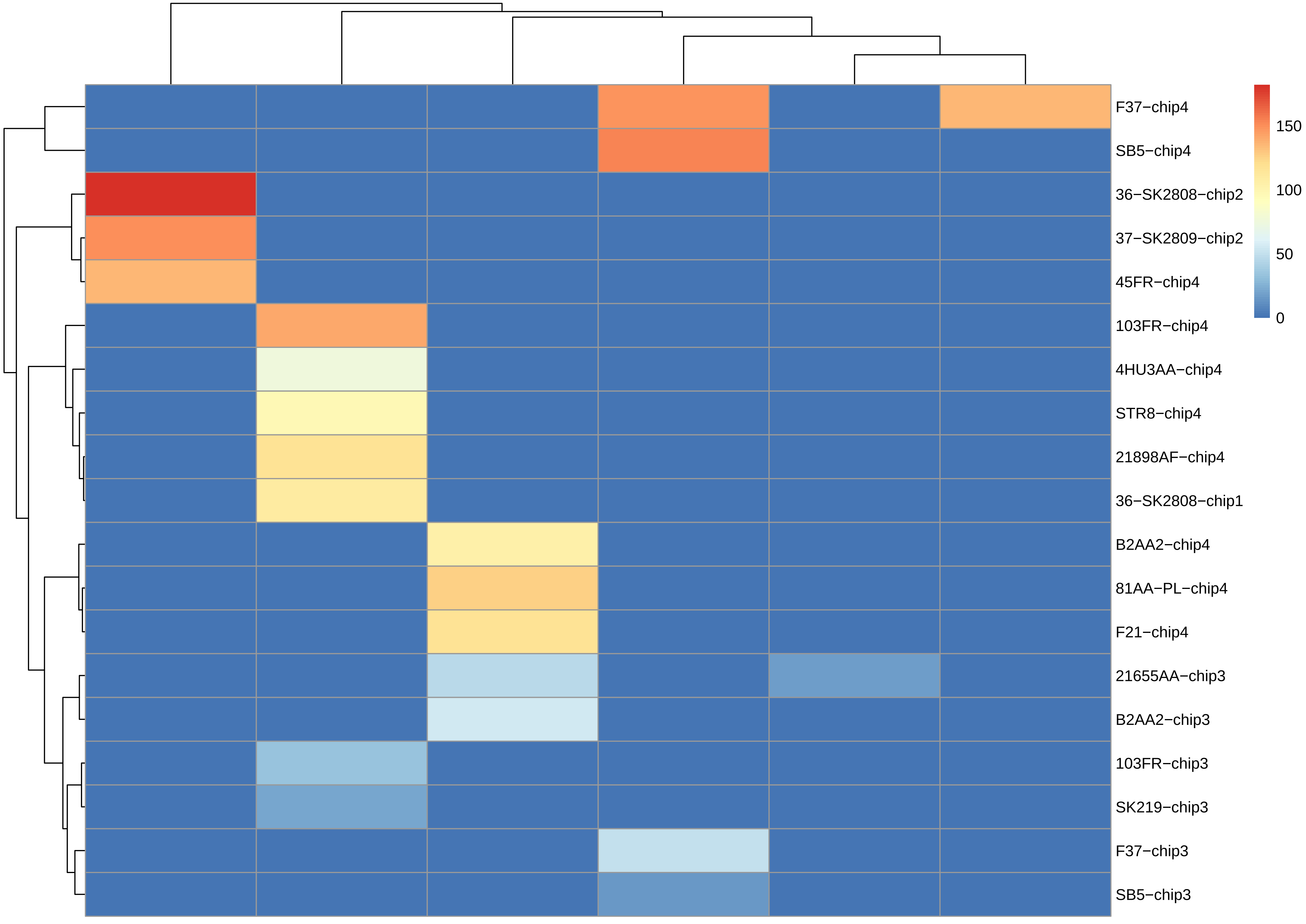
F37−chip4
SB5−chip4
36−SK2808−chip2
37−SK2809−chip2
45FR−chip4
103FR−chip4
4HU3AA−chip4
STR8−chip4
21898AF−chip4
36−SK2808−chip1
B2AA2−chip4
81AA−PL−chip4
F21−chip4
21655AA−chip3
B2AA2−chip3
103FR−chip3
SK219−chip3
F37−chip3
SB5−chip3
150
100
50
0

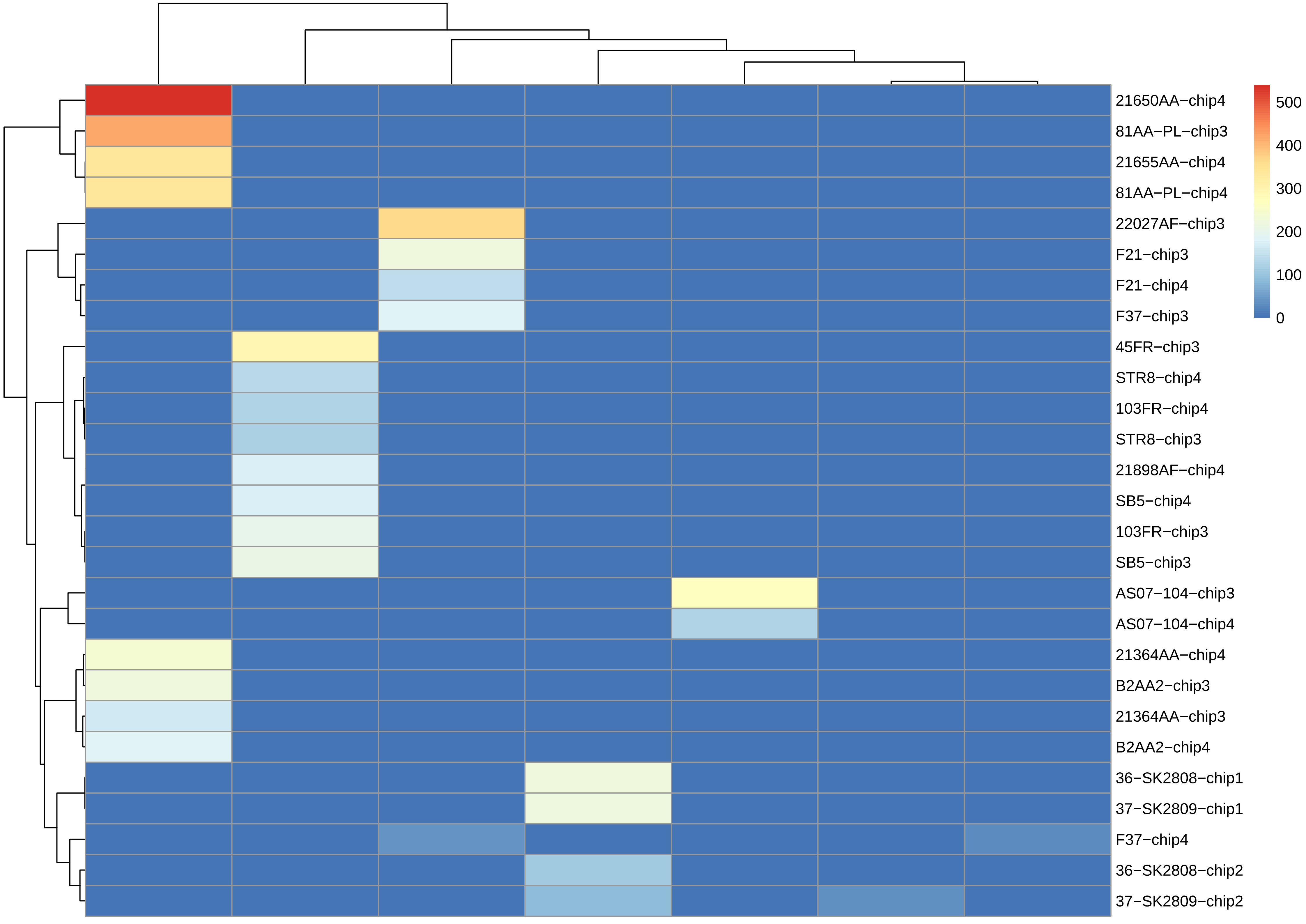

21650AA−chip4
81AA−PL−chip3
21655AA−chip4
81AA−PL−chip4
22027AF−chip3
F21−chip3
F21−chip4
F37−chip3
45FR−chip3
STR8−chip4
103FR−chip4
STR8−chip3
21898AF−chip4
SB5−chip4
103FR−chip3
SB5−chip3
AS07−104−chip3
AS07−104−chip4
21364AA−chip4
B2AA2−chip3
21364AA−chip3
B2AA2−chip4
36−SK2808−chip1
37−SK2809−chip1
F37−chip4
36−SK2808−chip2
37−SK2809−chip2
500
400
300
200
100
0

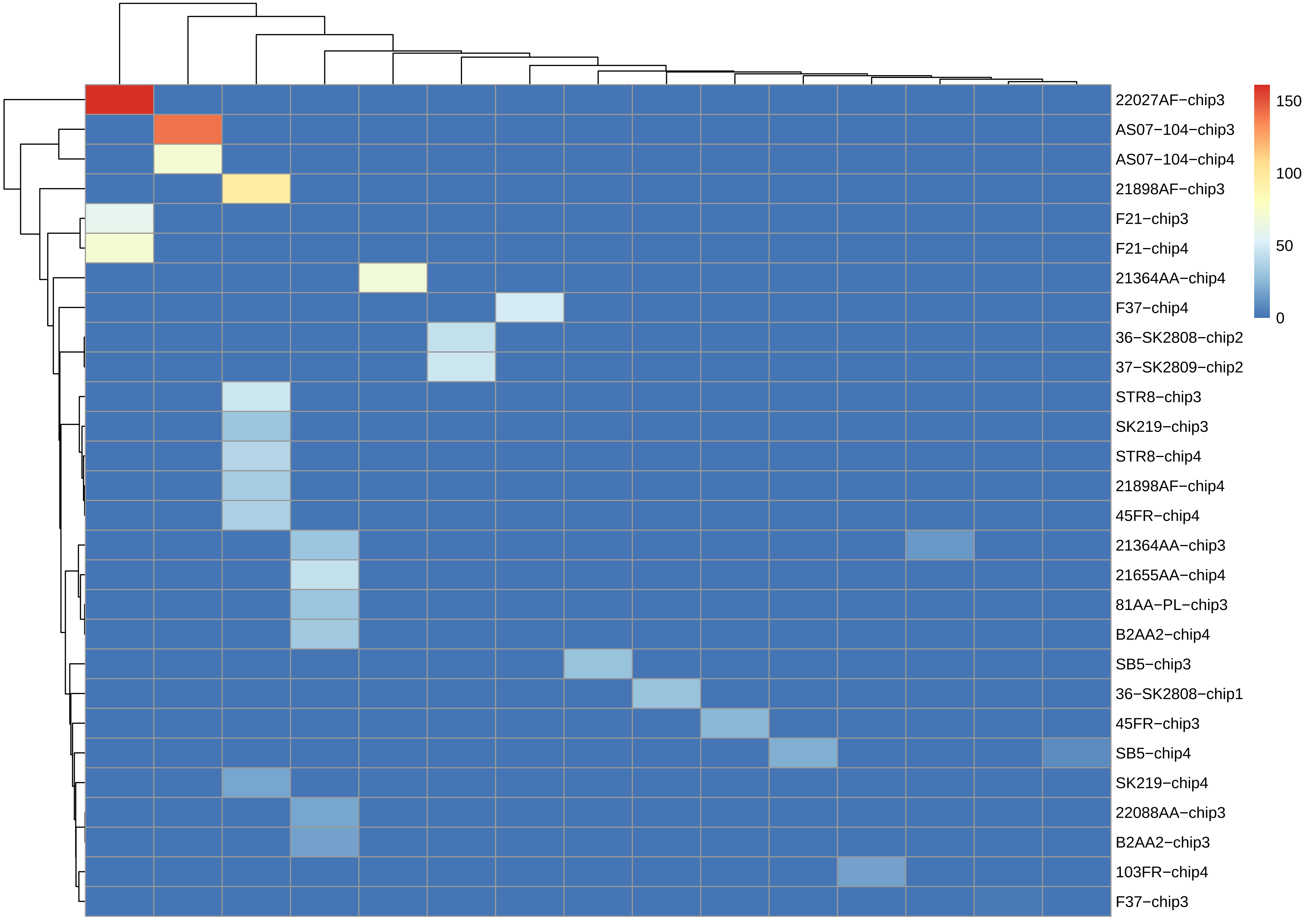

22027AF−chip3
AS07−104−chip3
AS07−104−chip4
21898AF−chip3
F21−chip3
F21−chip4
21364AA−chip4
F37−chip4
36−SK2808−chip2
37−SK2809−chip2
STR8−chip3
SK219−chip3
STR8−chip4
21898AF−chip4
45FR−chip4
21364AA−chip3
21655AA−chip4
81AA−PL−chip3
B2AA2−chip4
SB5−chip3
36−SK2808−chip1
45FR−chip3
SB5−chip4
SK219−chip4
22088AA−chip3
B2AA2−chip3
103FR−chip4
F37−chip3
150
100
50
0

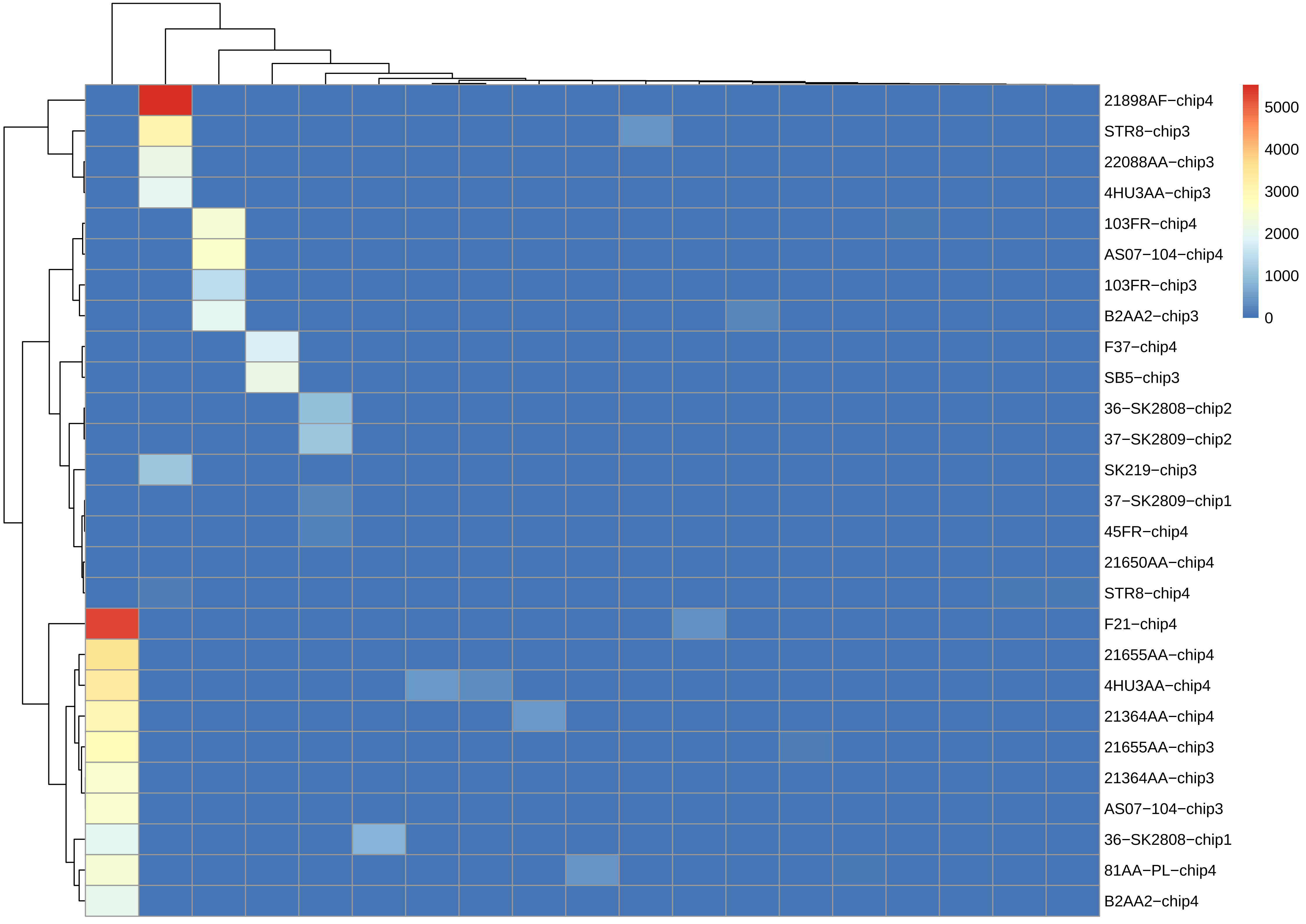
21898AF−chip4
STR8−chip3
22088AA−chip3
4HU3AA−chip3
103FR−chip4
AS07−104−chip4
103FR−chip3
B2AA2−chip3
F37−chip4
SB5−chip3
36−SK2808−chip2
37−SK2809−chip2
SK219−chip3
37−SK2809−chip1
45FR−chip4
21650AA−chip4
STR8−chip4
F21−chip4
21655AA−chip4
4HU3AA−chip4
21364AA−chip4
21655AA−chip3
21364AA−chip3
AS07−104−chip3
36−SK2808−chip1
81AA−PL−chip4
B2AA2−chip4
5000
4000
3000
2000
1000
0

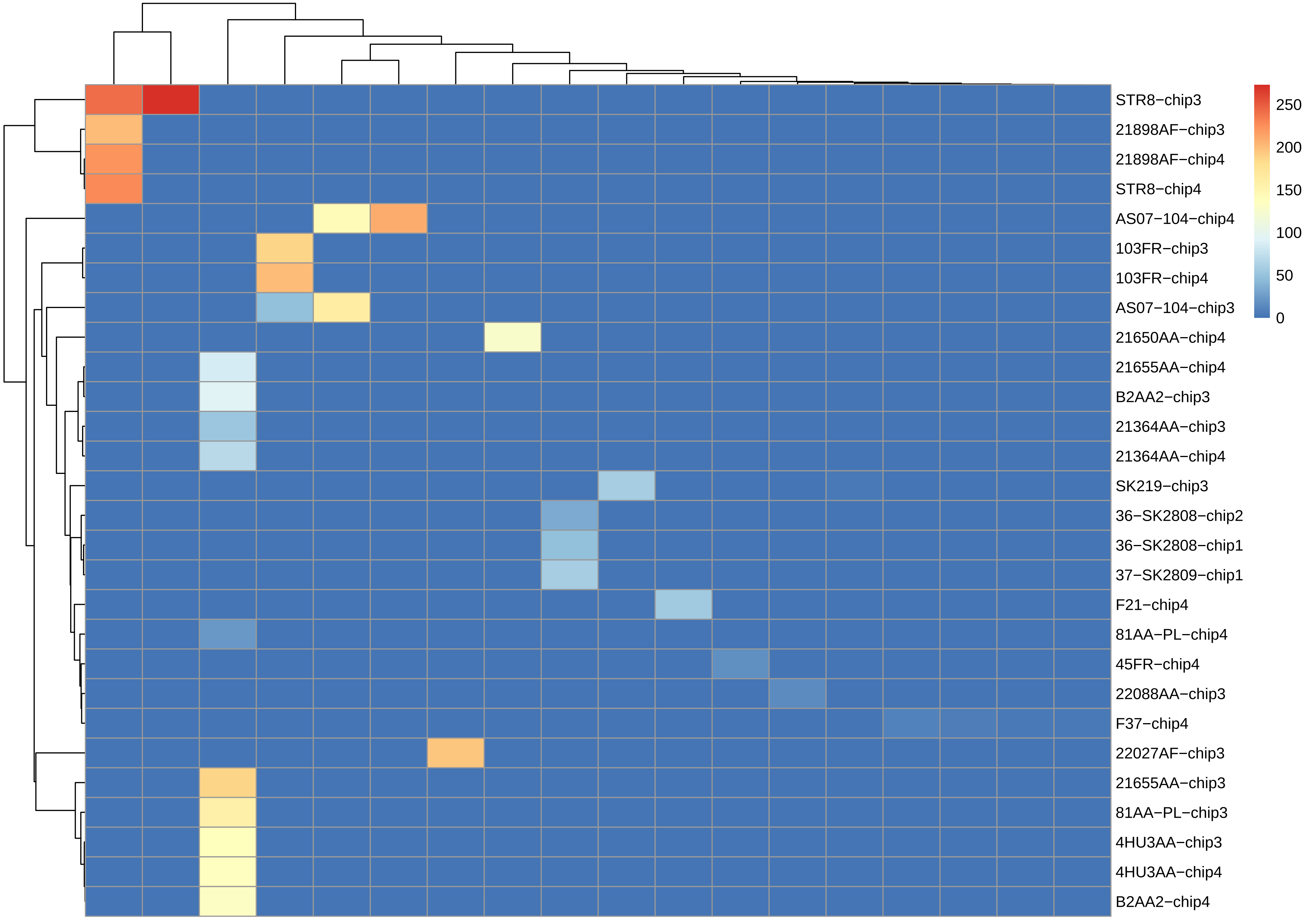

STR8−chip3
21898AF−chip3
21898AF−chip4
STR8−chip4
AS07−104−chip4
103FR−chip3
103FR−chip4
AS07−104−chip3
21650AA−chip4
21655AA−chip4
B2AA2−chip3
21364AA−chip3
21364AA−chip4
SK219−chip3
36−SK2808−chip2
36−SK2808−chip1
37−SK2809−chip1
F21−chip4
81AA−PL−chip4
45FR−chip4
22088AA−chip3
F37−chip4
22027AF−chip3
21655AA−chip3
81AA−PL−chip3
4HU3AA−chip3
4HU3AA−chip4
B2AA2−chip4
250
200
150
100
50
0

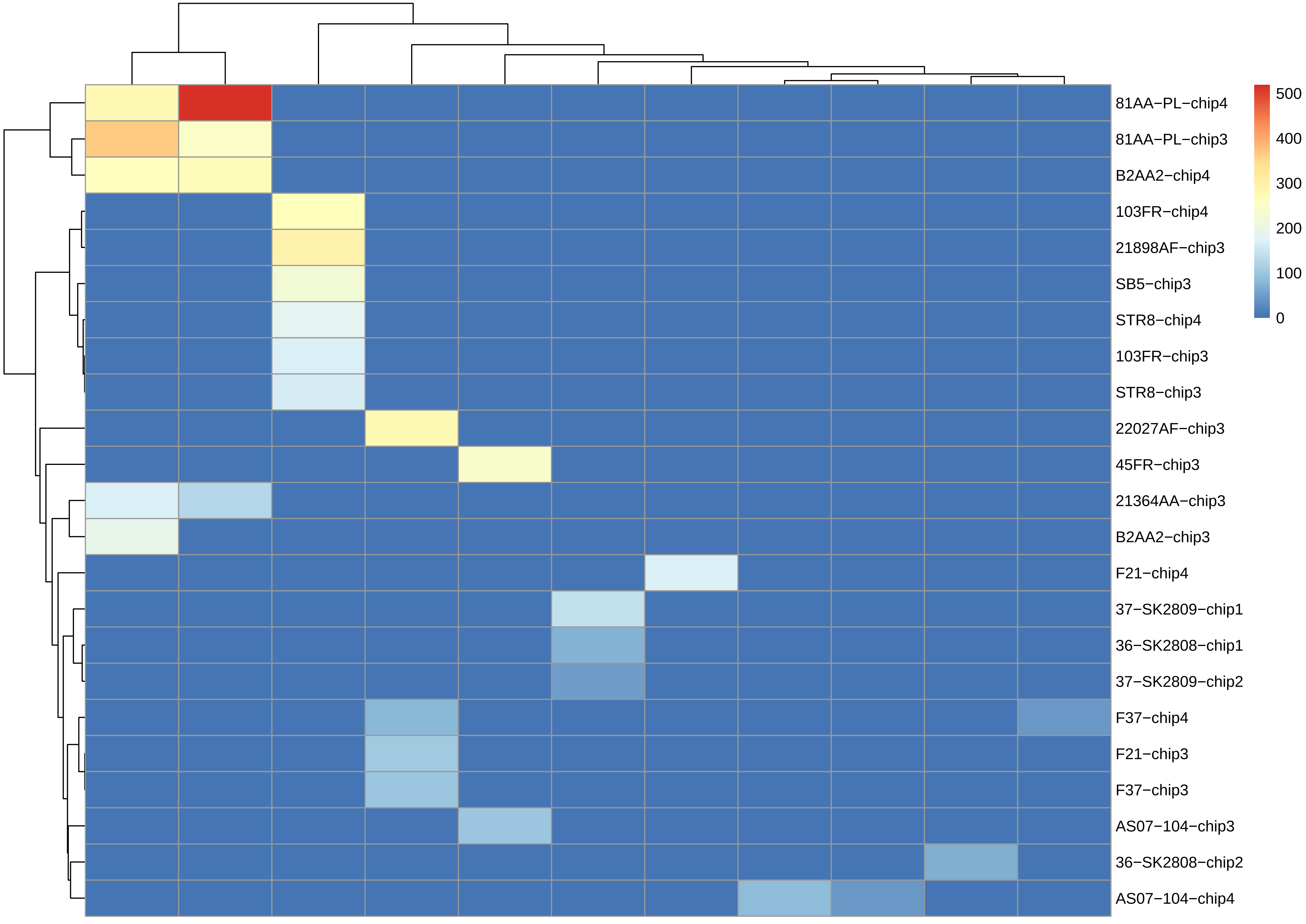

81AA−PL−chip4
81AA−PL−chip3
B2AA2−chip4
103FR−chip4
21898AF−chip3
SB5−chip3
STR8−chip4
103FR−chip3
STR8−chip3
22027AF−chip3
45FR−chip3
21364AA−chip3
B2AA2−chip3
F21−chip4
37−SK2809−chip1
36−SK2808−chip1
37−SK2809−chip2
F37−chip4
F21−chip3
F37−chip3
AS07−104−chip3
36−SK2808−chip2
AS07−104−chip4
500
400
300
200
100
0

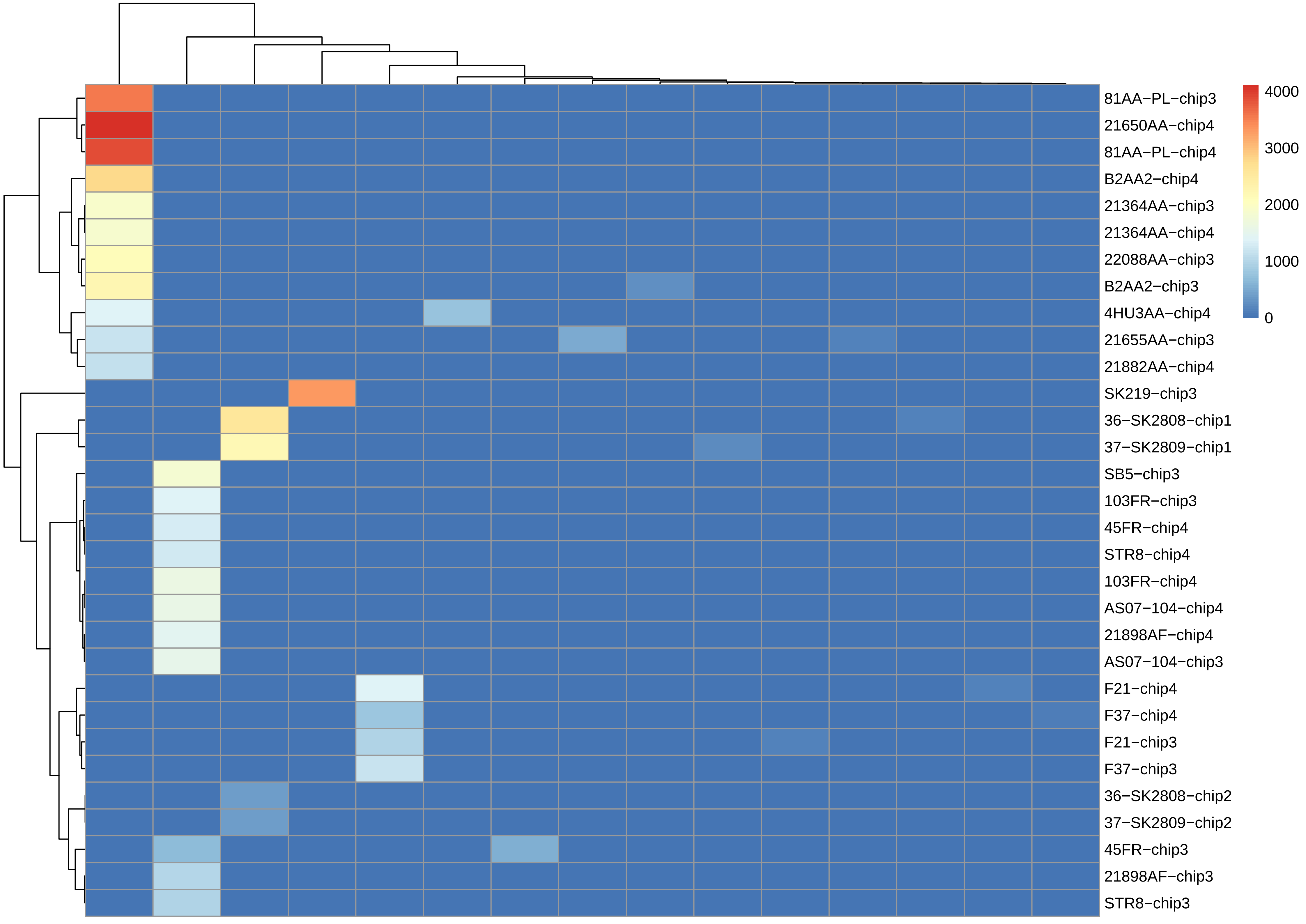

81AA−PL−chip3
21650AA−chip4
81AA−PL−chip4
B2AA2−chip4
21364AA−chip3
21364AA−chip4
22088AA−chip3
B2AA2−chip3
4HU3AA−chip4
21655AA−chip3
21882AA−chip4
SK219−chip3
36−SK2808−chip1
37−SK2809−chip1
SB5−chip3
103FR−chip3
45FR−chip4
STR8−chip4
103FR−chip4
AS07−104−chip4
21898AF−chip4
AS07−104−chip3
F21−chip4
F37−chip4
F21−chip3
F37−chip3
36−SK2808−chip2
37−SK2809−chip2
45FR−chip3
21898AF−chip3
STR8−chip3
4000
3000
2000
1000
0

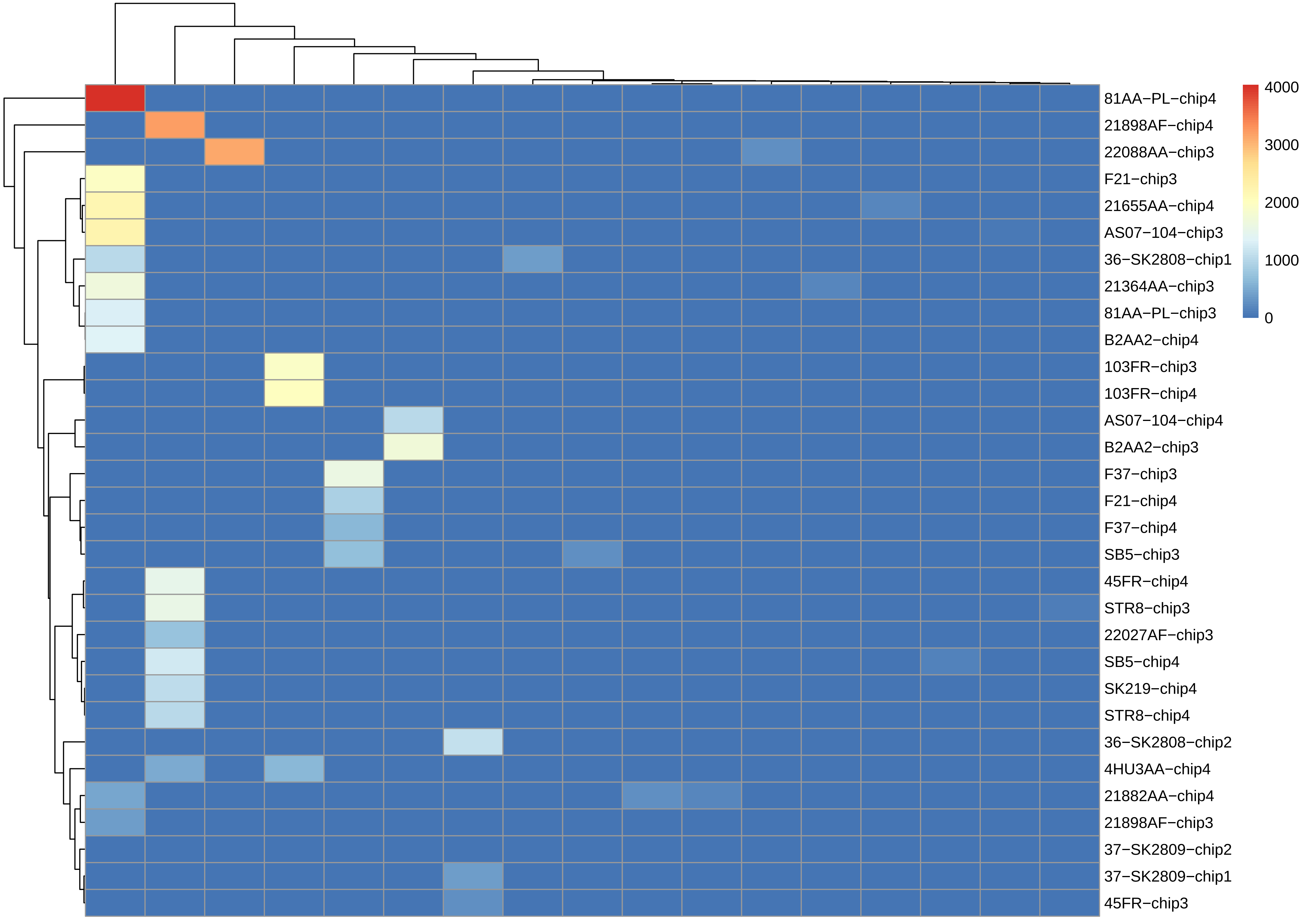
81AA−PL−chip4
21898AF−chip4
22088AA−chip3
F21−chip3
21655AA−chip4
AS07−104−chip3
36−SK2808−chip1
21364AA−chip3
81AA−PL−chip3
B2AA2−chip4
103FR−chip3
103FR−chip4
AS07−104−chip4
B2AA2−chip3
F37−chip3
F21−chip4
F37−chip4
SB5−chip3
45FR−chip4
STR8−chip3
22027AF−chip3
SB5−chip4
SK219−chip4
STR8−chip4
36−SK2808−chip2
4HU3AA−chip4
21882AA−chip4
21898AF−chip3
37−SK2809−chip2
37−SK2809−chip1
45FR−chip3
4000
3000
2000
1000
0

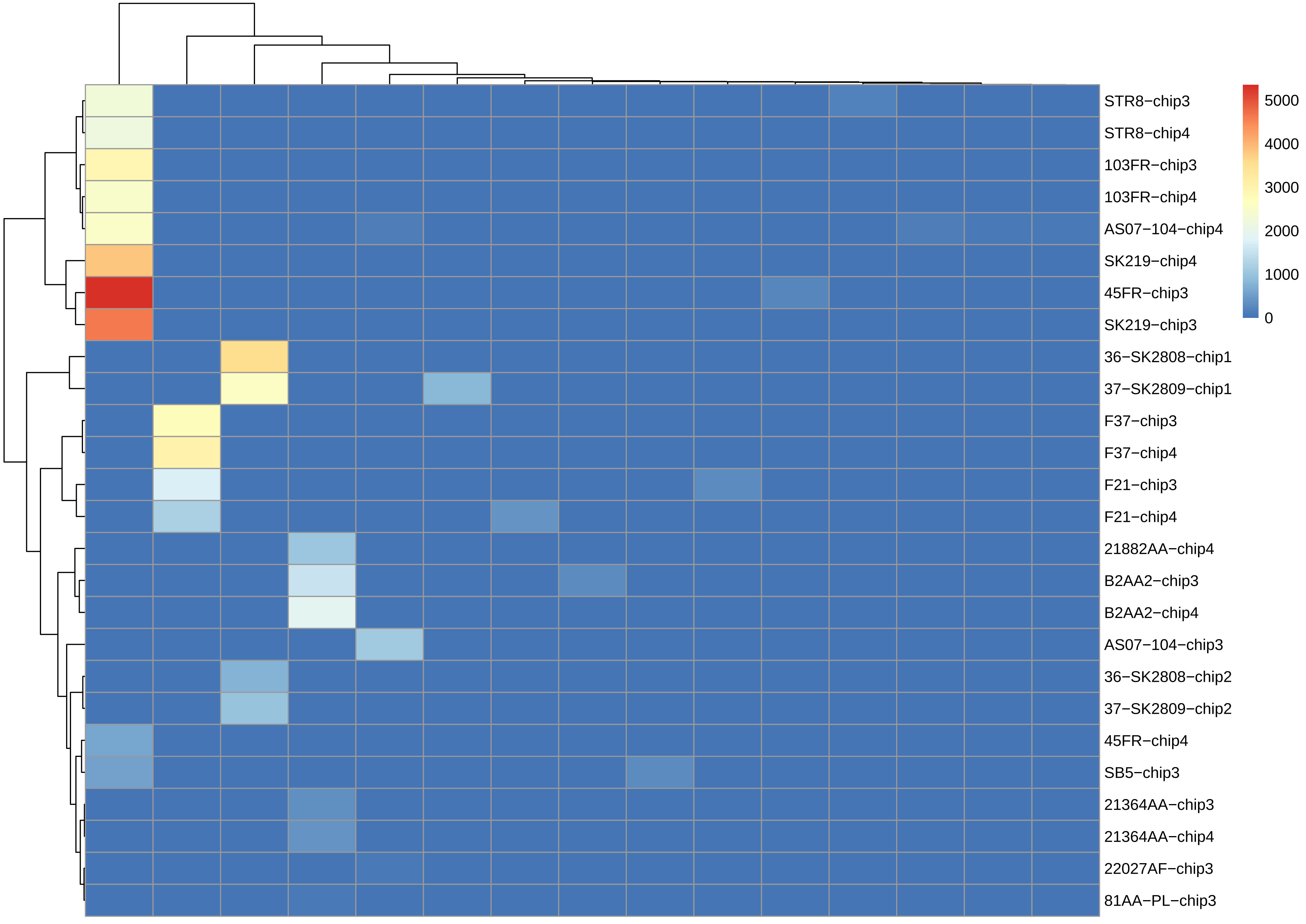
STR8−chip3
STR8−chip4
103FR−chip3
103FR−chip4
AS07−104−chip4
SK219−chip4
45FR−chip3
SK219−chip3
36−SK2808−chip1
37−SK2809−chip1
F37−chip3
F37−chip4
F21−chip3
F21−chip4
21882AA−chip4
B2AA2−chip3
B2AA2−chip4
AS07−104−chip3
36−SK2808−chip2
37−SK2809−chip2
45FR−chip4
SB5−chip3
21364AA−chip3
21364AA−chip4
22027AF−chip3
81AA−PL−chip3
5000
4000
3000
2000
1000
0

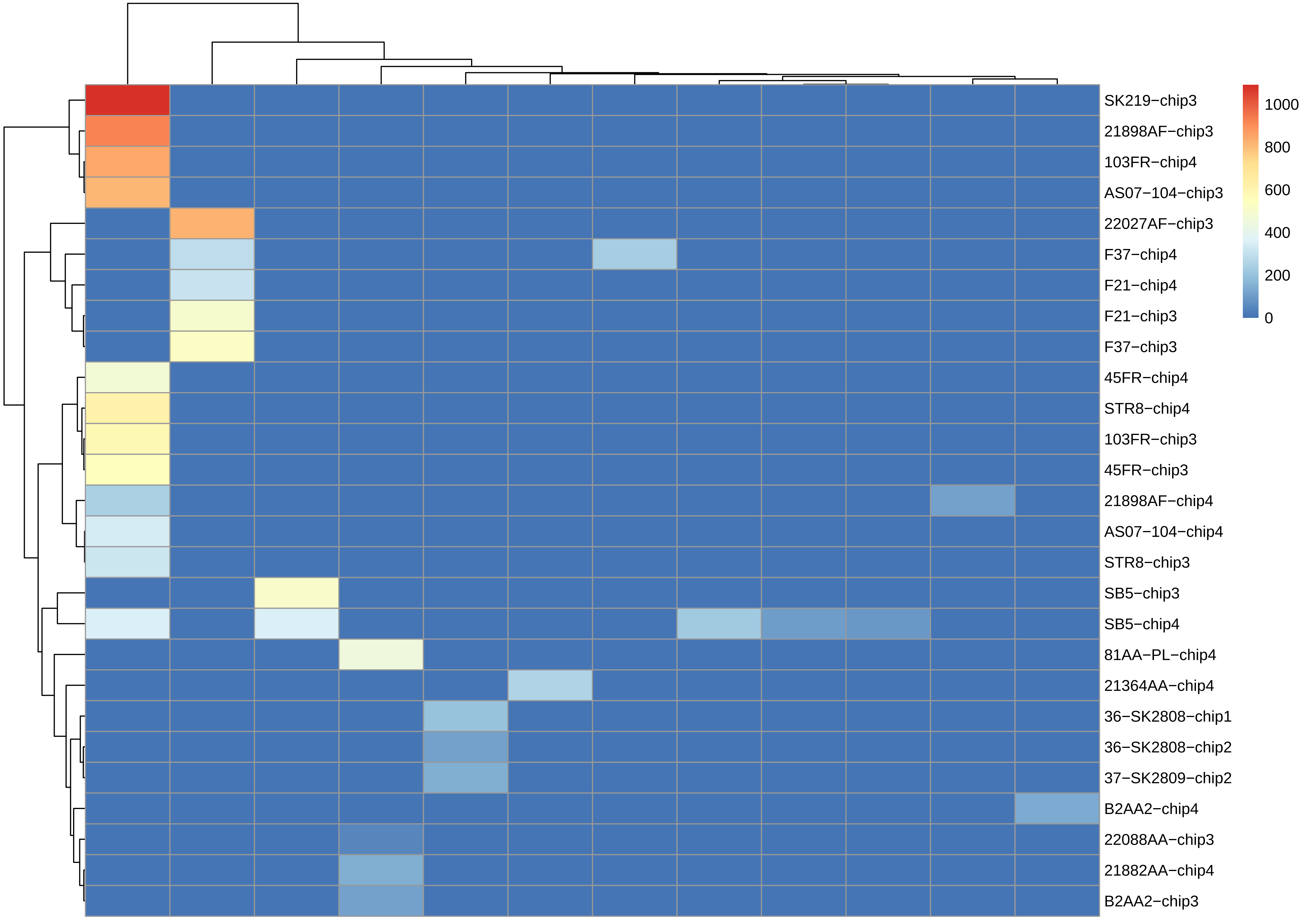

SK219−chip3
21898AF−chip3
103FR−chip4
AS07−104−chip3
22027AF−chip3
F37−chip4
F21−chip4
F21−chip3
F37−chip3
45FR−chip4
STR8−chip4
103FR−chip3
45FR−chip3
21898AF−chip4
AS07−104−chip4
STR8−chip3
SB5−chip3
SB5−chip4
81AA−PL−chip4
21364AA−chip4
36−SK2808−chip1
36−SK2808−chip2
37−SK2809−chip2
B2AA2−chip4
22088AA−chip3
21882AA−chip4
B2AA2−chip3
1000
800
600
400
200
0

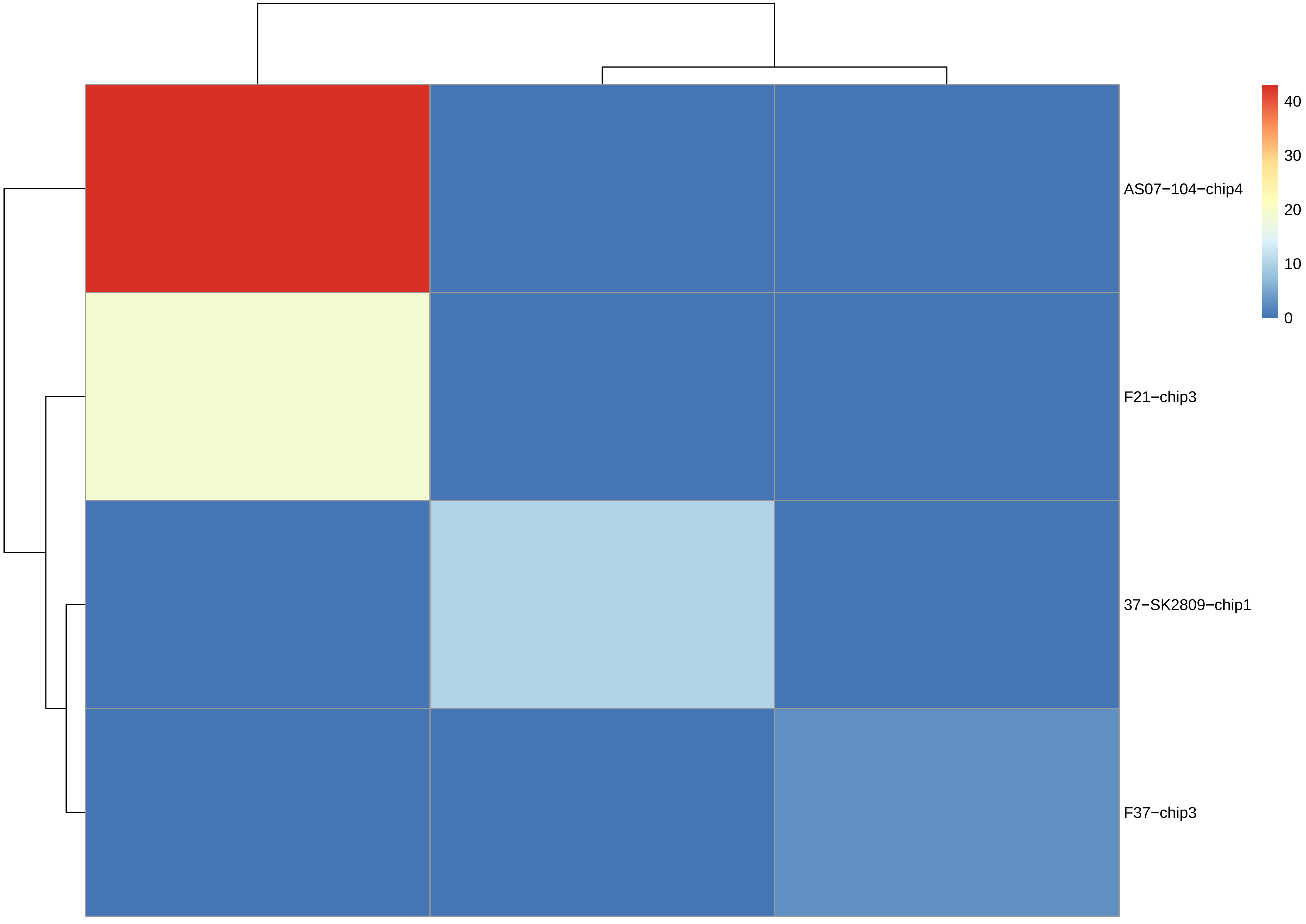

AS07−104−chip4
F21−chip3
37−SK2809−chip1
F37−chip3
40
30
20
10
0

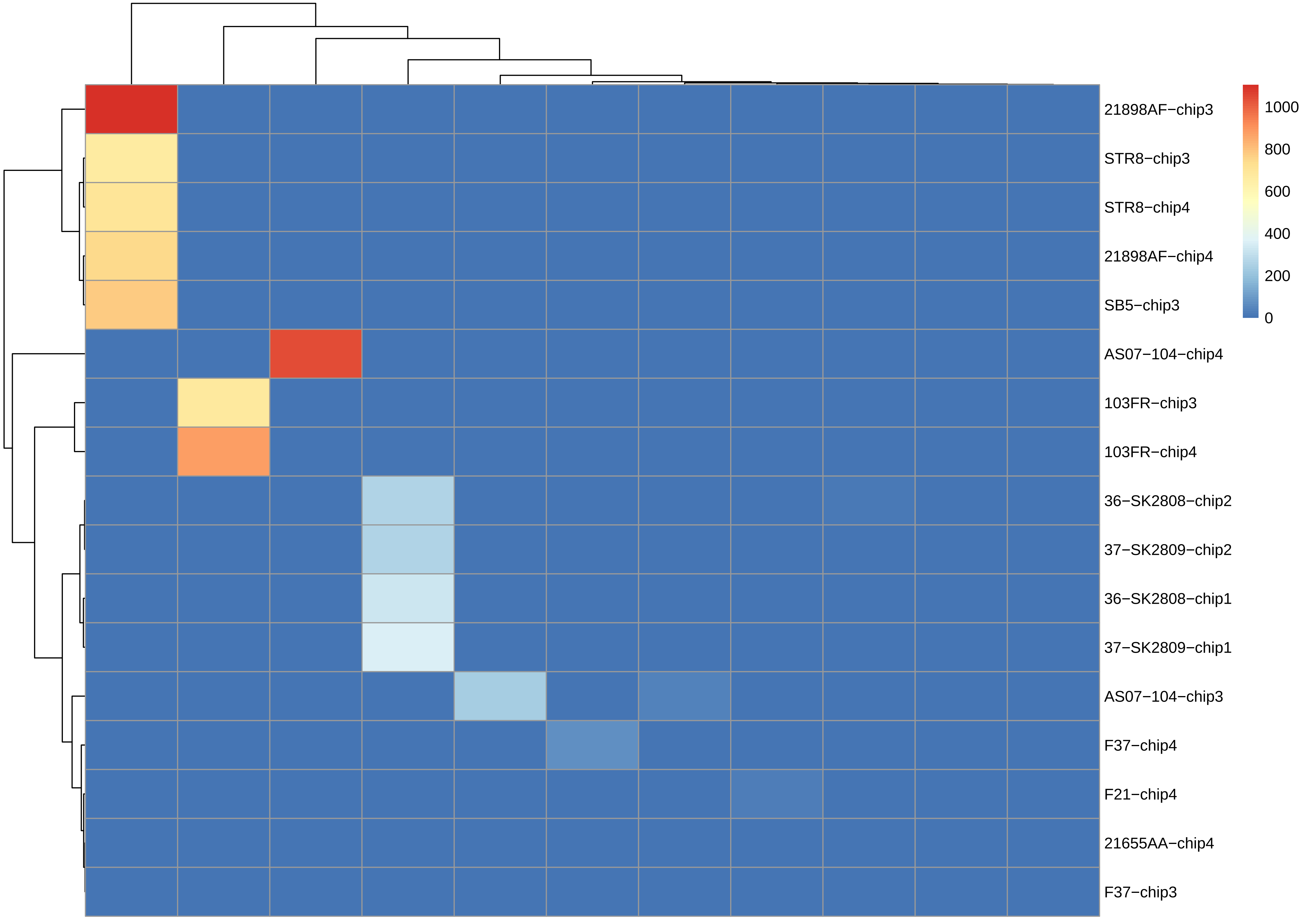
21898AF−chip3
STR8−chip3
STR8−chip4
21898AF−chip4
SB5−chip3
AS07−104−chip4
103FR−chip3
103FR−chip4
36−SK2808−chip2
37−SK2809−chip2
36−SK2808−chip1
37−SK2809−chip1
AS07−104−chip3
F37−chip4
F21−chip4
21655AA−chip4
F37−chip3
1000
800
600
400
200
0

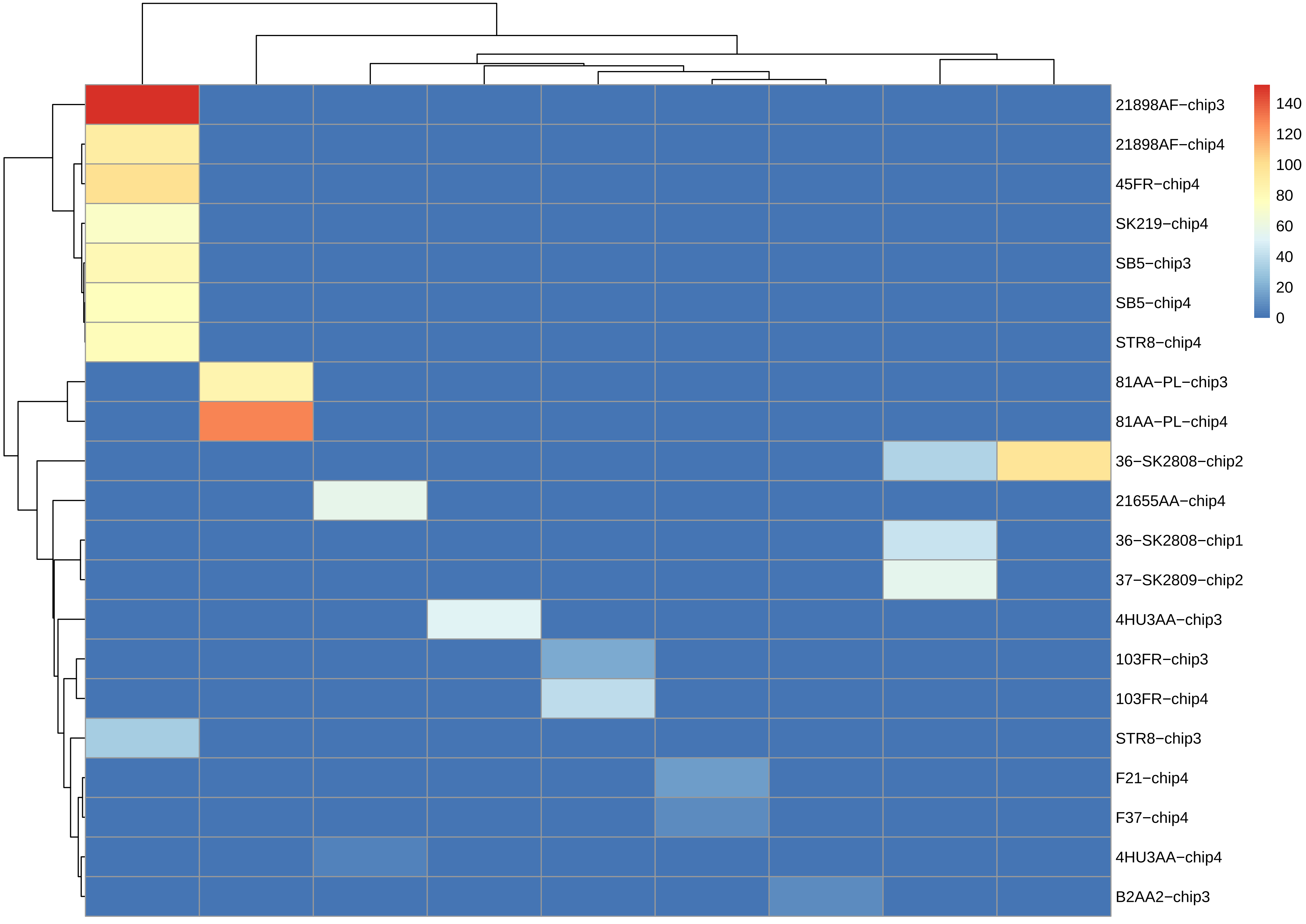
21898AF−chip3
21898AF−chip4
45FR−chip4
SK219−chip4
SB5−chip3
SB5−chip4
STR8−chip4
81AA−PL−chip3
81AA−PL−chip4
36−SK2808−chip2
21655AA−chip4
36−SK2808−chip1
37−SK2809−chip2
4HU3AA−chip3
103FR−chip3
103FR−chip4
STR8−chip3
F21−chip4
F37−chip4
4HU3AA−chip4
B2AA2−chip3
140
120
100
80
60
40
20
0

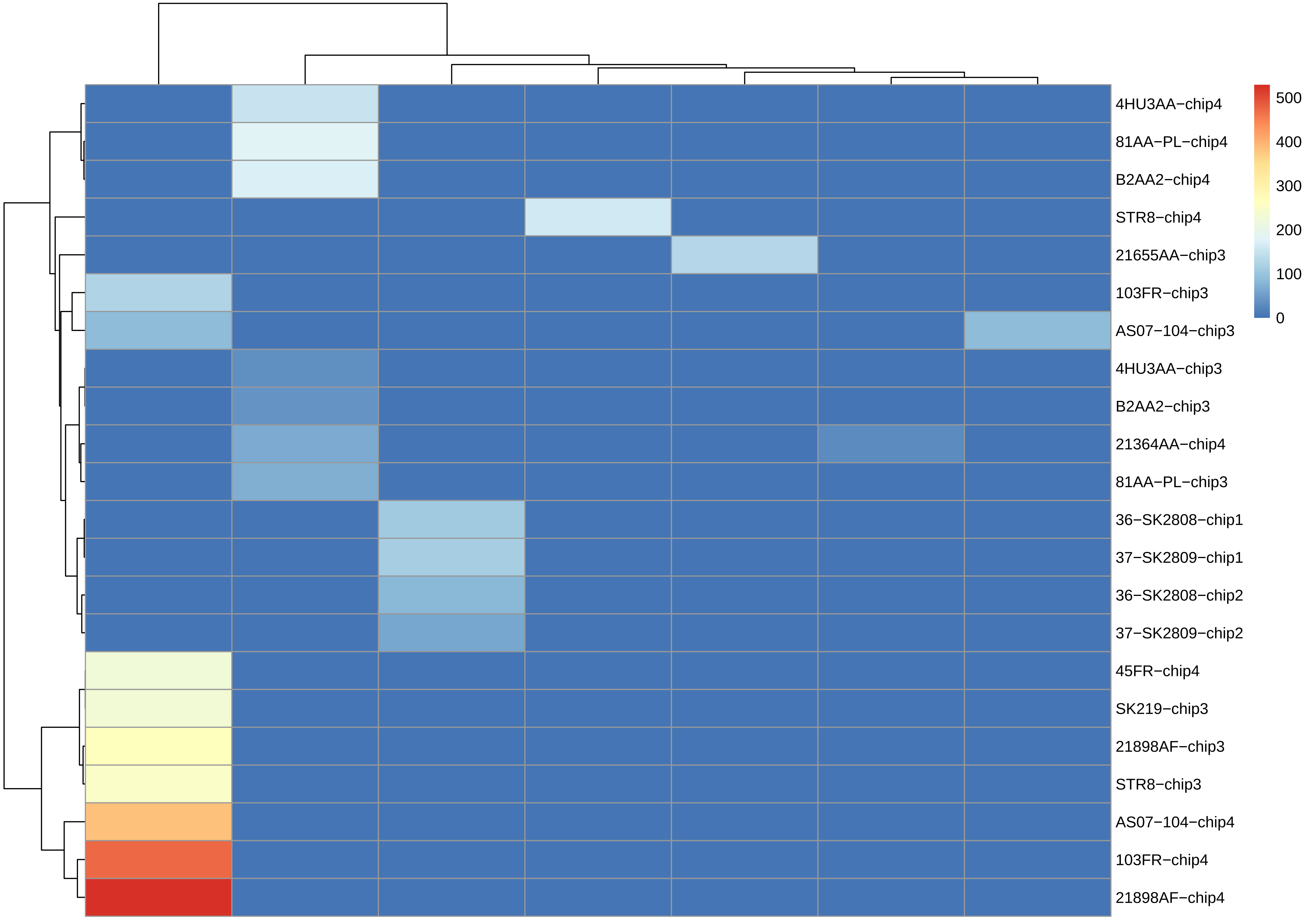

4HU3AA−chip4
81AA−PL−chip4
B2AA2−chip4
STR8−chip4
21655AA−chip3
103FR−chip3
AS07−104−chip3
4HU3AA−chip3
B2AA2−chip3
21364AA−chip4
81AA−PL−chip3
36−SK2808−chip1
37−SK2809−chip1
36−SK2808−chip2
37−SK2809−chip2
45FR−chip4
SK219−chip3
21898AF−chip3
STR8−chip3
AS07−104−chip4
103FR−chip4
21898AF−chip4
500
400
300
200
100
0

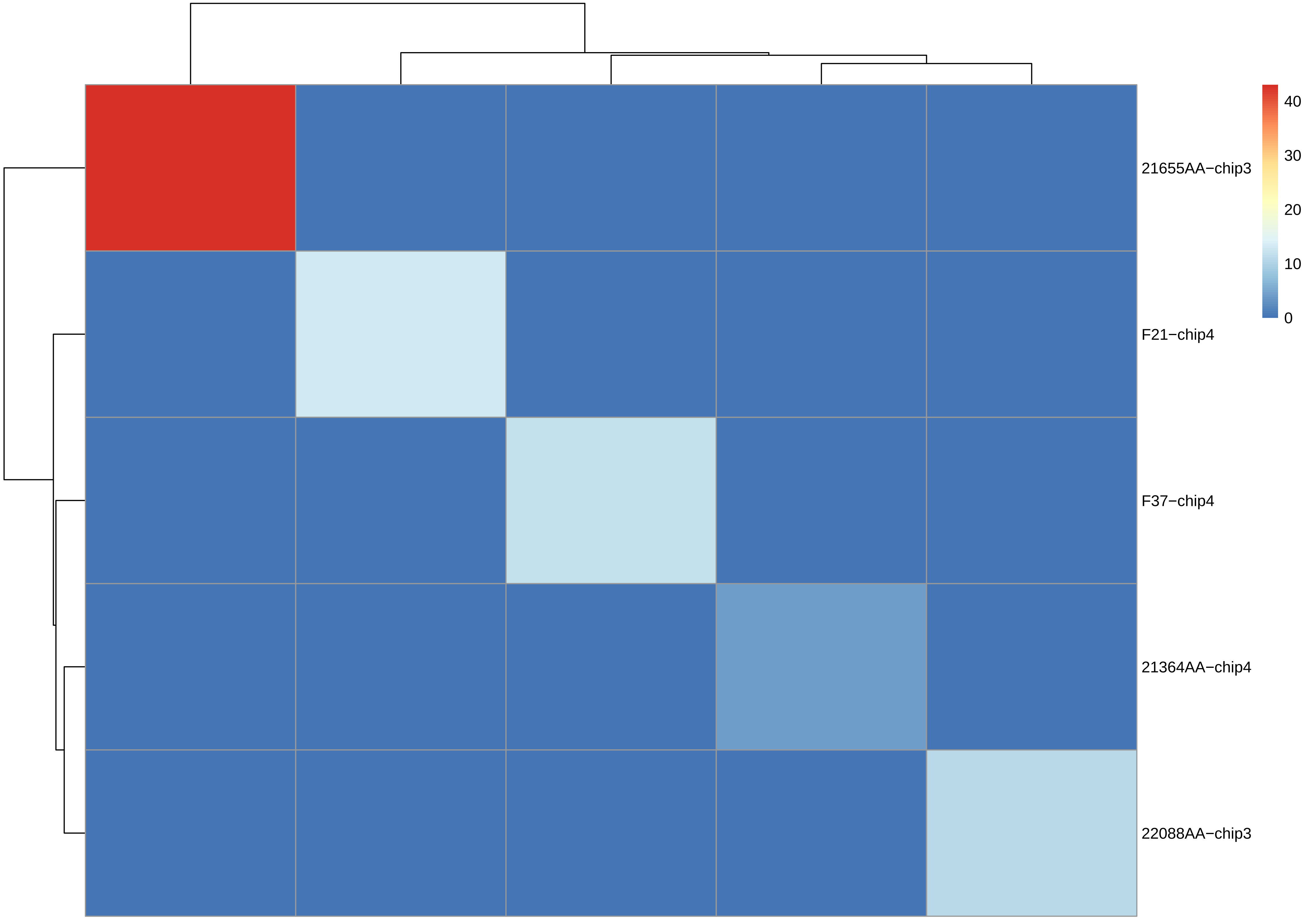
21655AA−chip3
F21−chip4
F37−chip4
21364AA−chip4
22088AA−chip3
40
30
20
10
0

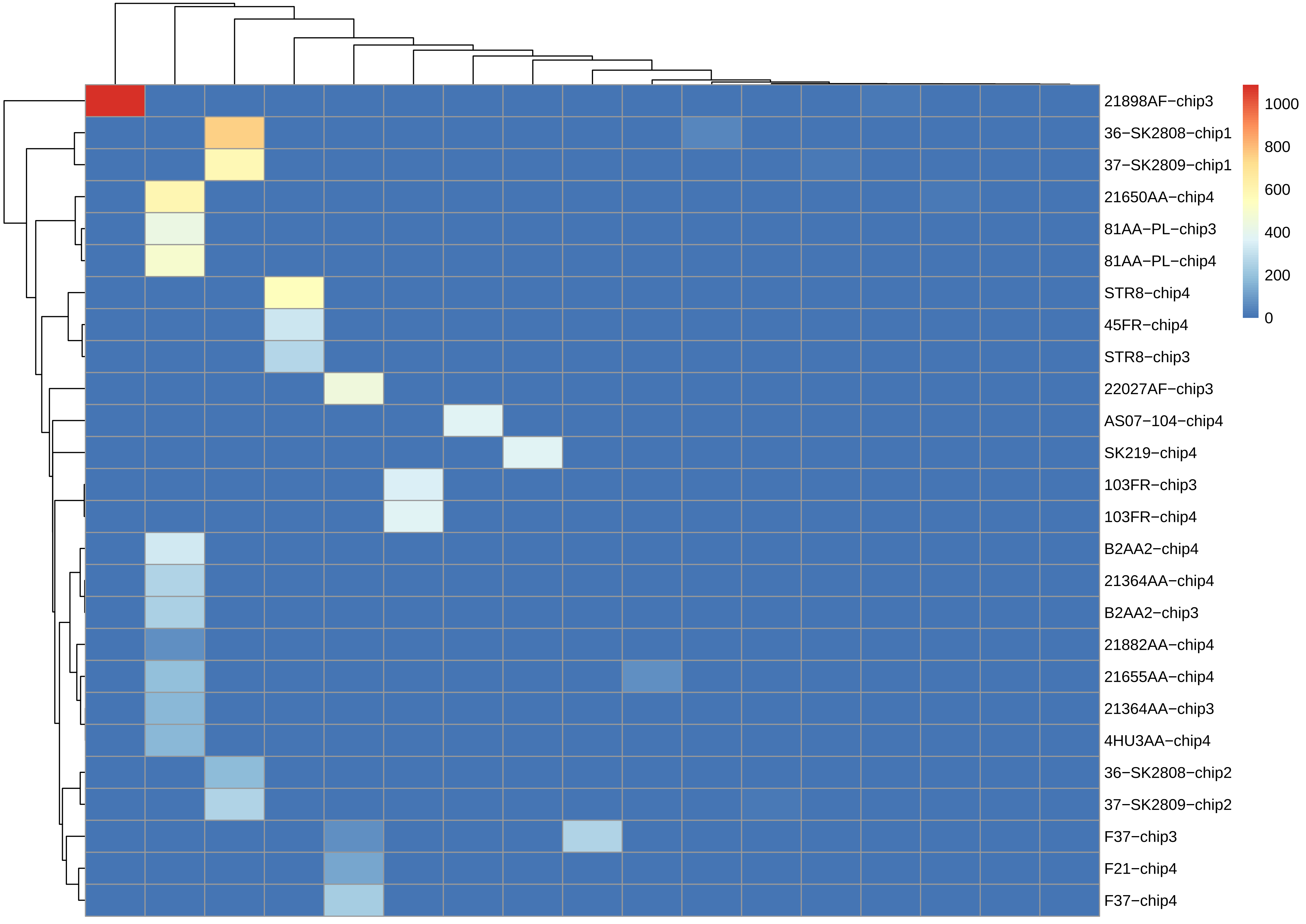
21898AF−chip3
36−SK2808−chip1
37−SK2809−chip1
21650AA−chip4
81AA−PL−chip3
81AA−PL−chip4
STR8−chip4
45FR−chip4
STR8−chip3
22027AF−chip3
AS07−104−chip4
SK219−chip4
103FR−chip3
103FR−chip4
B2AA2−chip4
21364AA−chip4
B2AA2−chip3
21882AA−chip4
21655AA−chip4
21364AA−chip3
4HU3AA−chip4
36−SK2808−chip2
37−SK2809−chip2
F37−chip3
F21−chip4
F37−chip4
1000
800
600
400
200
0

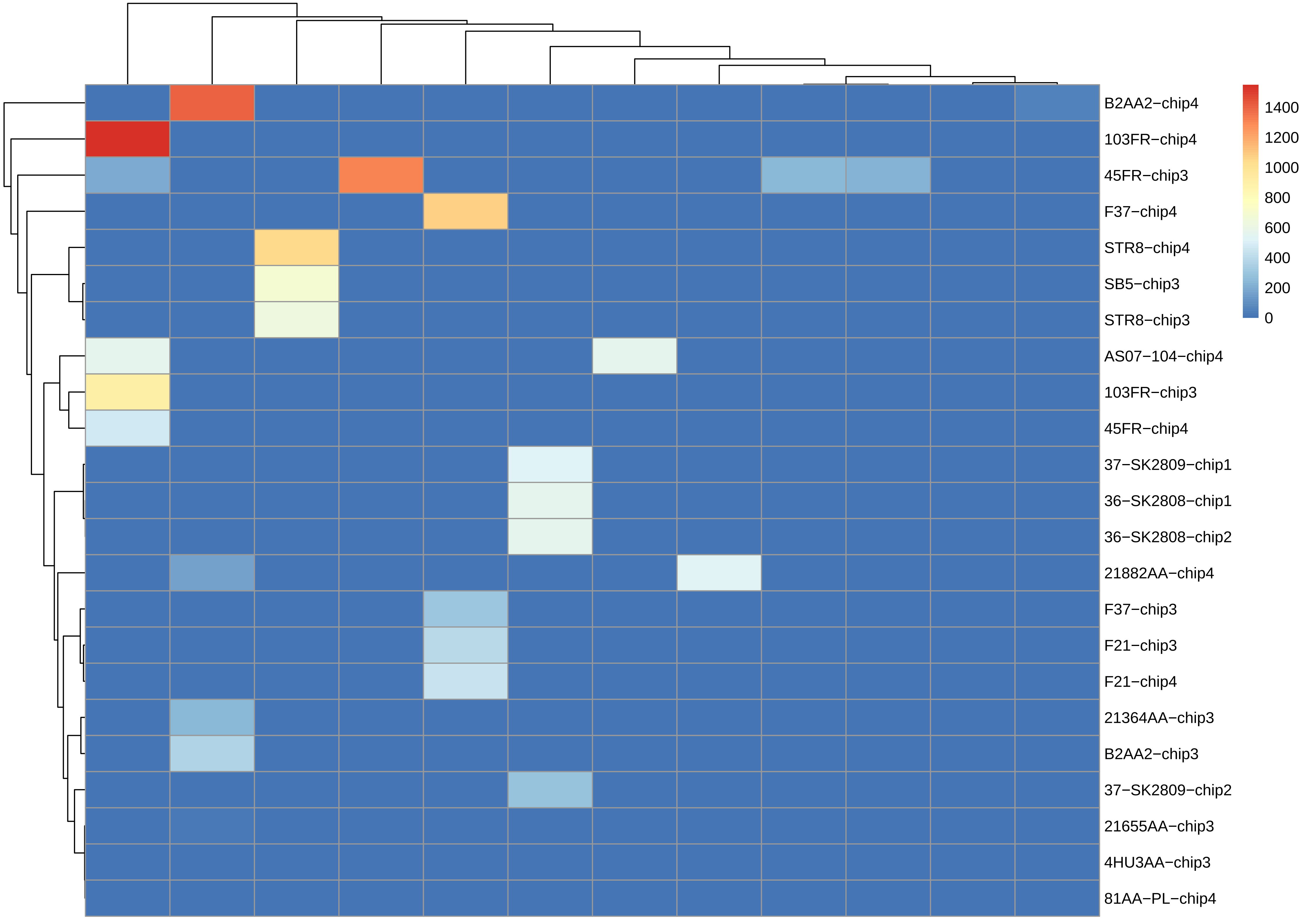
B2AA2−chip4
103FR−chip4
45FR−chip3
F37−chip4
STR8−chip4
SB5−chip3
STR8−chip3
AS07−104−chip4
103FR−chip3
45FR−chip4
37−SK2809−chip1
36−SK2808−chip1
36−SK2808−chip2
21882AA−chip4
F37−chip3
F21−chip3
F21−chip4
21364AA−chip3
B2AA2−chip3
37−SK2809−chip2
21655AA−chip3
4HU3AA−chip3
81AA−PL−chip4
1400
1200
1000
800
600
400
200
0

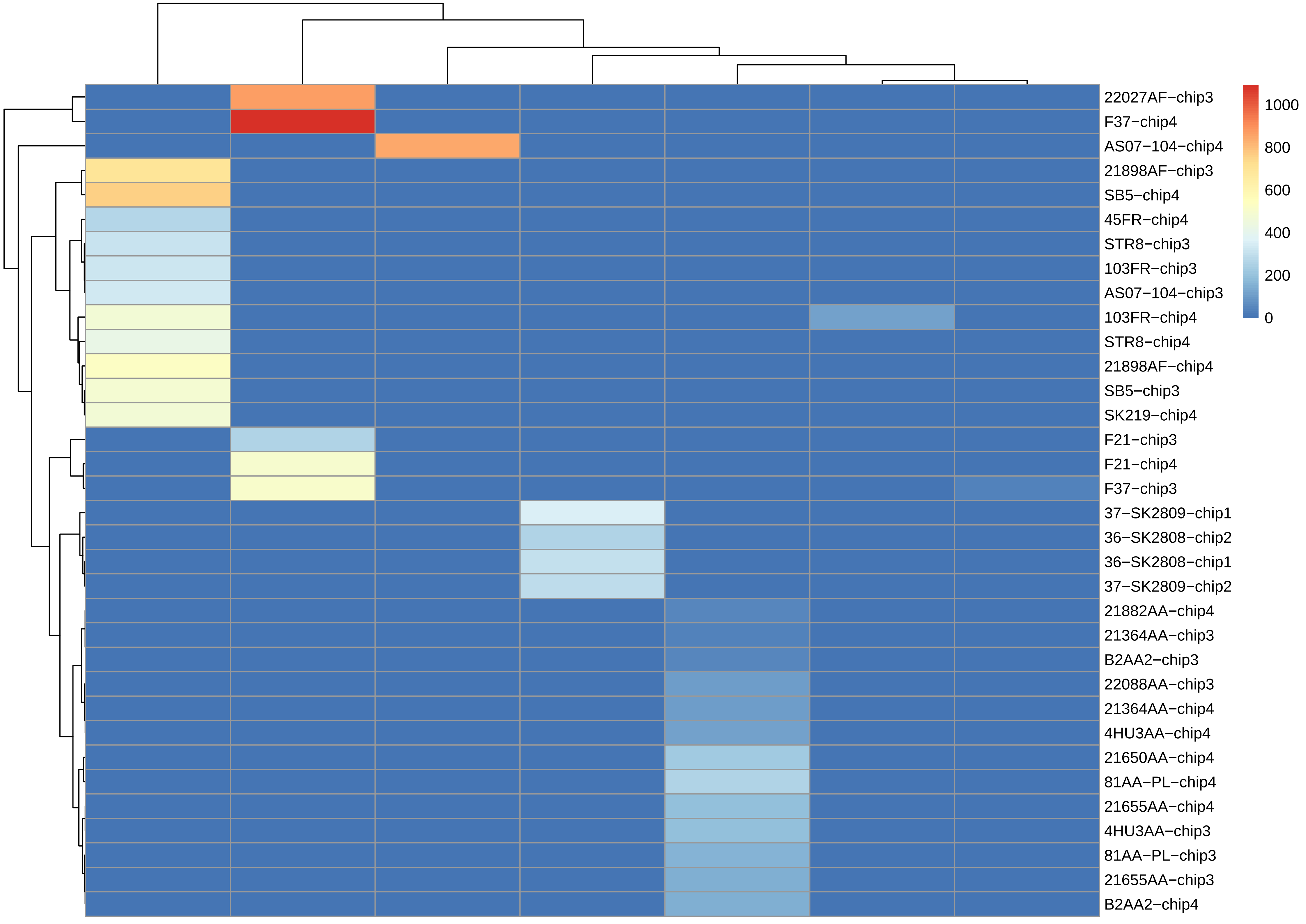

22027AF−chip3
F37−chip4
AS07−104−chip4
21898AF−chip3
SB5−chip4
45FR−chip4
STR8−chip3
103FR−chip3
AS07−104−chip3
103FR−chip4
STR8−chip4
21898AF−chip4
SB5−chip3
SK219−chip4
F21−chip3
F21−chip4
F37−chip3
37−SK2809−chip1
36−SK2808−chip2
36−SK2808−chip1
37−SK2809−chip2
21882AA−chip4
21364AA−chip3
B2AA2−chip3
22088AA−chip3
21364AA−chip4
4HU3AA−chip4
21650AA−chip4
81AA−PL−chip4
21655AA−chip4
4HU3AA−chip3
81AA−PL−chip3
21655AA−chip3
B2AA2−chip4
1000
800
600
400
200
0

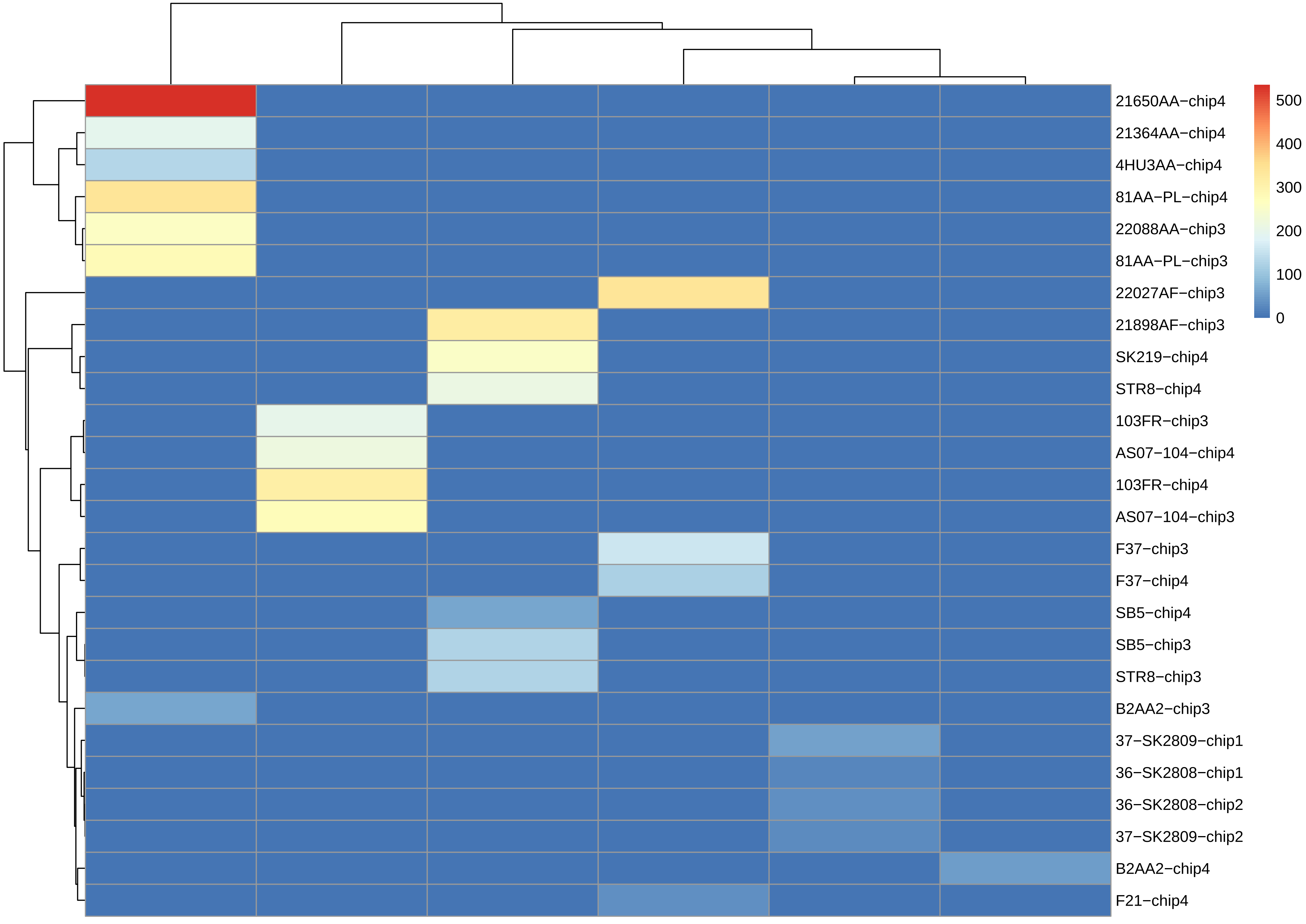

21650AA−chip4
21364AA−chip4
4HU3AA−chip4
81AA−PL−chip4
22088AA−chip3
81AA−PL−chip3
22027AF−chip3
21898AF−chip3
SK219−chip4
STR8−chip4
103FR−chip3
AS07−104−chip4
103FR−chip4
AS07−104−chip3
F37−chip3
F37−chip4
SB5−chip4
SB5−chip3
STR8−chip3
B2AA2−chip3
37−SK2809−chip1
36−SK2808−chip1
36−SK2808−chip2
37−SK2809−chip2
B2AA2−chip4
F21−chip4
500
400
300
200
100
0

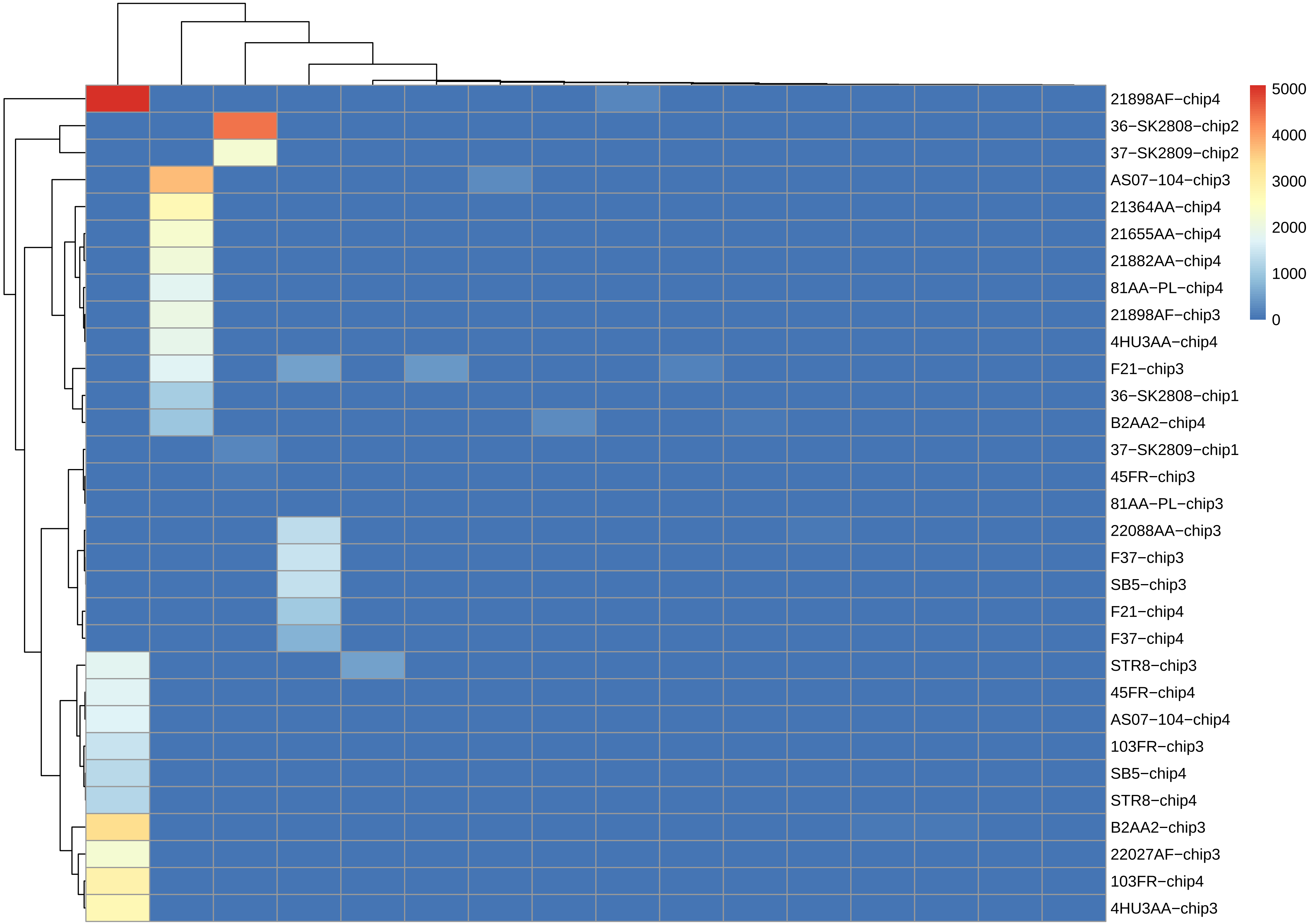
21898AF−chip4
36−SK2808−chip2
37−SK2809−chip2
AS07−104−chip3
21364AA−chip4
21655AA−chip4
21882AA−chip4
81AA−PL−chip4
21898AF−chip3
4HU3AA−chip4
F21−chip3
36−SK2808−chip1
B2AA2−chip4
37−SK2809−chip1
45FR−chip3
81AA−PL−chip3
22088AA−chip3
F37−chip3
SB5−chip3
F21−chip4
F37−chip4
STR8−chip3
45FR−chip4
AS07−104−chip4
103FR−chip3
SB5−chip4
STR8−chip4
B2AA2−chip3
22027AF−chip3
103FR−chip4
4HU3AA−chip3
5000
4000
3000
2000
1000
0

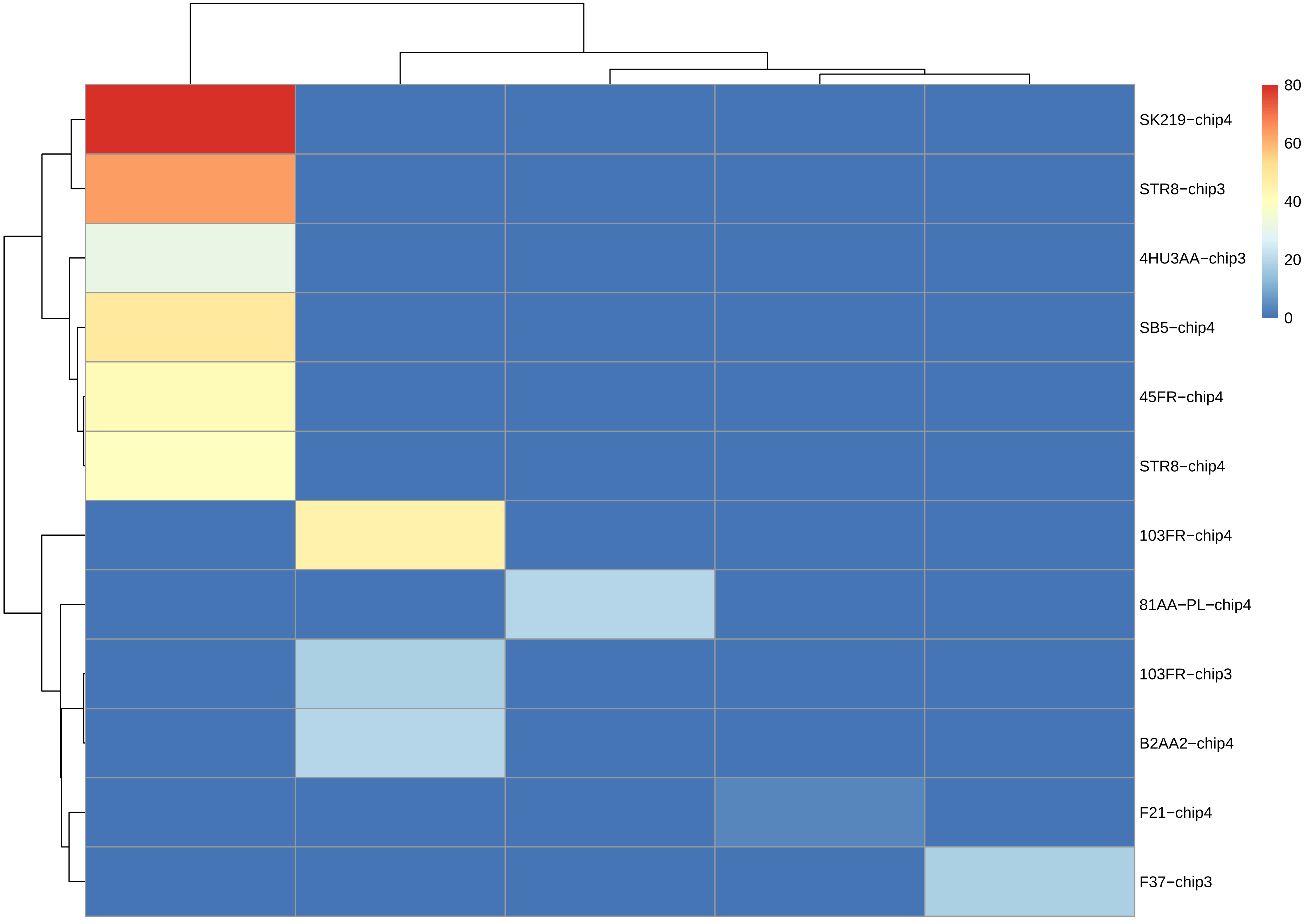
SK219−chip4
STR8−chip3
4HU3AA−chip3
SB5−chip4
45FR−chip4
STR8−chip4
103FR−chip4
81AA−PL−chip4
103FR−chip3
B2AA2−chip4
F21−chip4
F37−chip3
80
60
40
20
0

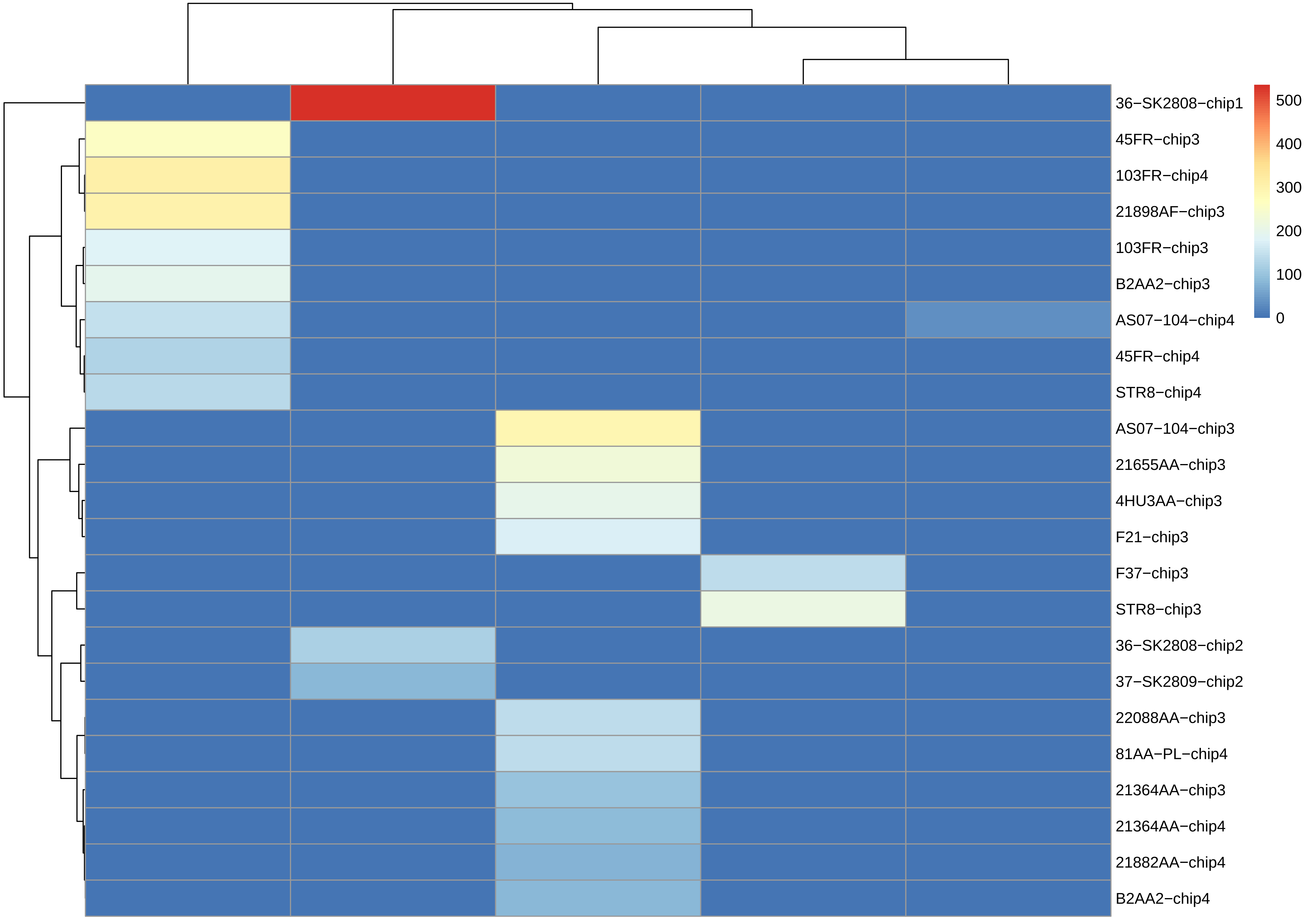

36−SK2808−chip1
45FR−chip3
103FR−chip4
21898AF−chip3
103FR−chip3
B2AA2−chip3
AS07−104−chip4
45FR−chip4
STR8−chip4
AS07−104−chip3
21655AA−chip3
4HU3AA−chip3
F21−chip3
F37−chip3
STR8−chip3
36−SK2808−chip2
37−SK2809−chip2
22088AA−chip3
81AA−PL−chip4
21364AA−chip3
21364AA−chip4
21882AA−chip4
B2AA2−chip4
500
400
300
200
100
0

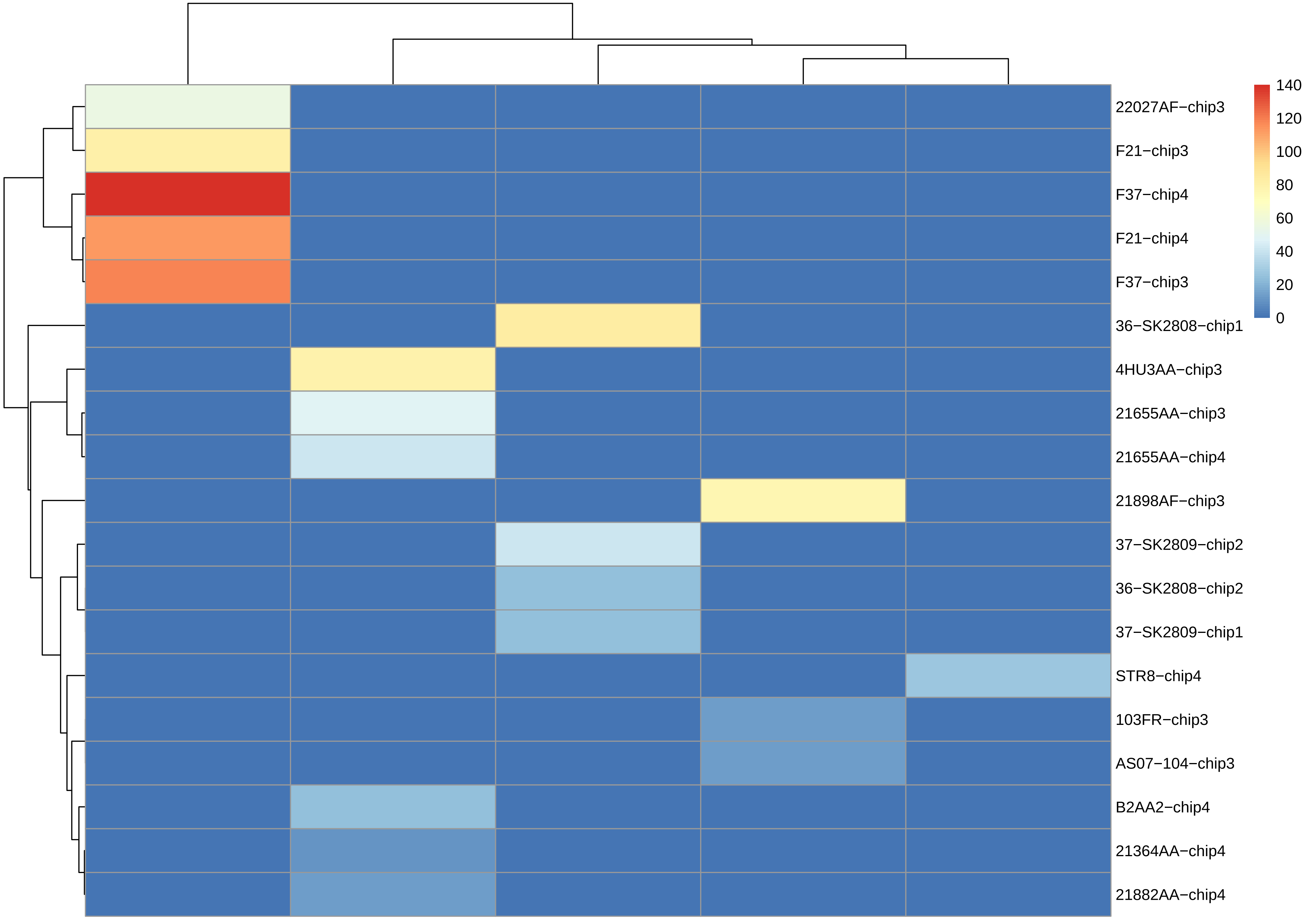

22027AF−chip3
F21−chip3
F37−chip4
F21−chip4
F37−chip3
36−SK2808−chip1
4HU3AA−chip3
21655AA−chip3
21655AA−chip4
21898AF−chip3
37−SK2809−chip2
36−SK2808−chip2
37−SK2809−chip1
STR8−chip4
103FR−chip3
AS07−104−chip3
B2AA2−chip4
21364AA−chip4
21882AA−chip4
140
120
100
80
60
40
20
0

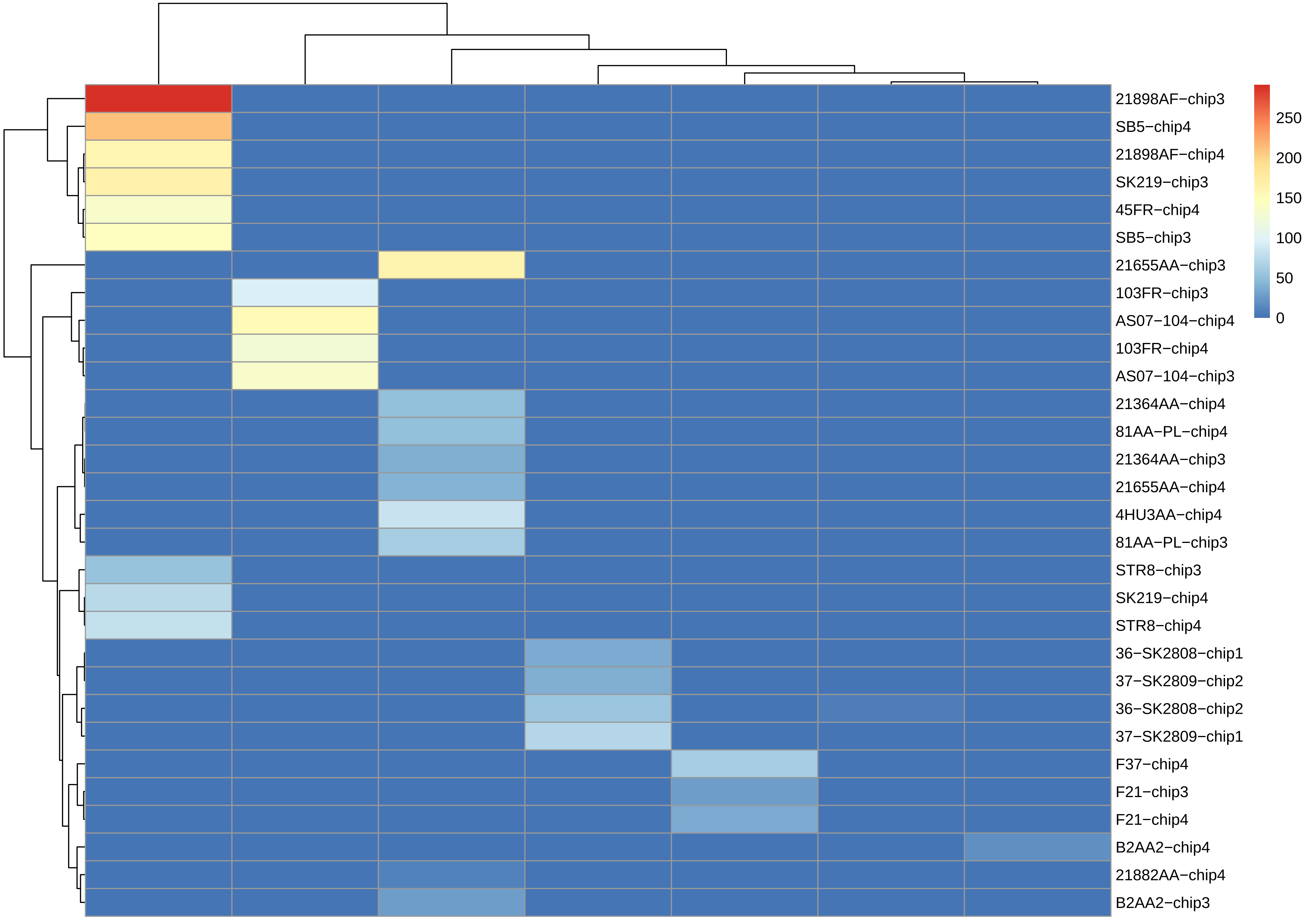

21898AF−chip3
SB5−chip4
21898AF−chip4
SK219−chip3
45FR−chip4
SB5−chip3
21655AA−chip3
103FR−chip3
AS07−104−chip4
103FR−chip4
AS07−104−chip3
21364AA−chip4
81AA−PL−chip4
21364AA−chip3
21655AA−chip4
4HU3AA−chip4
81AA−PL−chip3
STR8−chip3
SK219−chip4
STR8−chip4
36−SK2808−chip1
37−SK2809−chip2
36−SK2808−chip2
37−SK2809−chip1
F37−chip4
F21−chip3
F21−chip4
B2AA2−chip4
21882AA−chip4
B2AA2−chip3
250
200
150
100
50
0

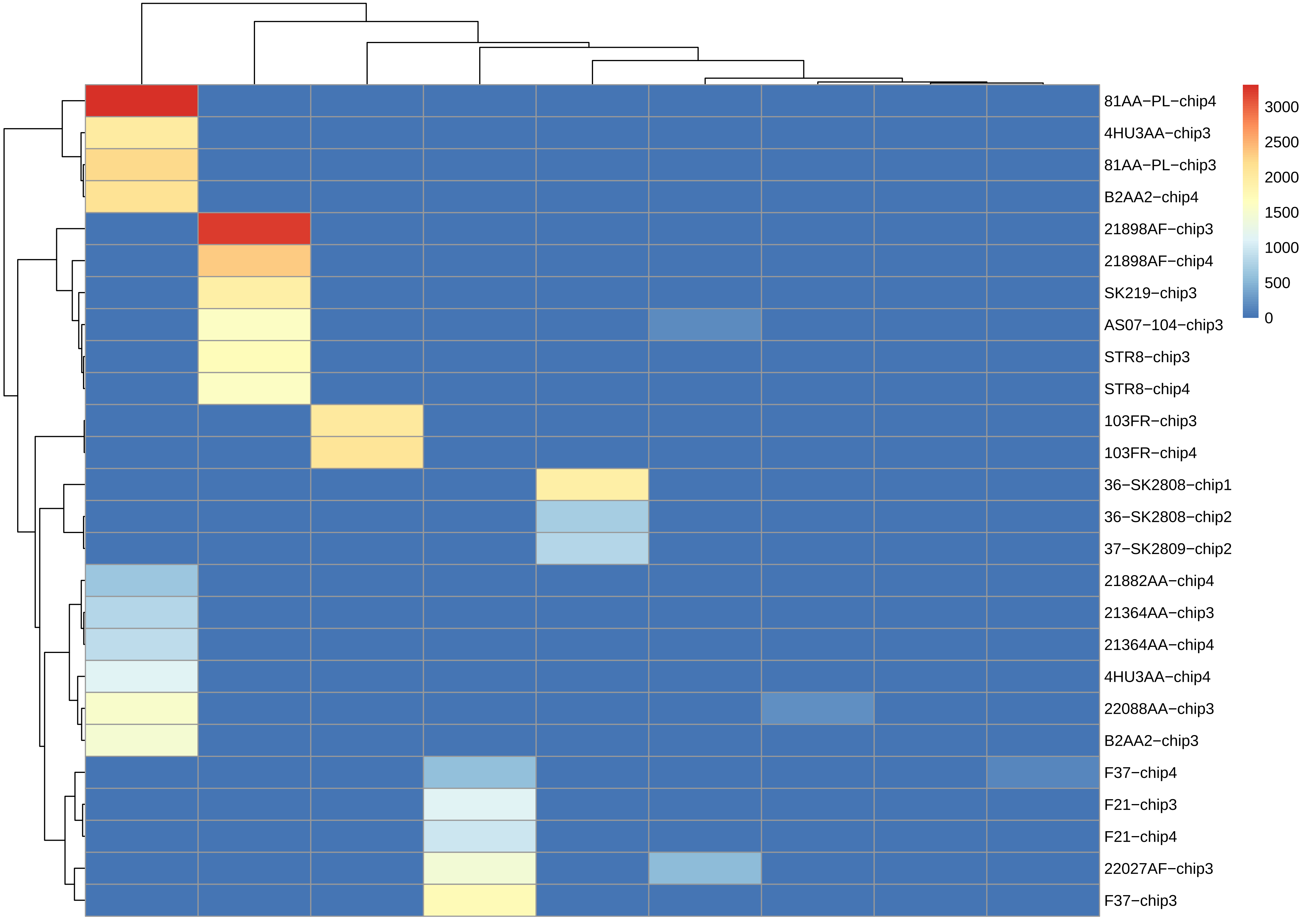

81AA−PL−chip4
4HU3AA−chip3
81AA−PL−chip3
B2AA2−chip4
21898AF−chip3
21898AF−chip4
SK219−chip3
AS07−104−chip3
STR8−chip3
STR8−chip4
103FR−chip3
103FR−chip4
36−SK2808−chip1
36−SK2808−chip2
37−SK2809−chip2
21882AA−chip4
21364AA−chip3
21364AA−chip4
4HU3AA−chip4
22088AA−chip3
B2AA2−chip3
F37−chip4
F21−chip3
F21−chip4
22027AF−chip3
F37−chip3
3000
2500
2000
1500
1000
500
0

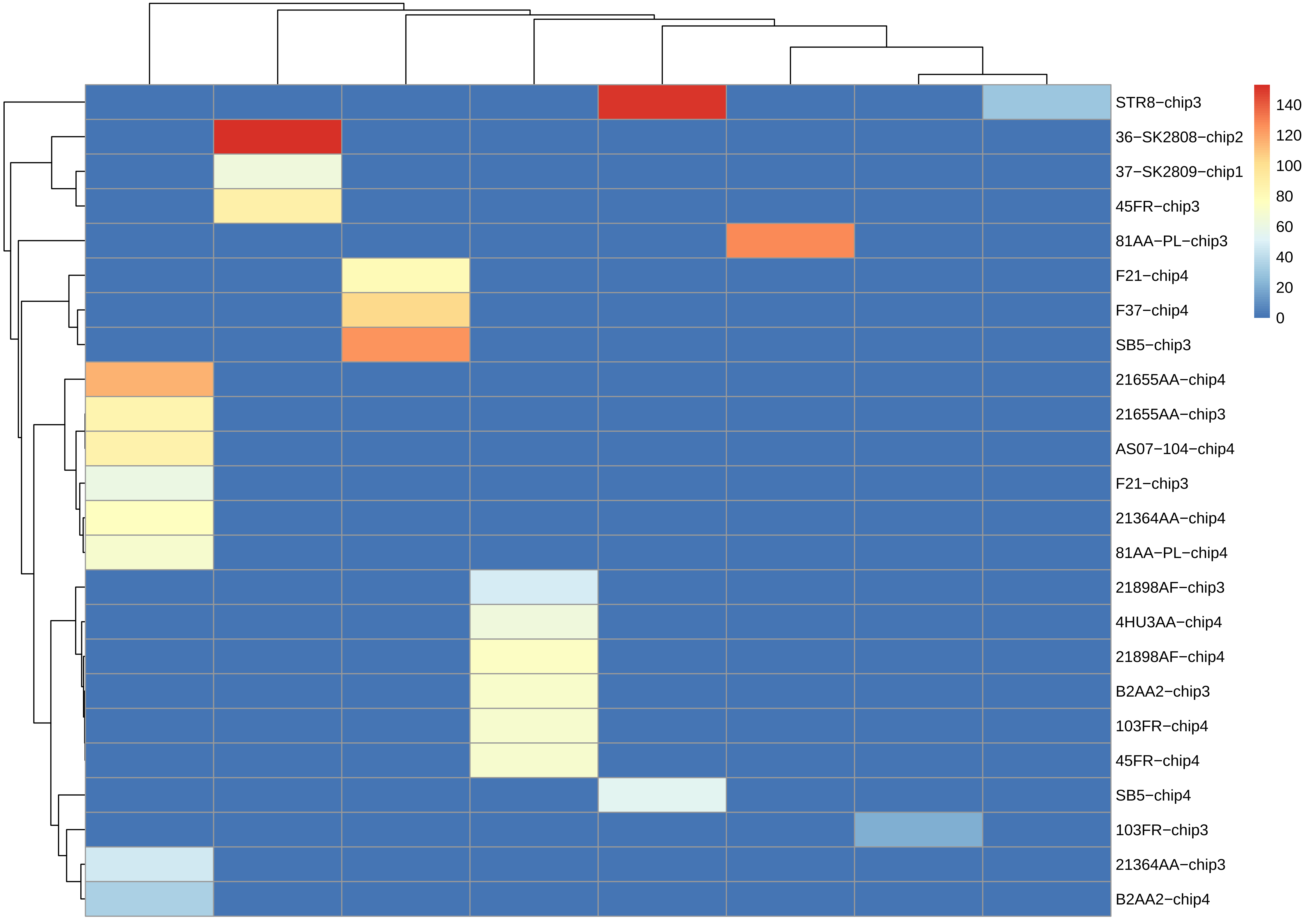
STR8−chip3
36−SK2808−chip2
37−SK2809−chip1
45FR−chip3
81AA−PL−chip3
F21−chip4
F37−chip4
SB5−chip3
21655AA−chip4
21655AA−chip3
AS07−104−chip4
F21−chip3
21364AA−chip4
81AA−PL−chip4
21898AF−chip3
4HU3AA−chip4
21898AF−chip4
B2AA2−chip3
103FR−chip4
45FR−chip4
SB5−chip4
103FR−chip3
21364AA−chip3
B2AA2−chip4
140
120
100
80
60
40
20
0

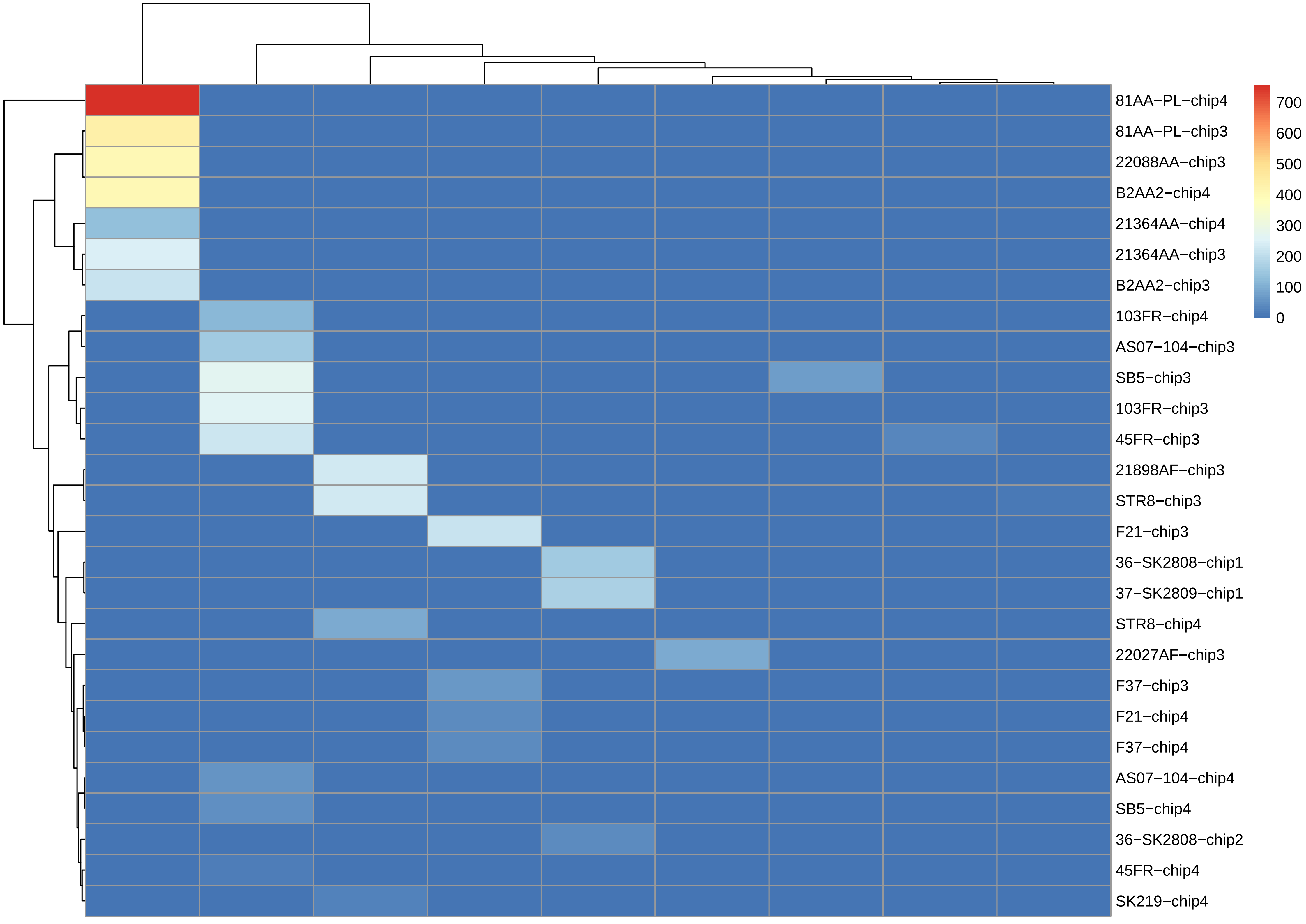

81AA−PL−chip4
81AA−PL−chip3
22088AA−chip3
B2AA2−chip4
21364AA−chip4
21364AA−chip3
B2AA2−chip3
103FR−chip4
AS07−104−chip3
SB5−chip3
103FR−chip3
45FR−chip3
21898AF−chip3
STR8−chip3
F21−chip3
36−SK2808−chip1
37−SK2809−chip1
STR8−chip4
22027AF−chip3
F37−chip3
F21−chip4
F37−chip4
AS07−104−chip4
SB5−chip4
36−SK2808−chip2
45FR−chip4
SK219−chip4
700
600
500
400
300
200
100
0

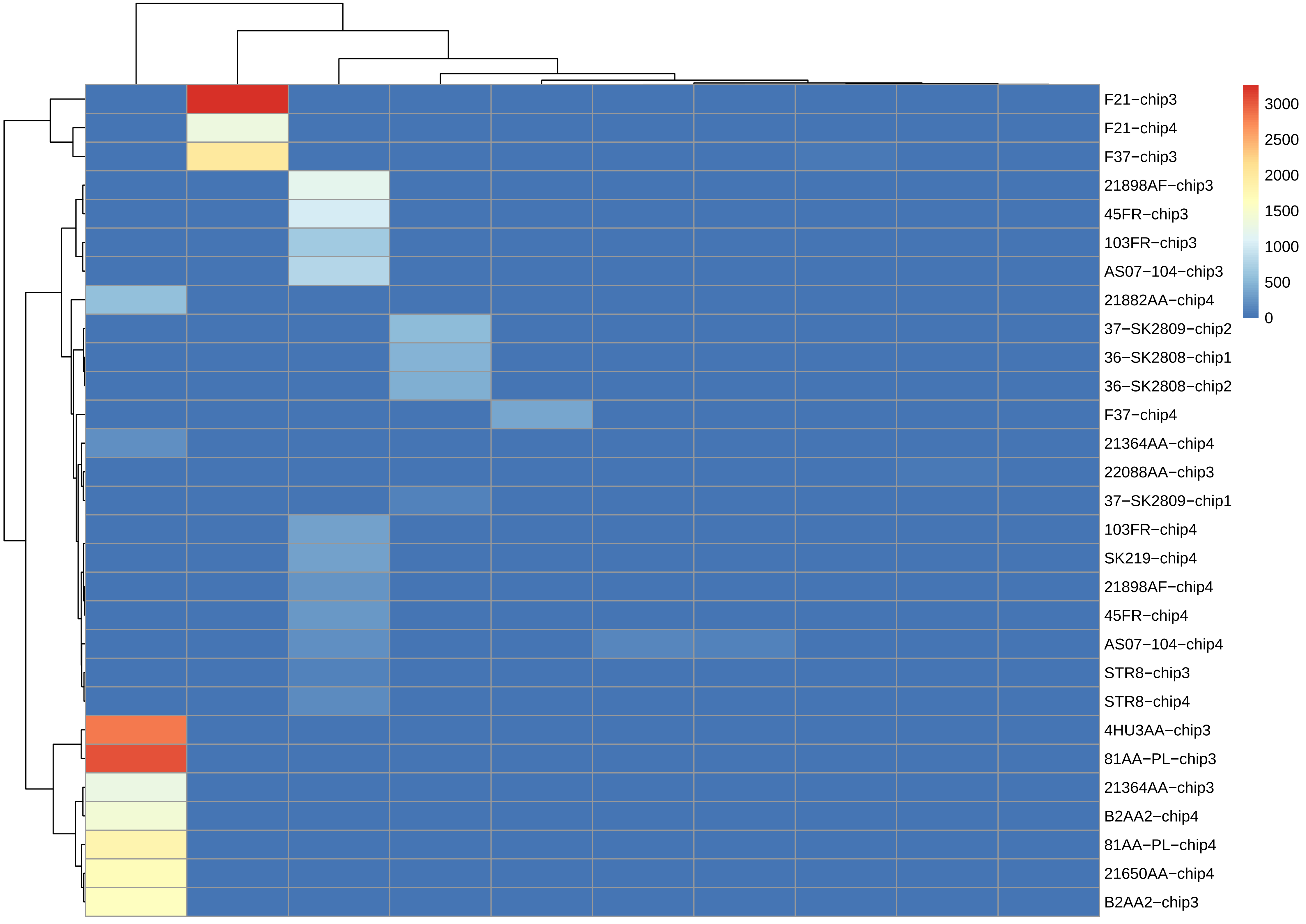
F21−chip3
F21−chip4
F37−chip3
21898AF−chip3
45FR−chip3
103FR−chip3
AS07−104−chip3
21882AA−chip4
37−SK2809−chip2
36−SK2808−chip1
36−SK2808−chip2
F37−chip4
21364AA−chip4
22088AA−chip3
37−SK2809−chip1
103FR−chip4
SK219−chip4
21898AF−chip4
45FR−chip4
AS07−104−chip4
STR8−chip3
STR8−chip4
4HU3AA−chip3
81AA−PL−chip3
21364AA−chip3
B2AA2−chip4
81AA−PL−chip4
21650AA−chip4
B2AA2−chip3
3000
2500
2000
1500
1000
500
0

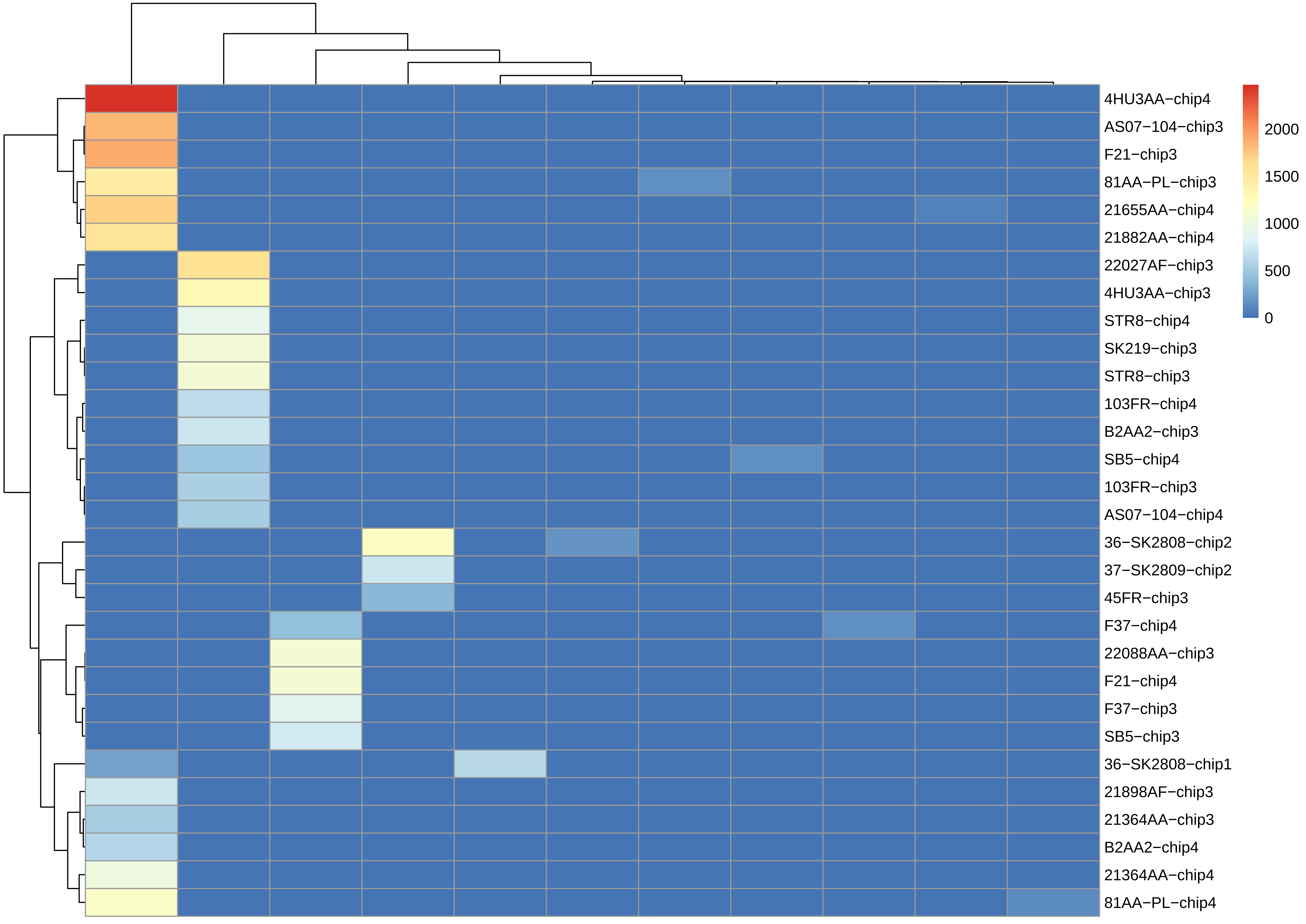
4HU3AA−chip4
AS07−104−chip3
F21−chip3
81AA−PL−chip3
21655AA−chip4
21882AA−chip4
22027AF−chip3
4HU3AA−chip3
STR8−chip4
SK219−chip3
STR8−chip3
103FR−chip4
B2AA2−chip3
SB5−chip4
103FR−chip3
AS07−104−chip4
36−SK2808−chip2
37−SK2809−chip2
45FR−chip3
F37−chip4
22088AA−chip3
F21−chip4
F37−chip3
SB5−chip3
36−SK2808−chip1
21898AF−chip3
21364AA−chip3
B2AA2−chip4
21364AA−chip4
81AA−PL−chip4
2000
1500
1000
500
0

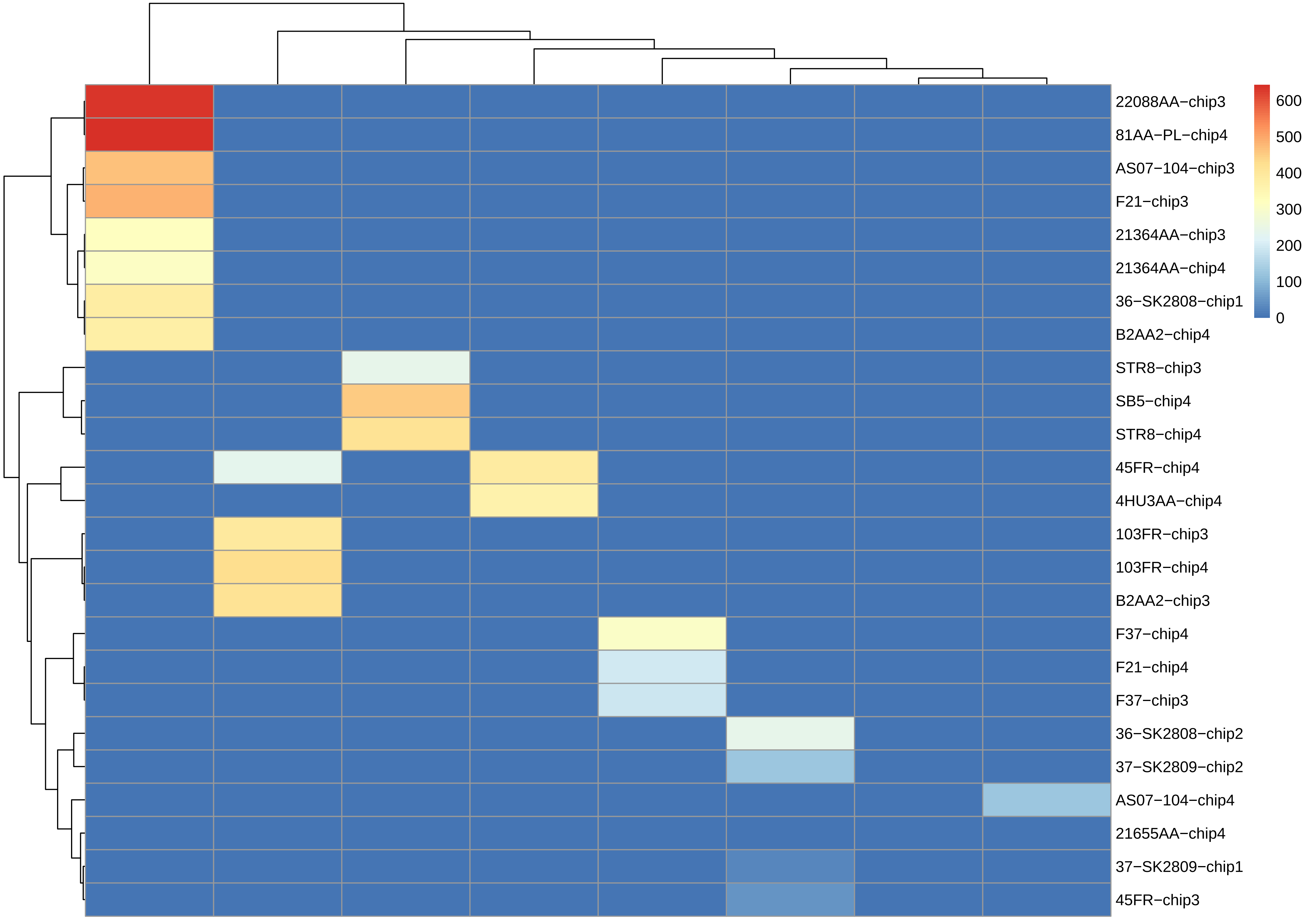
22088AA−chip3
81AA−PL−chip4
AS07−104−chip3
F21−chip3
21364AA−chip3
21364AA−chip4
36−SK2808−chip1
B2AA2−chip4
STR8−chip3
SB5−chip4
STR8−chip4
45FR−chip4
4HU3AA−chip4
103FR−chip3
103FR−chip4
B2AA2−chip3
F37−chip4
F21−chip4
F37−chip3
36−SK2808−chip2
37−SK2809−chip2
AS07−104−chip4
21655AA−chip4
37−SK2809−chip1
45FR−chip3
600
500
400
300
200
100
0

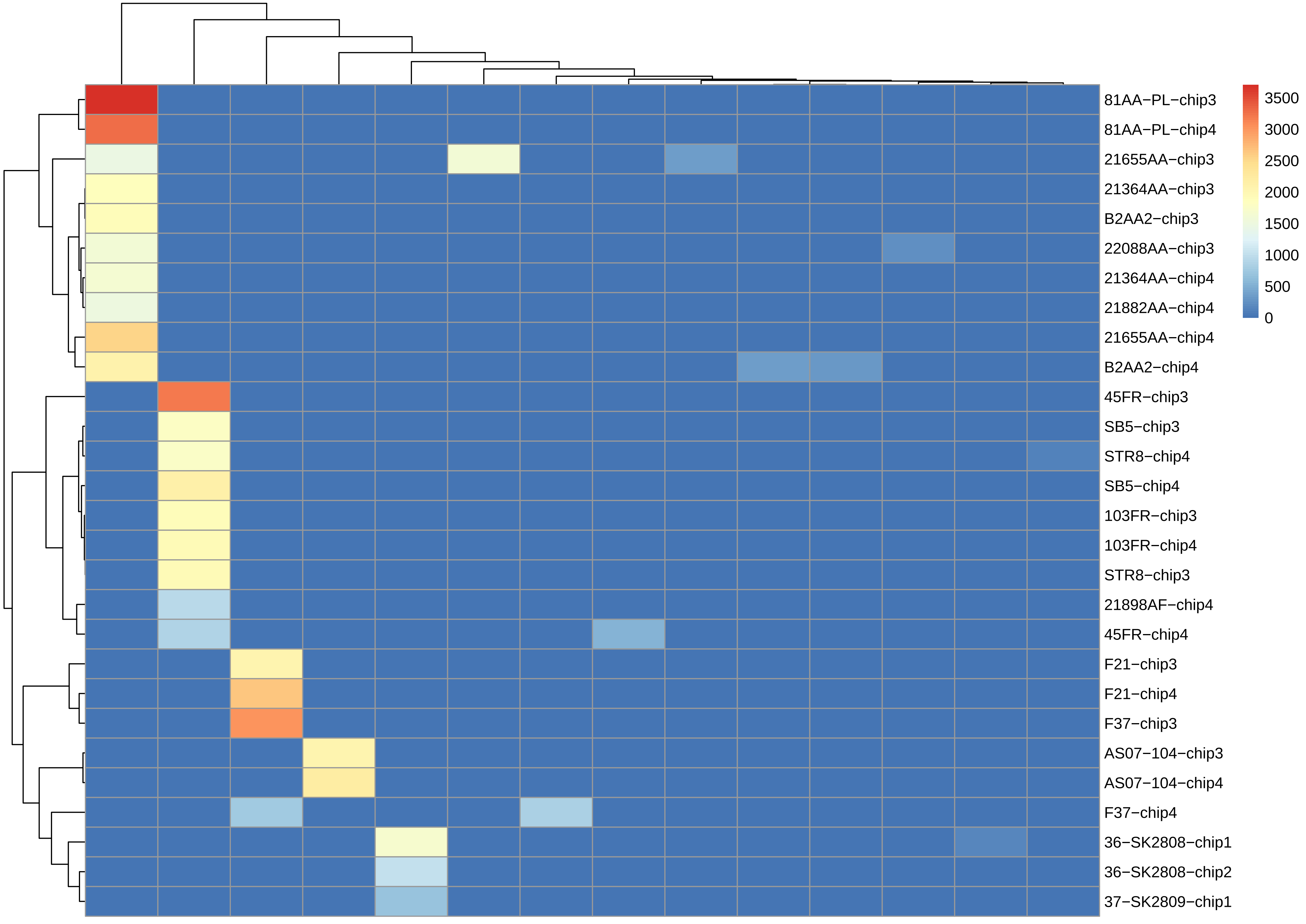

81AA−PL−chip3
81AA−PL−chip4
21655AA−chip3
21364AA−chip3
B2AA2−chip3
22088AA−chip3
21364AA−chip4
21882AA−chip4
21655AA−chip4
B2AA2−chip4
45FR−chip3
SB5−chip3
STR8−chip4
SB5−chip4
103FR−chip3
103FR−chip4
STR8−chip3
21898AF−chip4
45FR−chip4
F21−chip3
F21−chip4
F37−chip3
AS07−104−chip3
AS07−104−chip4
F37−chip4
36−SK2808−chip1
36−SK2808−chip2
37−SK2809−chip1
3500
3000
2500
2000
1500
1000
500
0

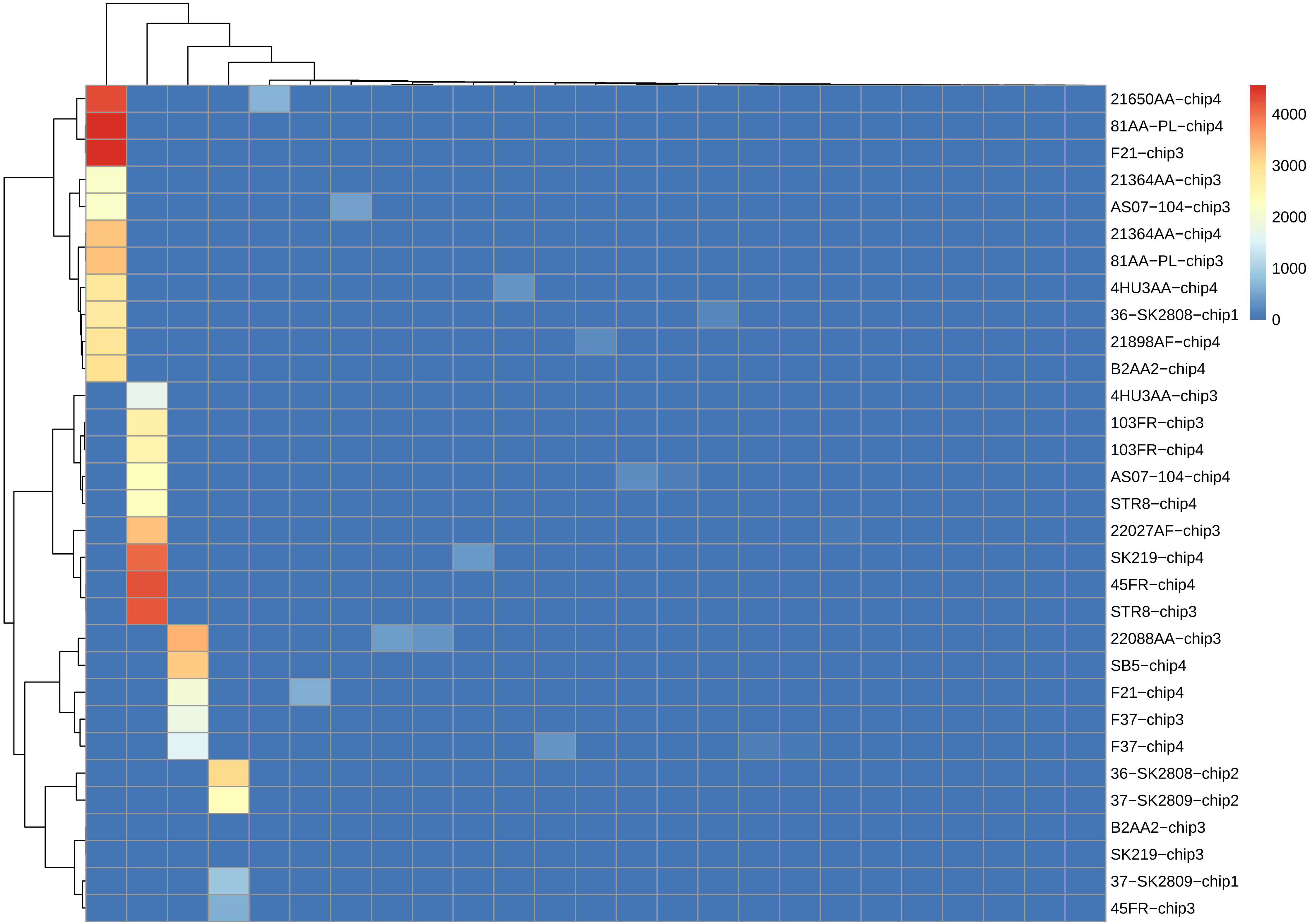
21650AA−chip4
81AA−PL−chip4
F21−chip3
21364AA−chip3
AS07−104−chip3
21364AA−chip4
81AA−PL−chip3
4HU3AA−chip4
36−SK2808−chip1
21898AF−chip4
B2AA2−chip4
4HU3AA−chip3
103FR−chip3
103FR−chip4
AS07−104−chip4
STR8−chip4
22027AF−chip3
SK219−chip4
45FR−chip4
STR8−chip3
22088AA−chip3
SB5−chip4
F21−chip4
F37−chip3
F37−chip4
36−SK2808−chip2
37−SK2809−chip2
B2AA2−chip3
SK219−chip3
37−SK2809−chip1
45FR−chip3
4000
3000
2000
1000
0

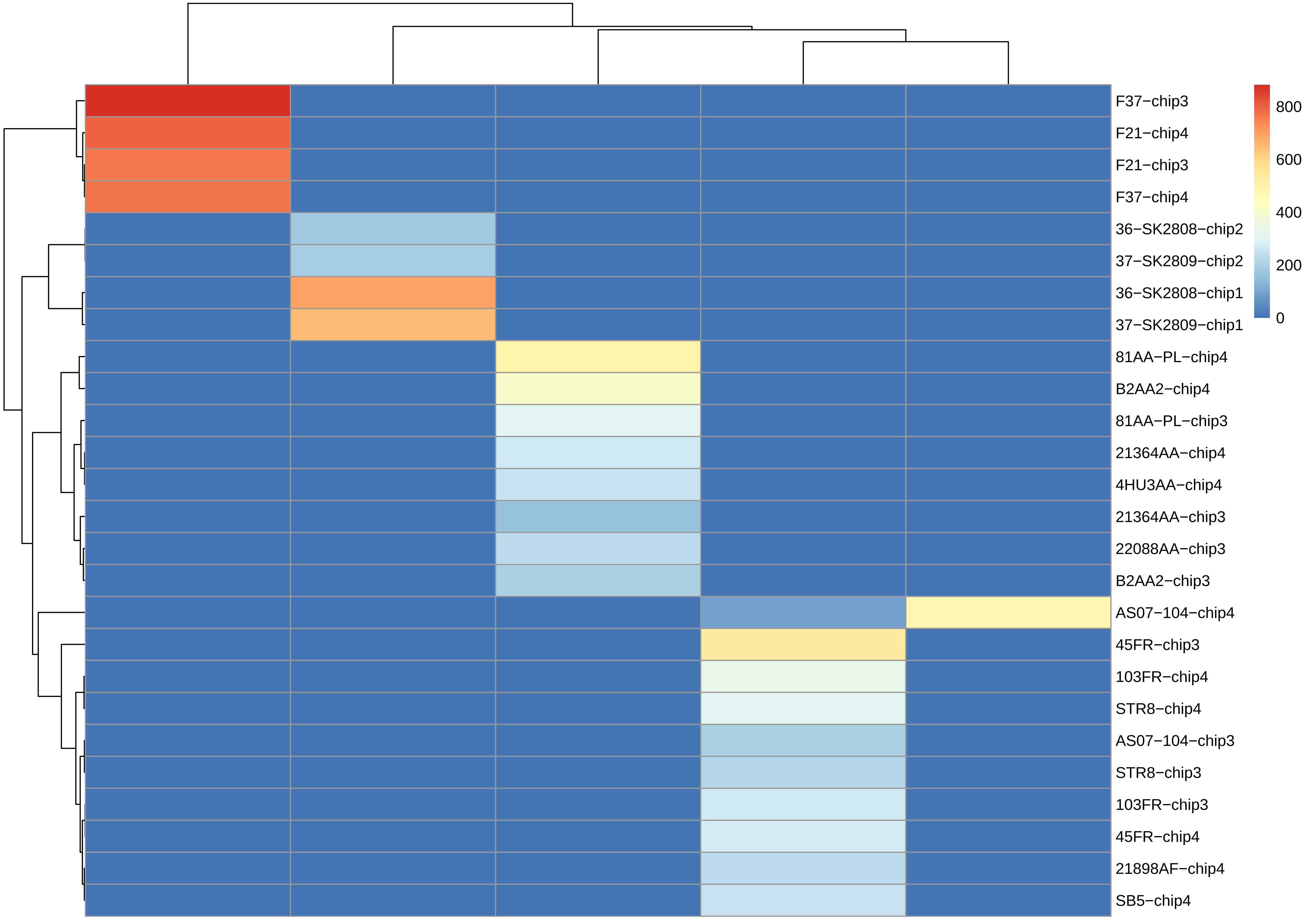
800
600
400
200
0
F37−chip3
F21−chip4
F21−chip3
F37−chip4
36−SK2808−chip2
37−SK2809−chip2
36−SK2808−chip1
37−SK2809−chip1
81AA−PL−chip4
B2AA2−chip4
81AA−PL−chip3
21364AA−chip4
4HU3AA−chip4
21364AA−chip3
22088AA−chip3
B2AA2−chip3
AS07−104−chip4
45FR−chip3
103FR−chip4
STR8−chip4
AS07−104−chip3
STR8−chip3
103FR−chip3
45FR−chip4
21898AF−chip4
SB5−chip4

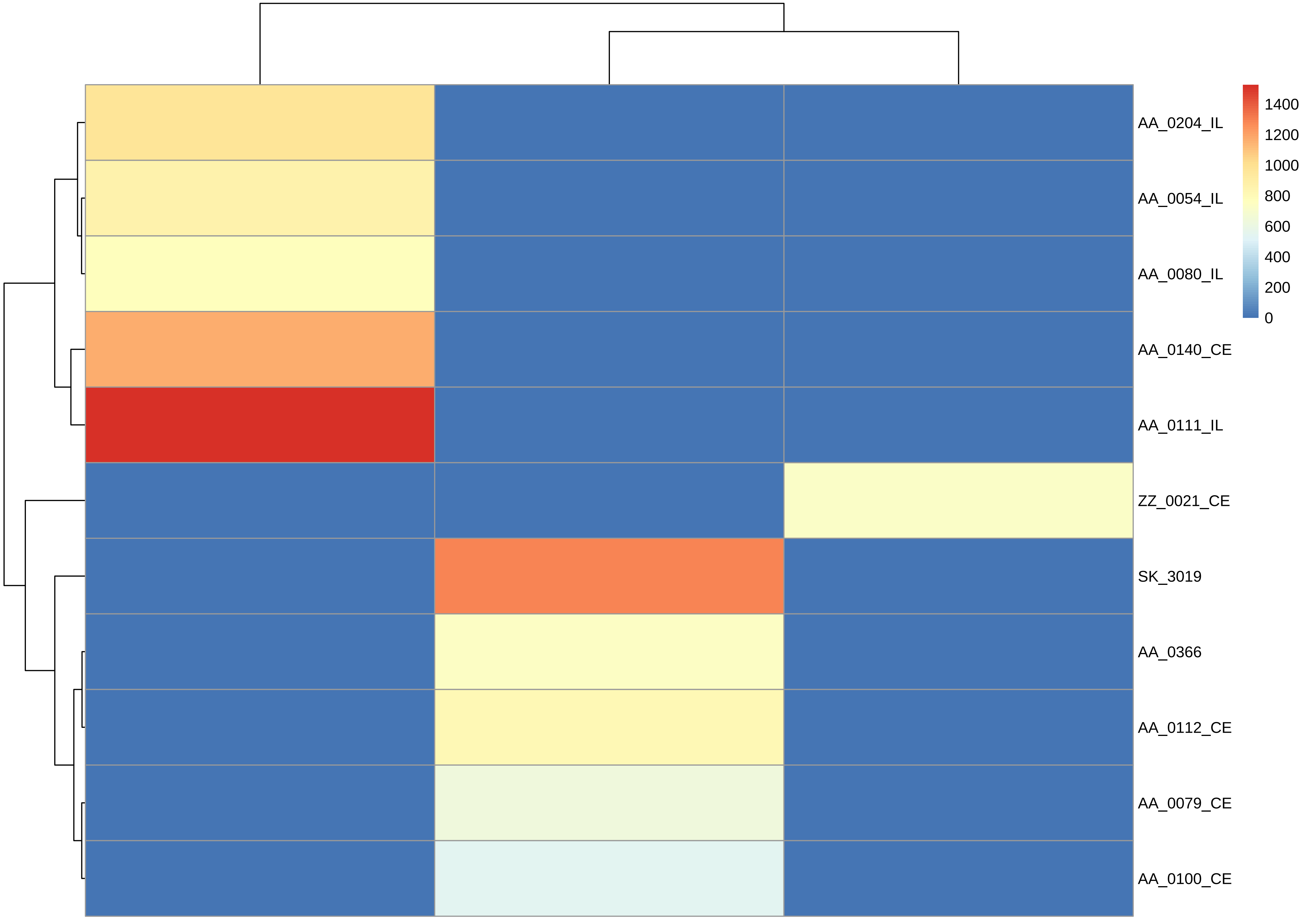
1400
1200
1000
800
600
400
200
0
AA_0204_IL
AA_0054_IL
AA_0080_IL
AA_0140_CE
AA_0111_IL
ZZ_0021_CE
SK_3019
AA_0366
AA_0112_CE
AA_0079_CE
AA_0100_CE

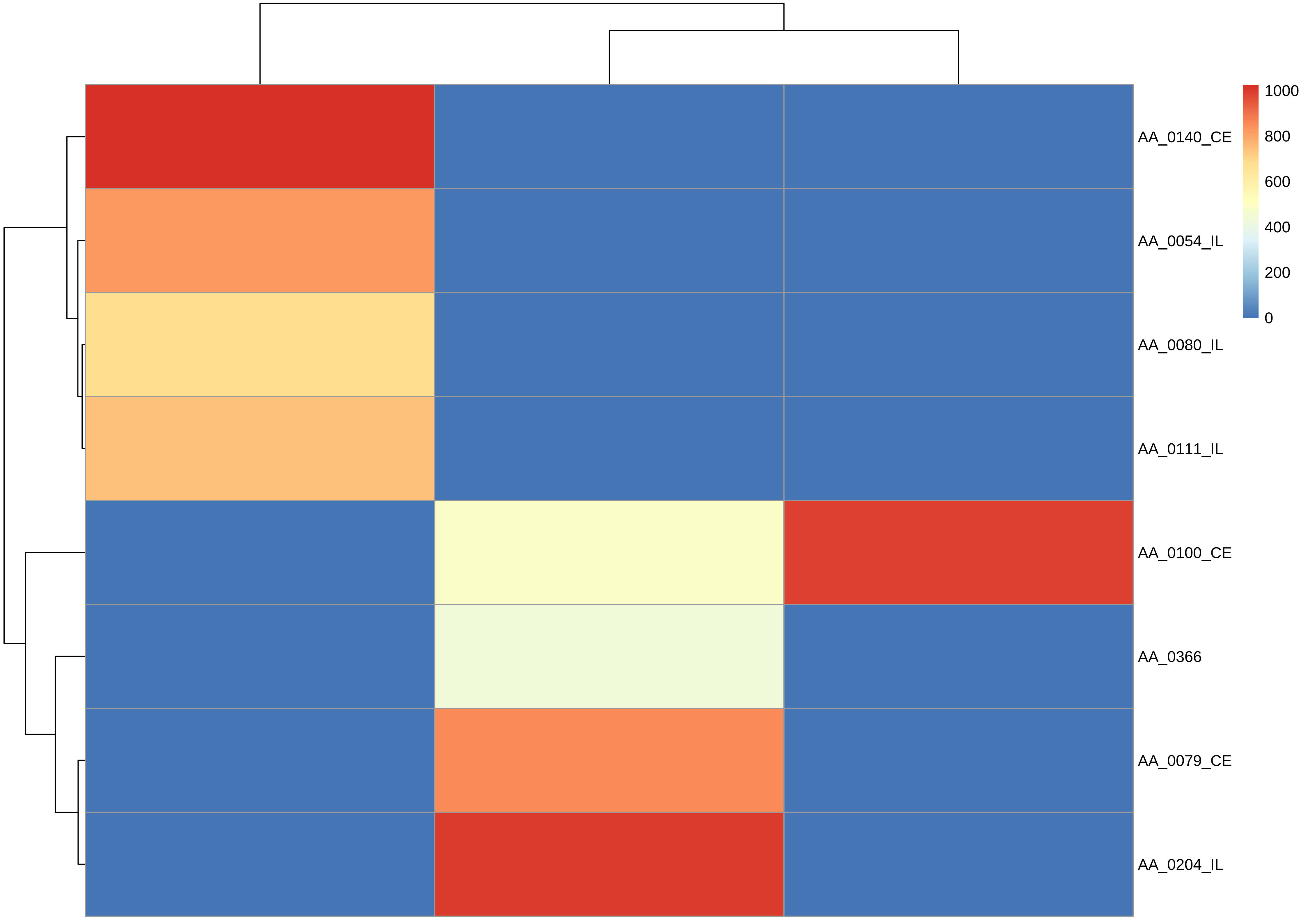

1000
800
600
400
200
0
AA_0140_CE
AA_0054_IL
AA_0080_IL
AA_0111_IL
AA_0100_CE
AA_0366
AA_0079_CE
AA_0204_IL

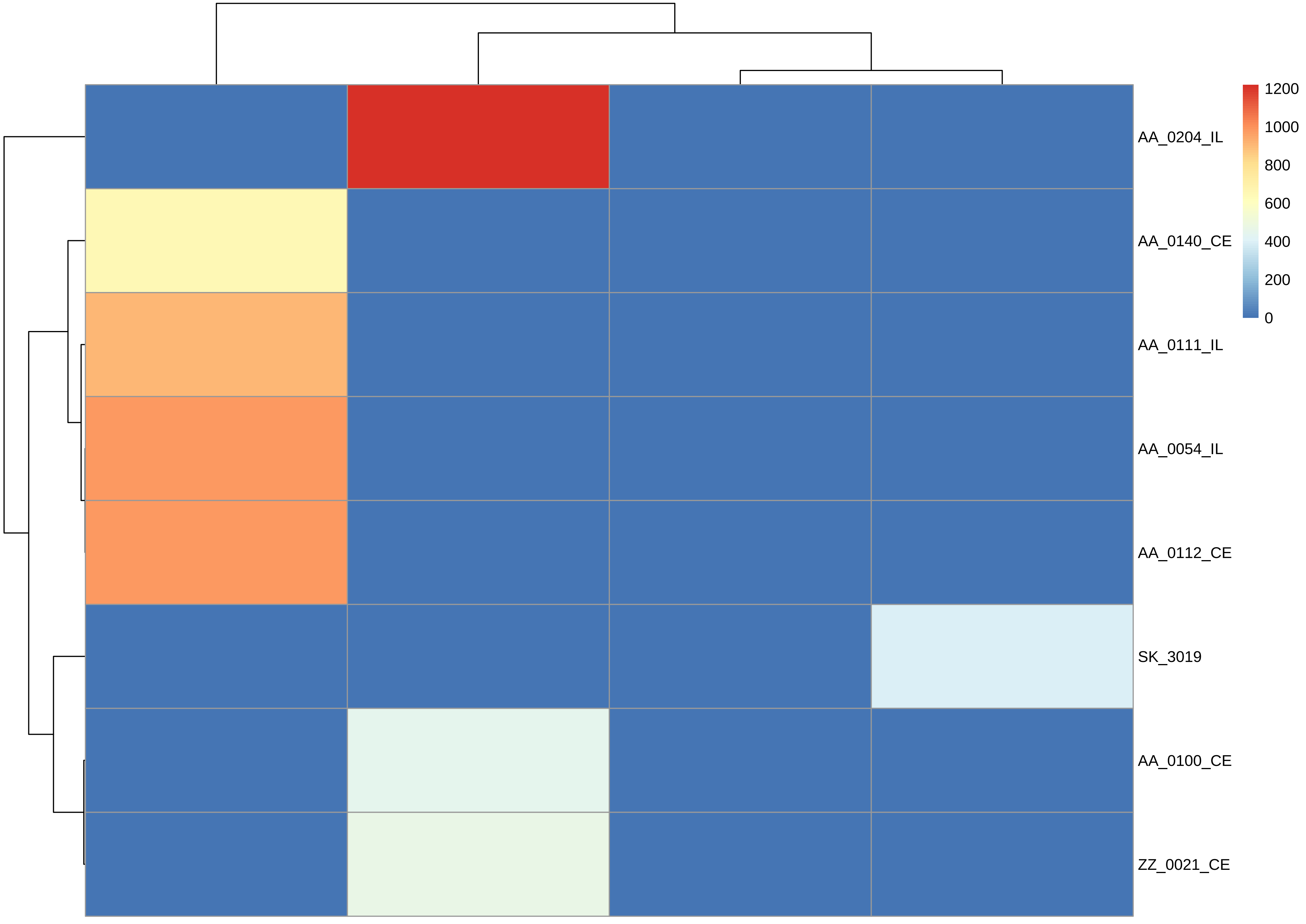
1200
1000
800
600
400
200
0
AA_0204_IL
AA_0140_CE
AA_0111_IL
AA_0054_IL
AA_0112_CE
SK_3019
AA_0100_CE
ZZ_0021_CE

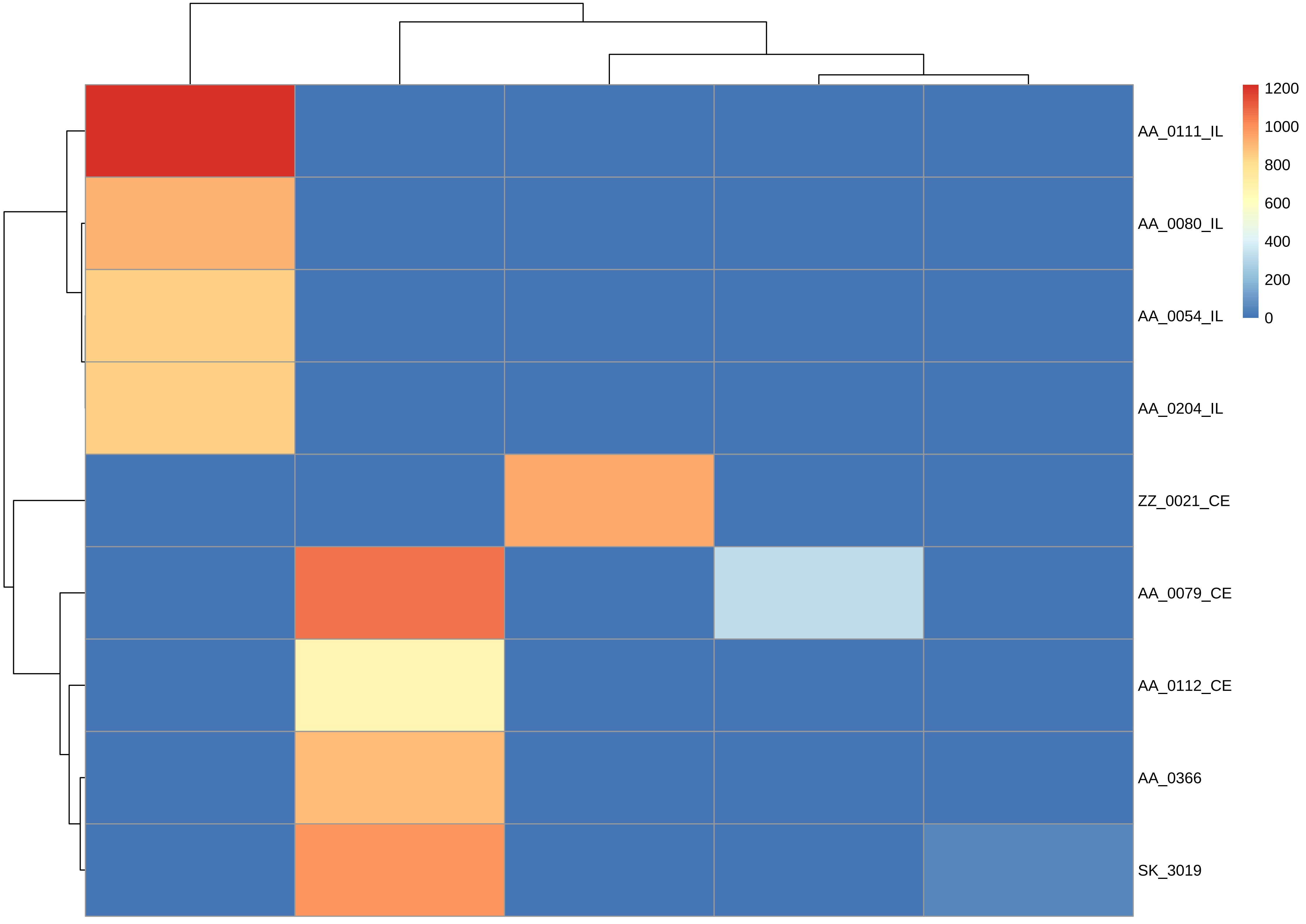
AA_0111_IL
AA_0080_IL
AA_0054_IL
AA_0204_IL
ZZ_0021_CE
AA_0079_CE
AA_0112_CE
AA_0366
SK_3019
1200
1000
800
600
400
200
0

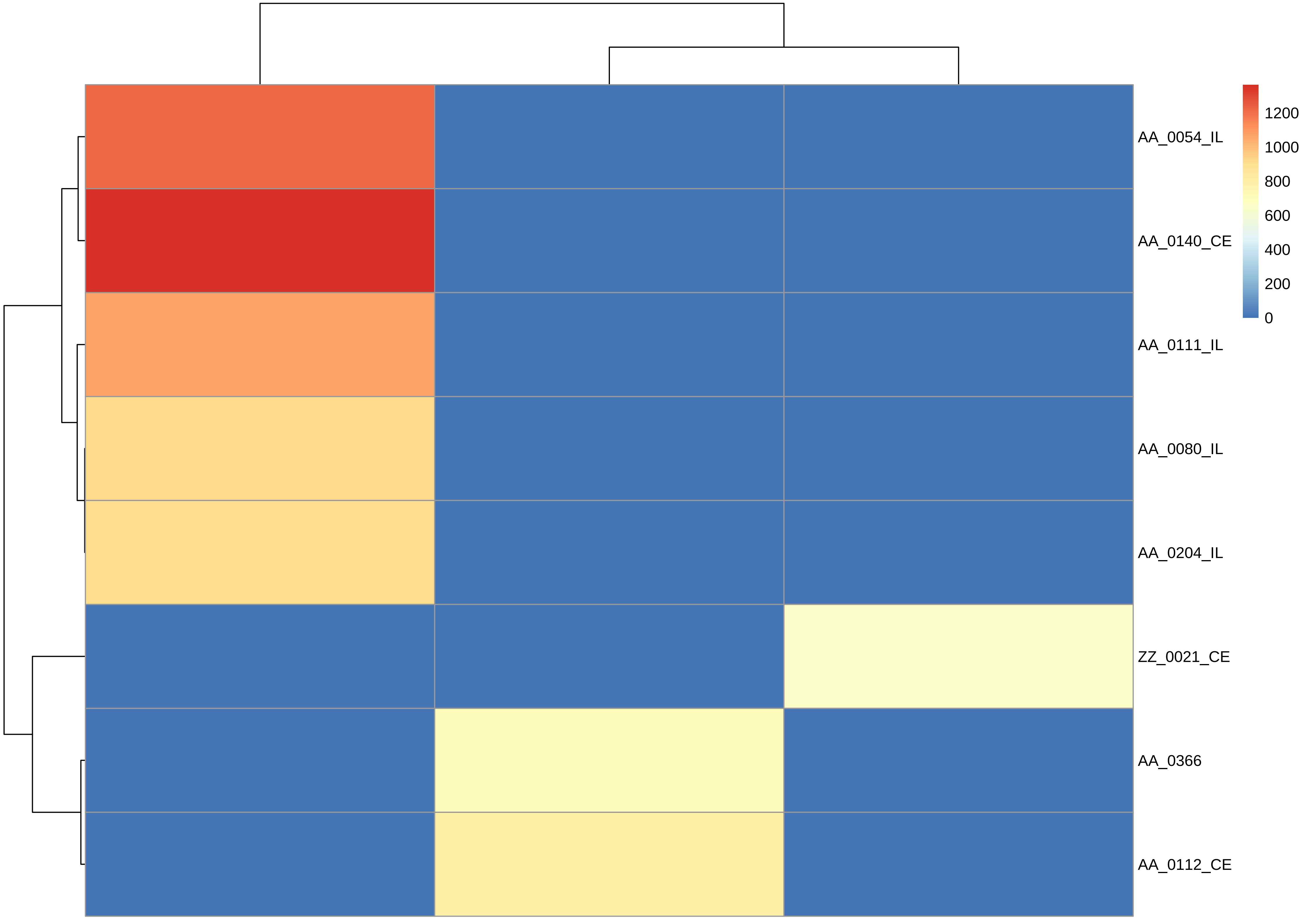

AA_0054_IL
AA_0140_CE
AA_0111_IL
AA_0080_IL
AA_0204_IL
ZZ_0021_CE
AA_0366
AA_0112_CE
1200
1000
800
600
400
200
0

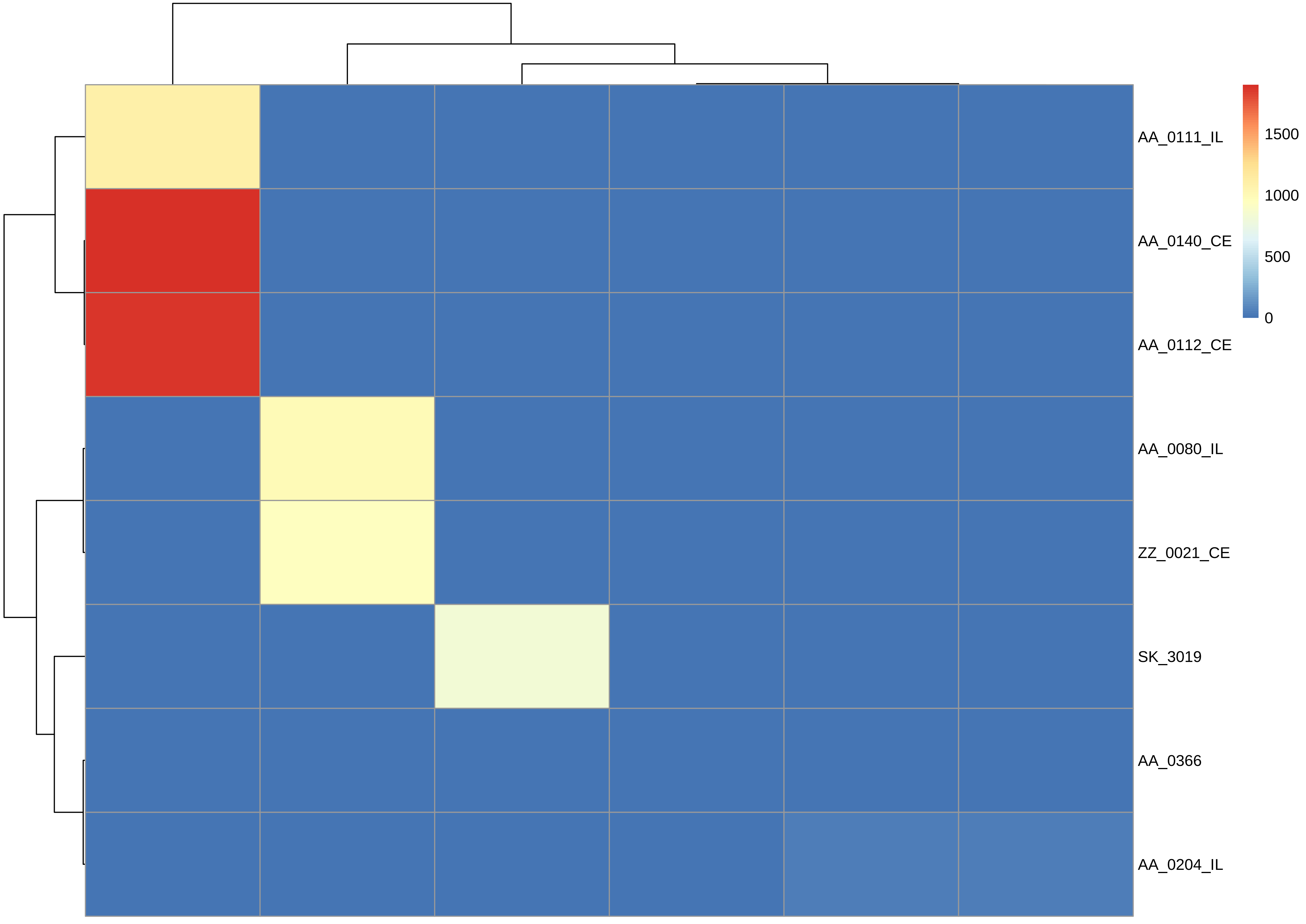

AA_0111_IL
AA_0140_CE
AA_0112_CE
AA_0080_IL
ZZ_0021_CE
SK_3019
AA_0366
AA_0204_IL
1500
1000
500
0

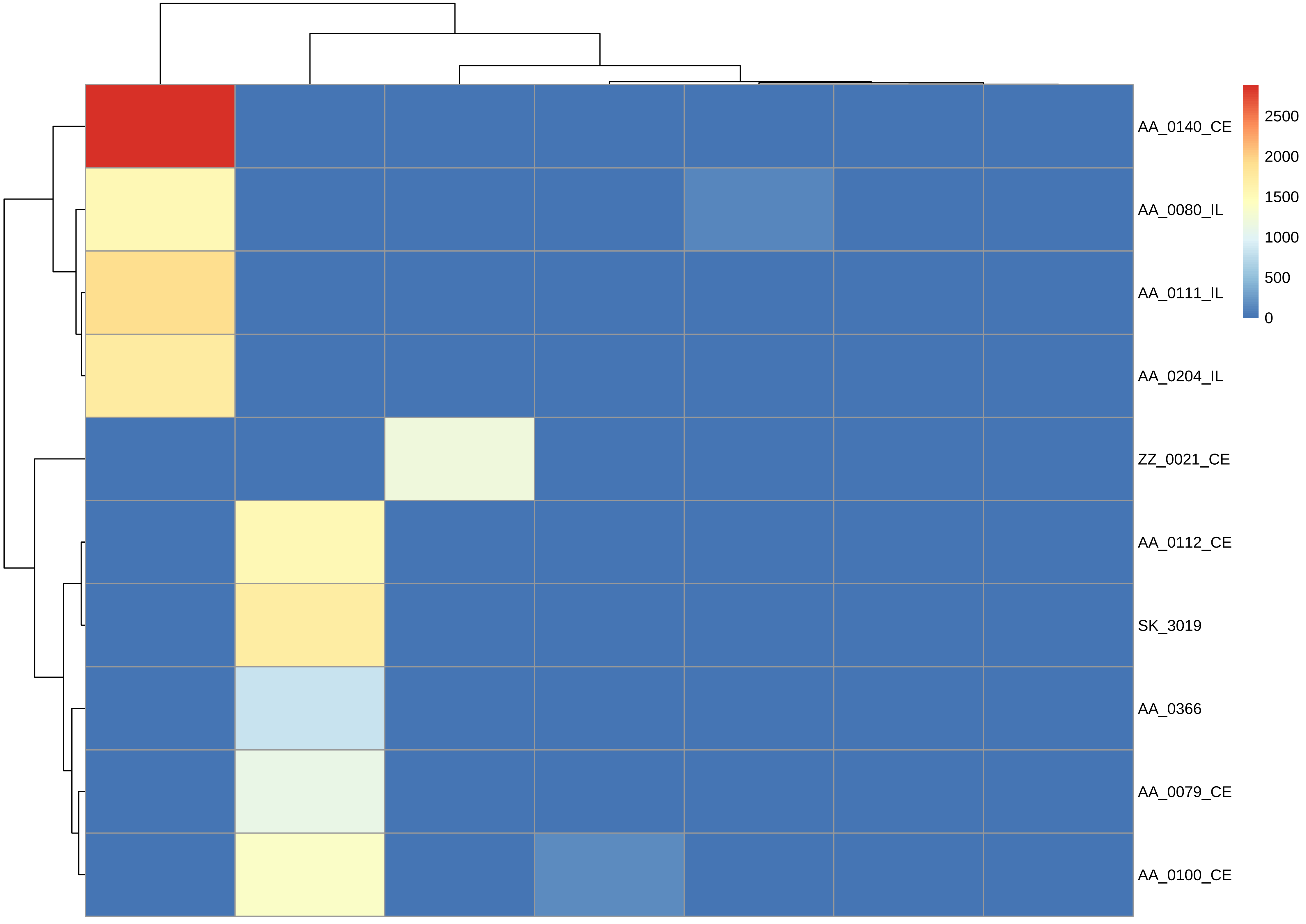

AA_0140_CE
AA_0080_IL
AA_0111_IL
AA_0204_IL
ZZ_0021_CE
AA_0112_CE
SK_3019
AA_0366
AA_0079_CE
AA_0100_CE
2500
2000
1500
1000
500
0

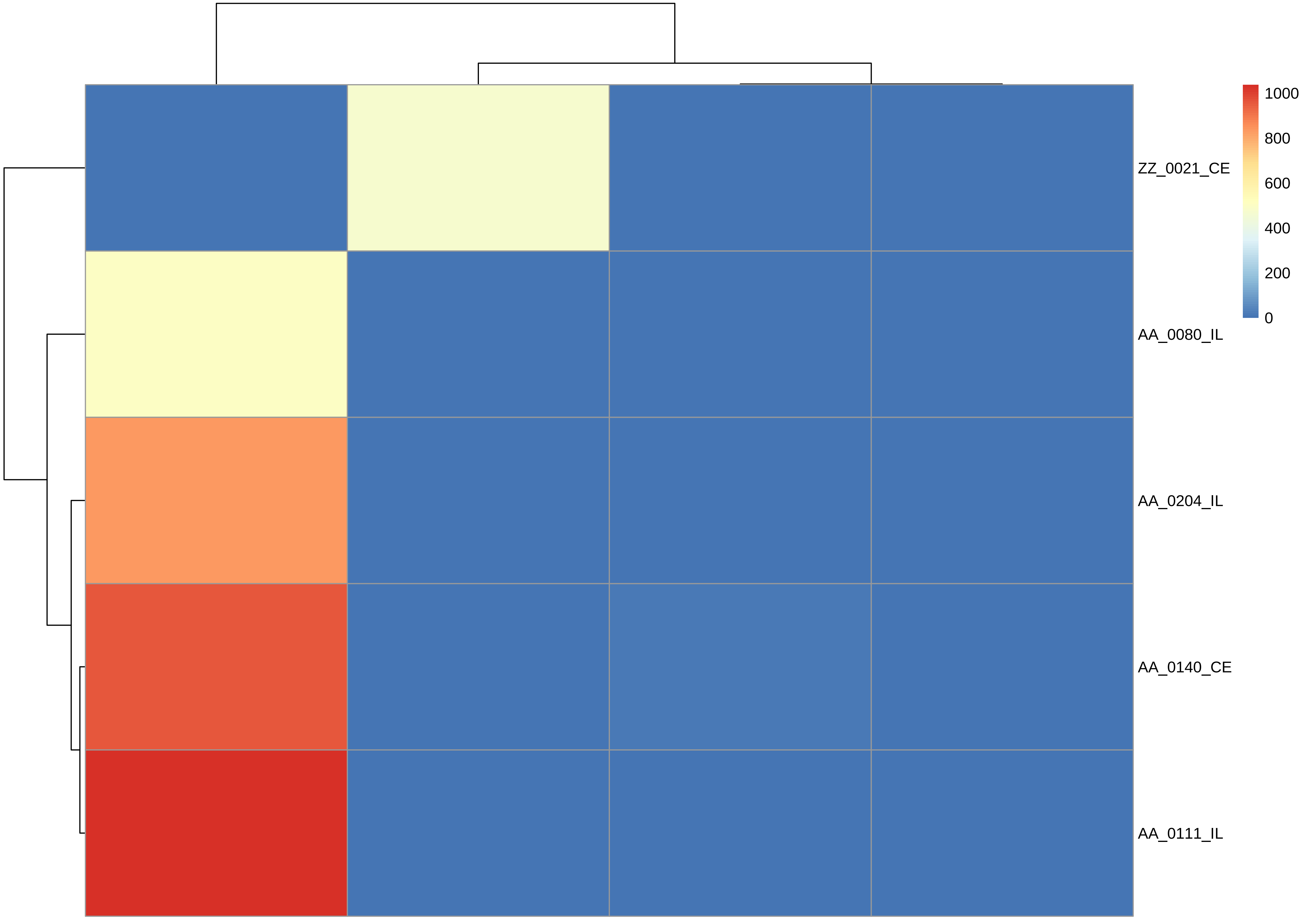

ZZ_0021_CE
AA_0080_IL
AA_0204_IL
AA_0140_CE
AA_0111_IL
1000
800
600
400
200
0

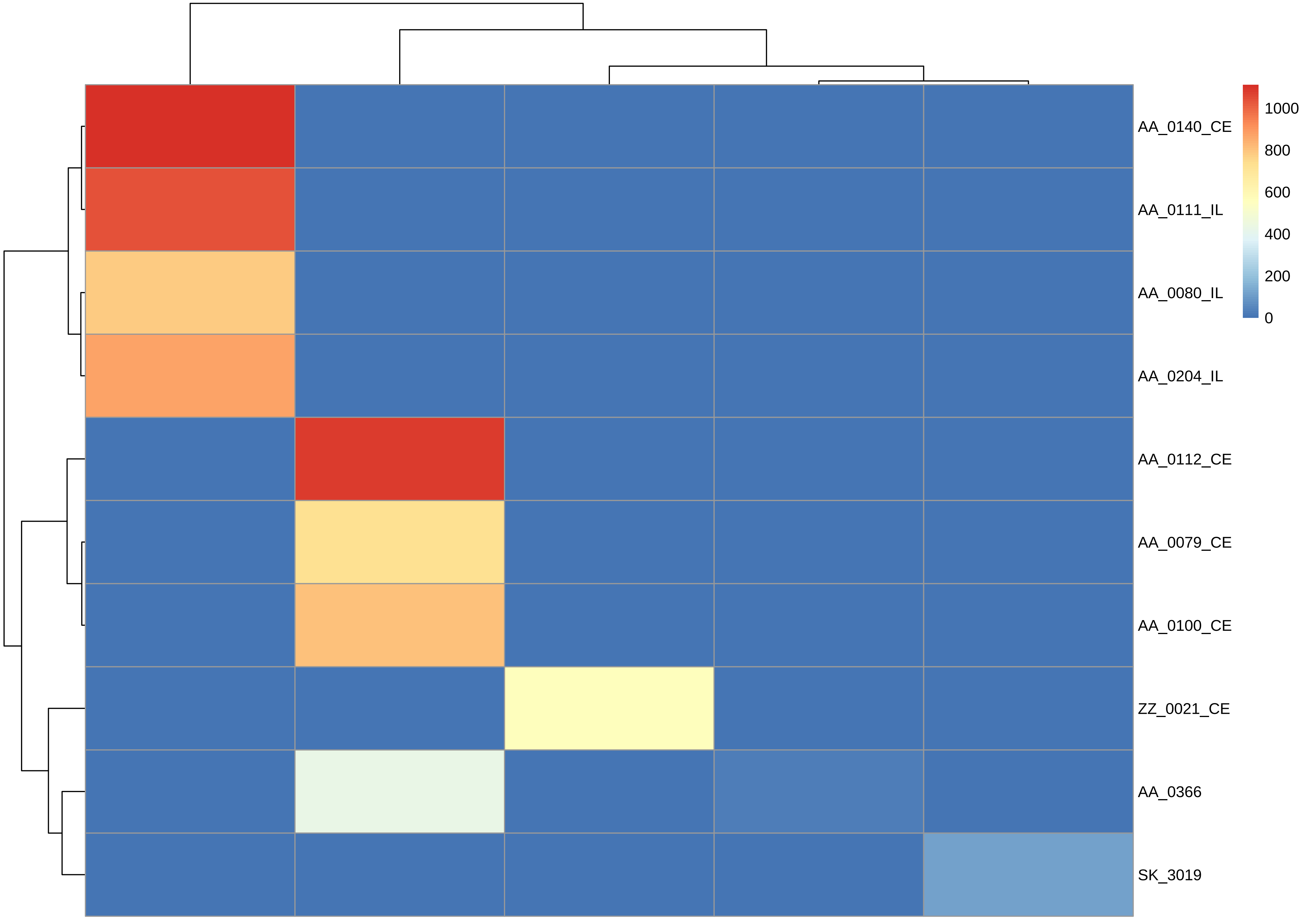

1000
800
600
400
200
0
AA_0140_CE
AA_0111_IL
AA_0080_IL
AA_0204_IL
AA_0112_CE
AA_0079_CE
AA_0100_CE
ZZ_0021_CE
AA_0366
SK_3019

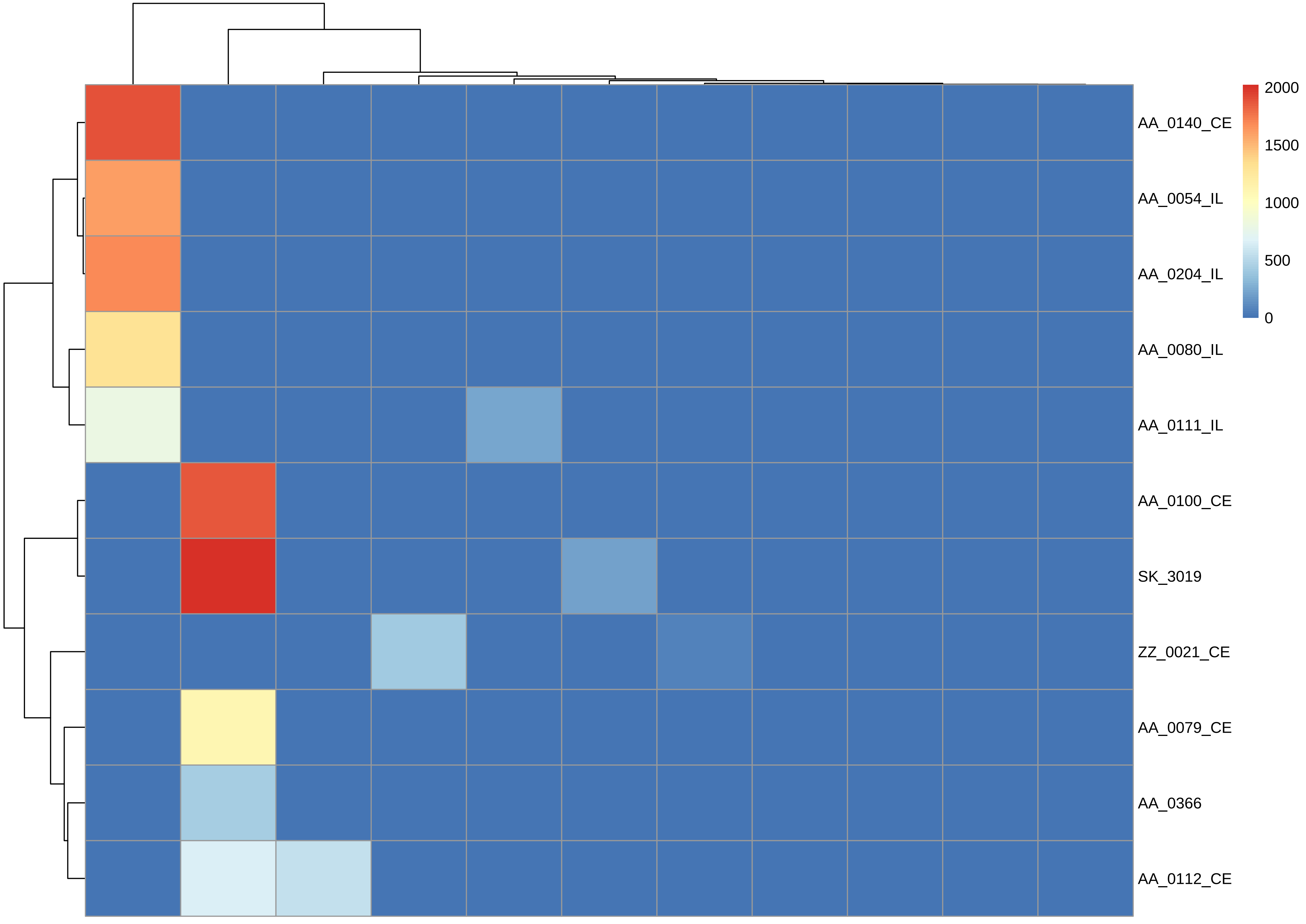

AA_0140_CE
AA_0054_IL
AA_0204_IL
AA_0080_IL
AA_0111_IL
AA_0100_CE
SK_3019
ZZ_0021_CE
AA_0079_CE
AA_0366
AA_0112_CE
2000
1500
1000
500
0

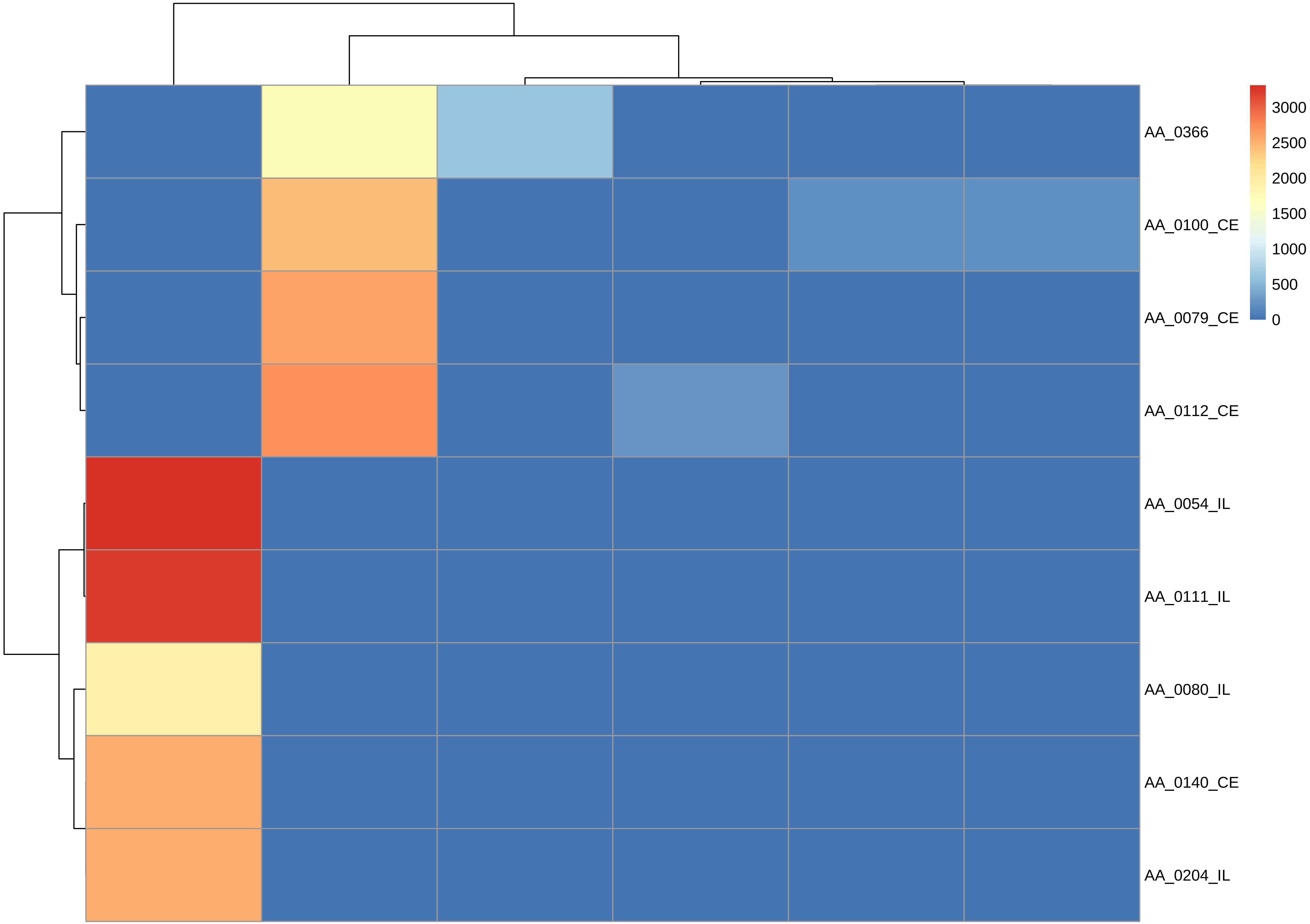
AA_0366
AA_0100_CE
AA_0079_CE
AA_0112_CE
AA_0054_IL
AA_0111_IL
AA_0080_IL
AA_0140_CE
AA_0204_IL
3000
2500
2000
1500
1000
500
0

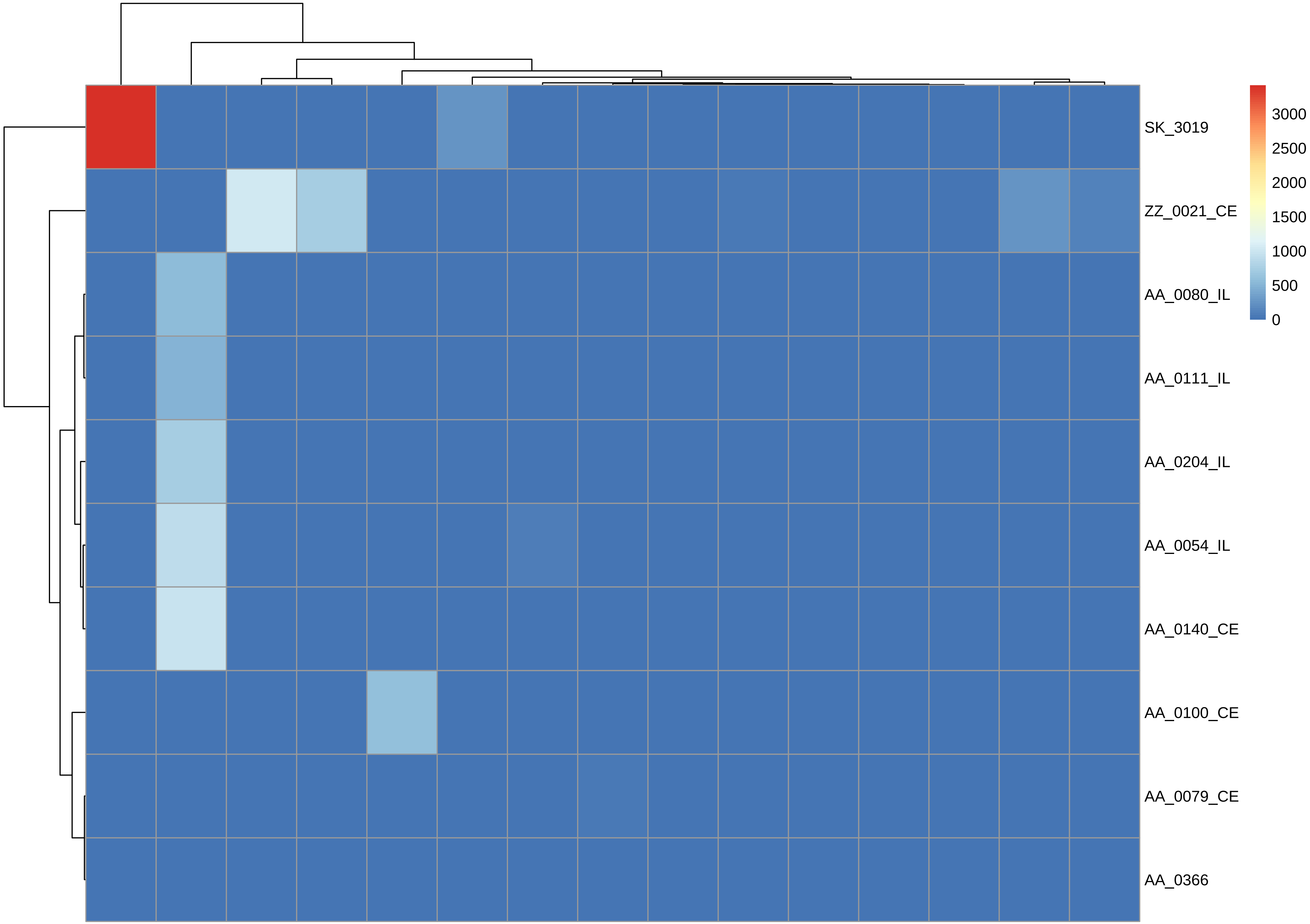

SK_3019
ZZ_0021_CE
AA_0080_IL
AA_0111_IL
AA_0204_IL
AA_0054_IL
AA_0140_CE
AA_0100_CE
AA_0079_CE
AA_0366
3000
2500
2000
1500
1000
500
0

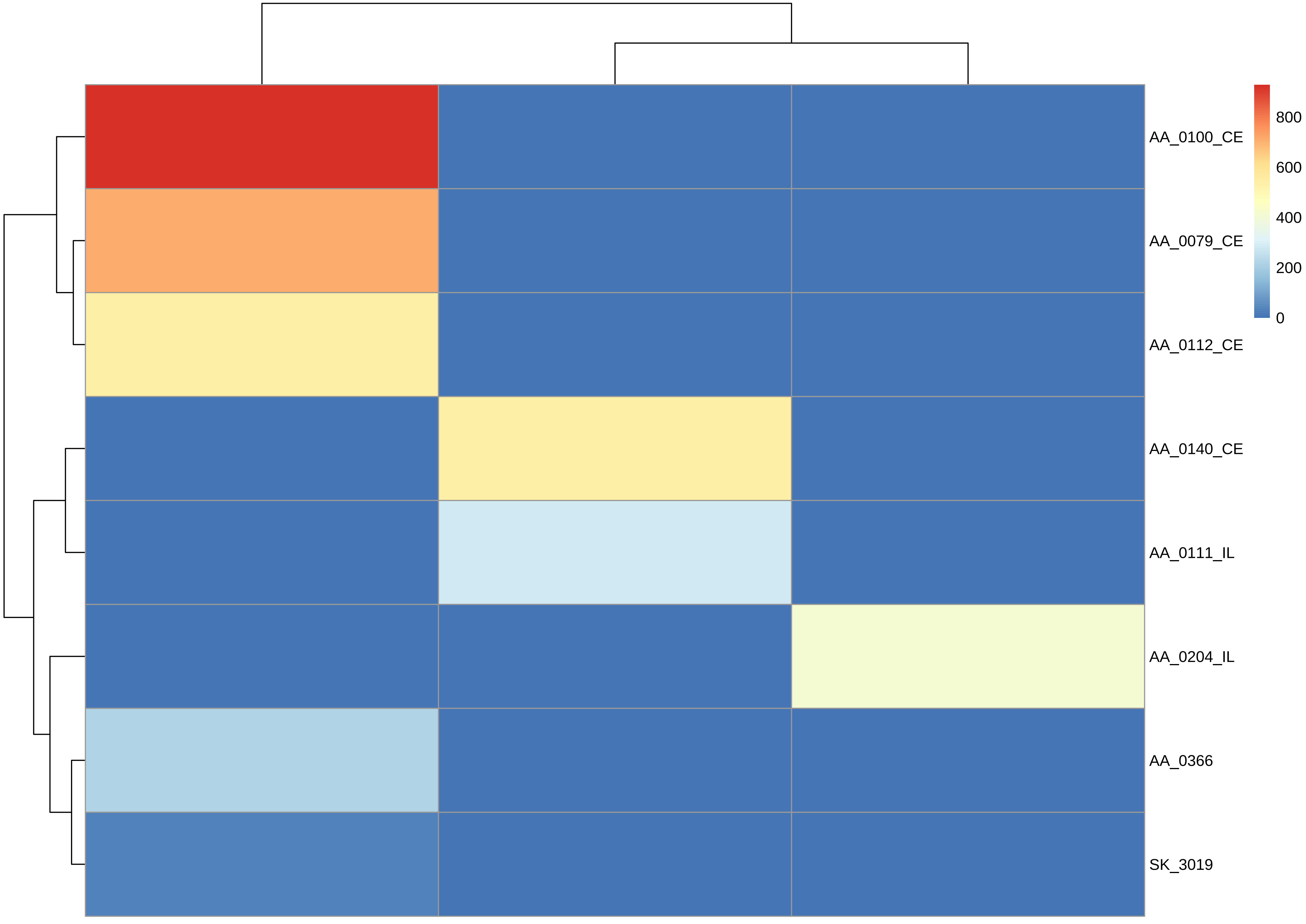
AA_0100_CE
AA_0079_CE
AA_0112_CE
AA_0140_CE
AA_0111_IL
AA_0204_IL
AA_0366
SK_3019
800
600
400
200
0

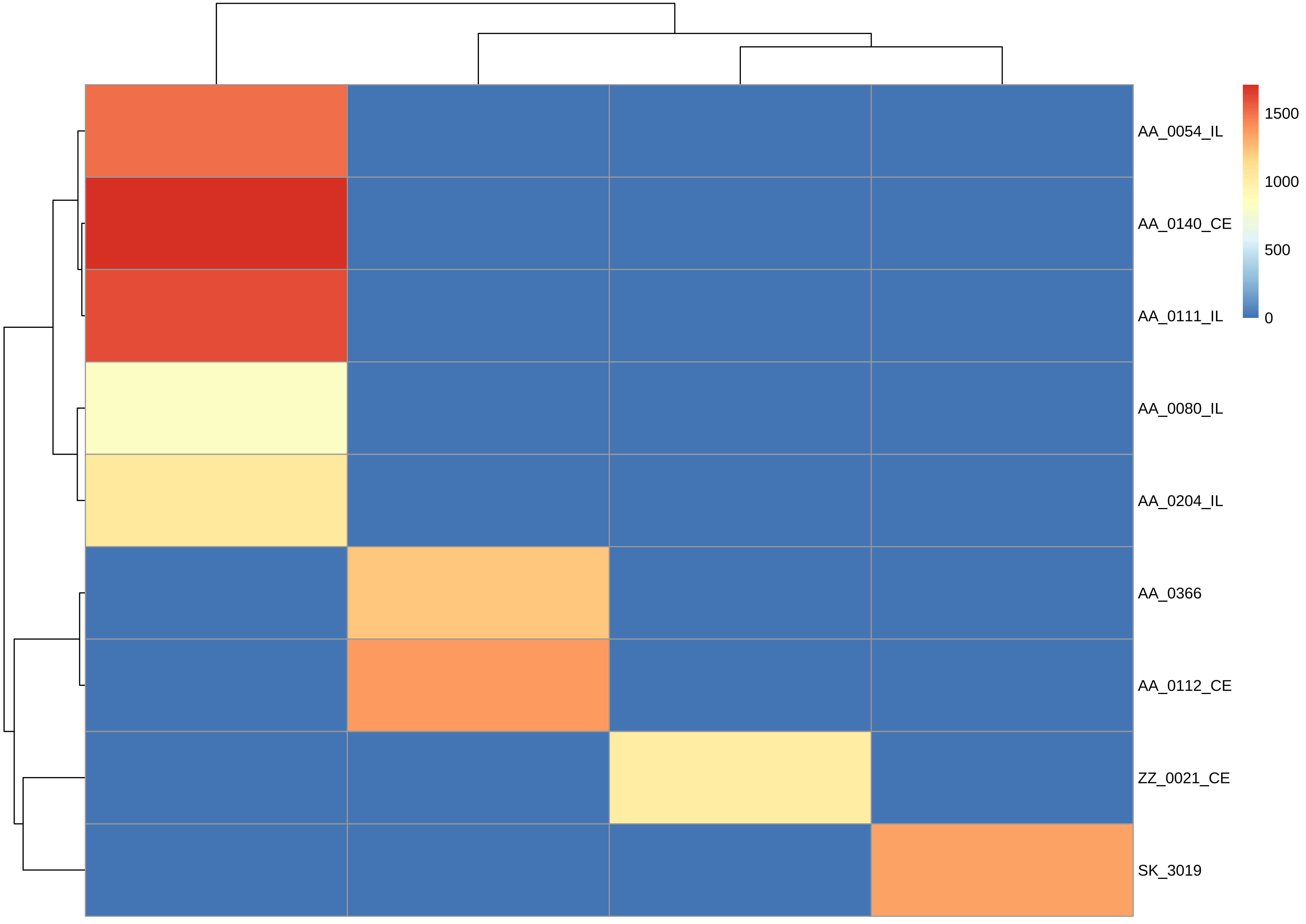

1500
1000
500
0
AA_0054_IL
AA_0140_CE
AA_0111_IL
AA_0080_IL
AA_0204_IL
AA_0366
AA_0112_CE
ZZ_0021_CE
SK_3019

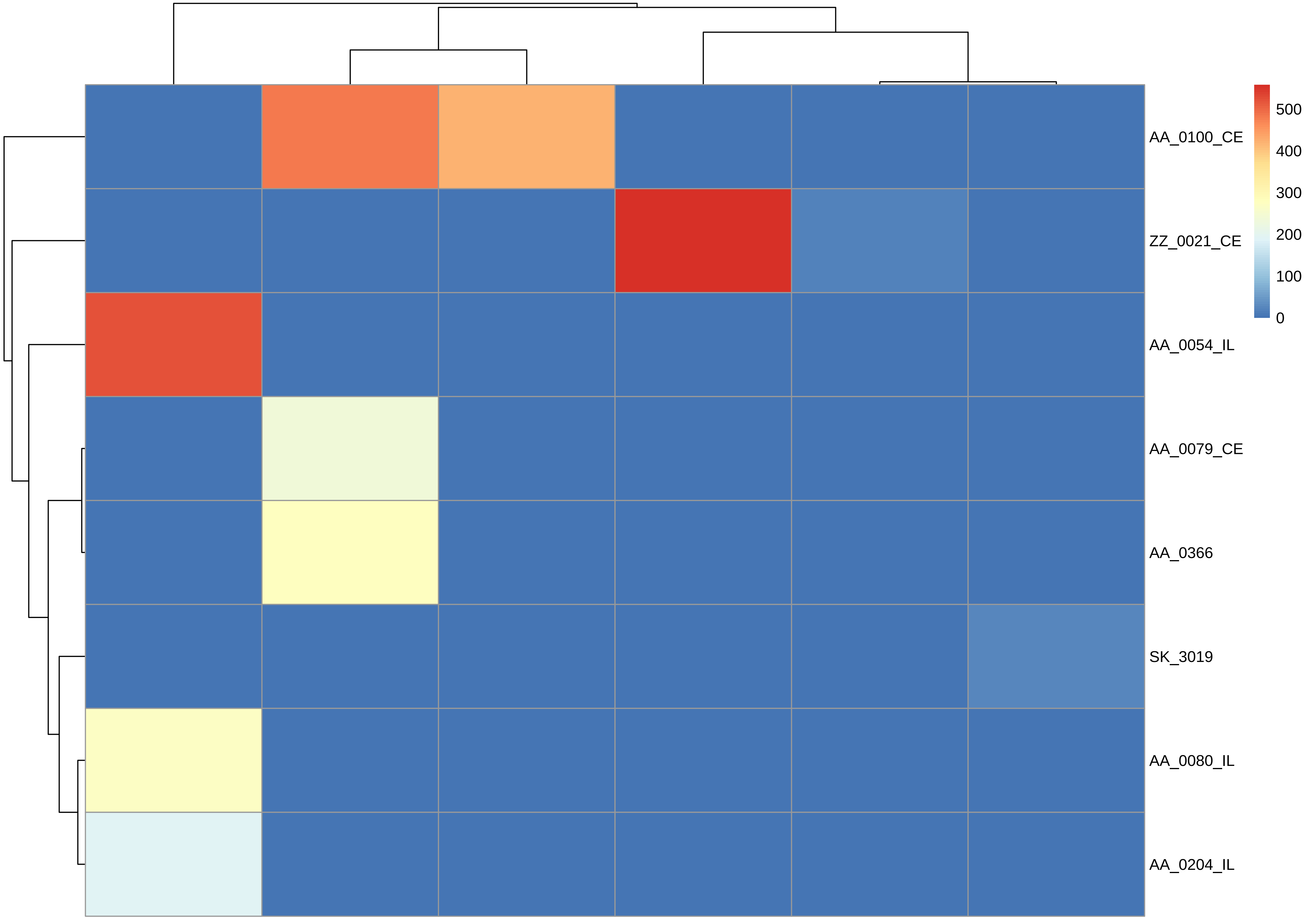
AA_0100_CE
ZZ_0021_CE
AA_0054_IL
AA_0079_CE
AA_0366
SK_3019
AA_0080_IL
AA_0204_IL
500
400
300
200
100
0

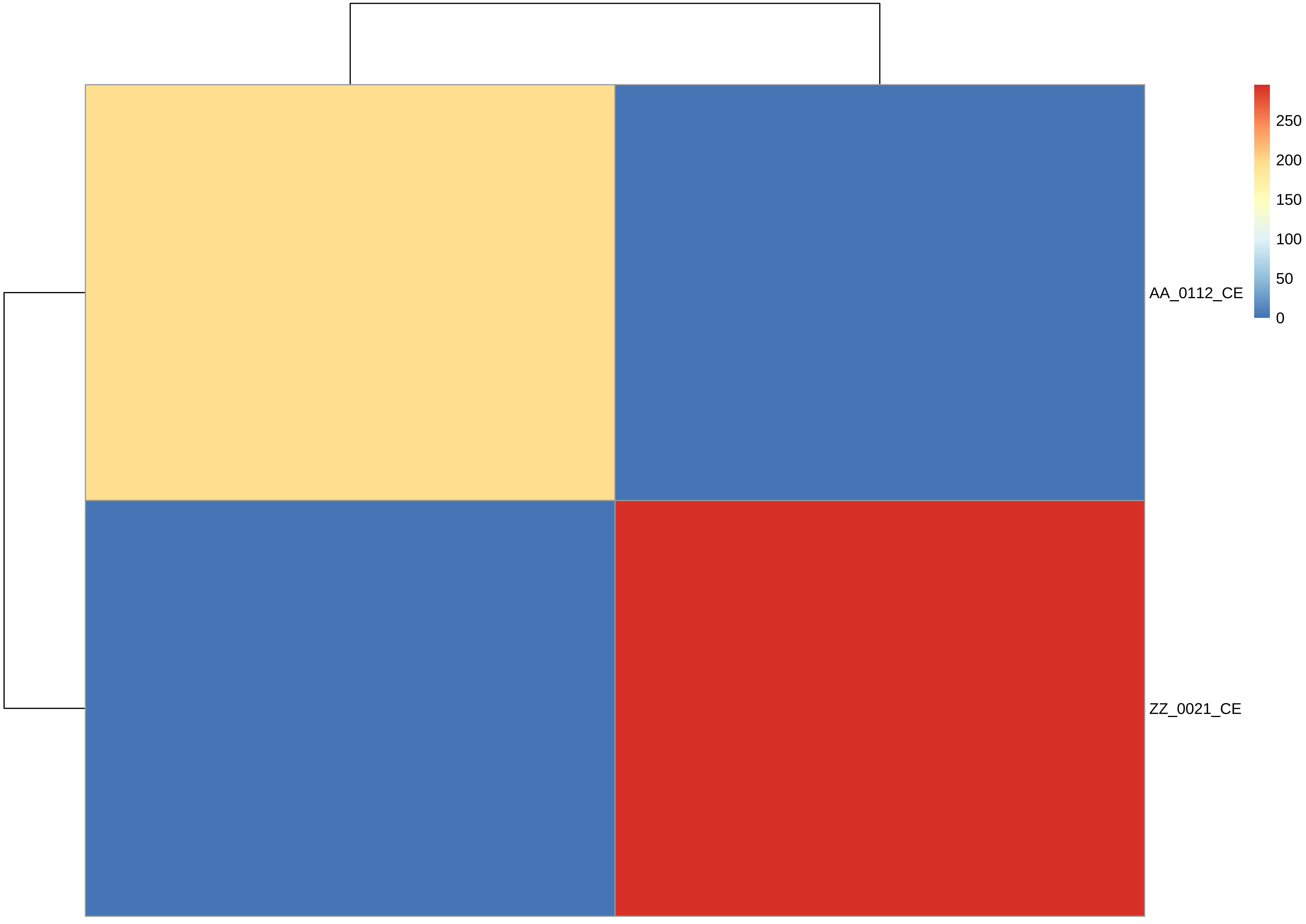
AA_0112_CE
ZZ_0021_CE
250
200
150
100
50
0

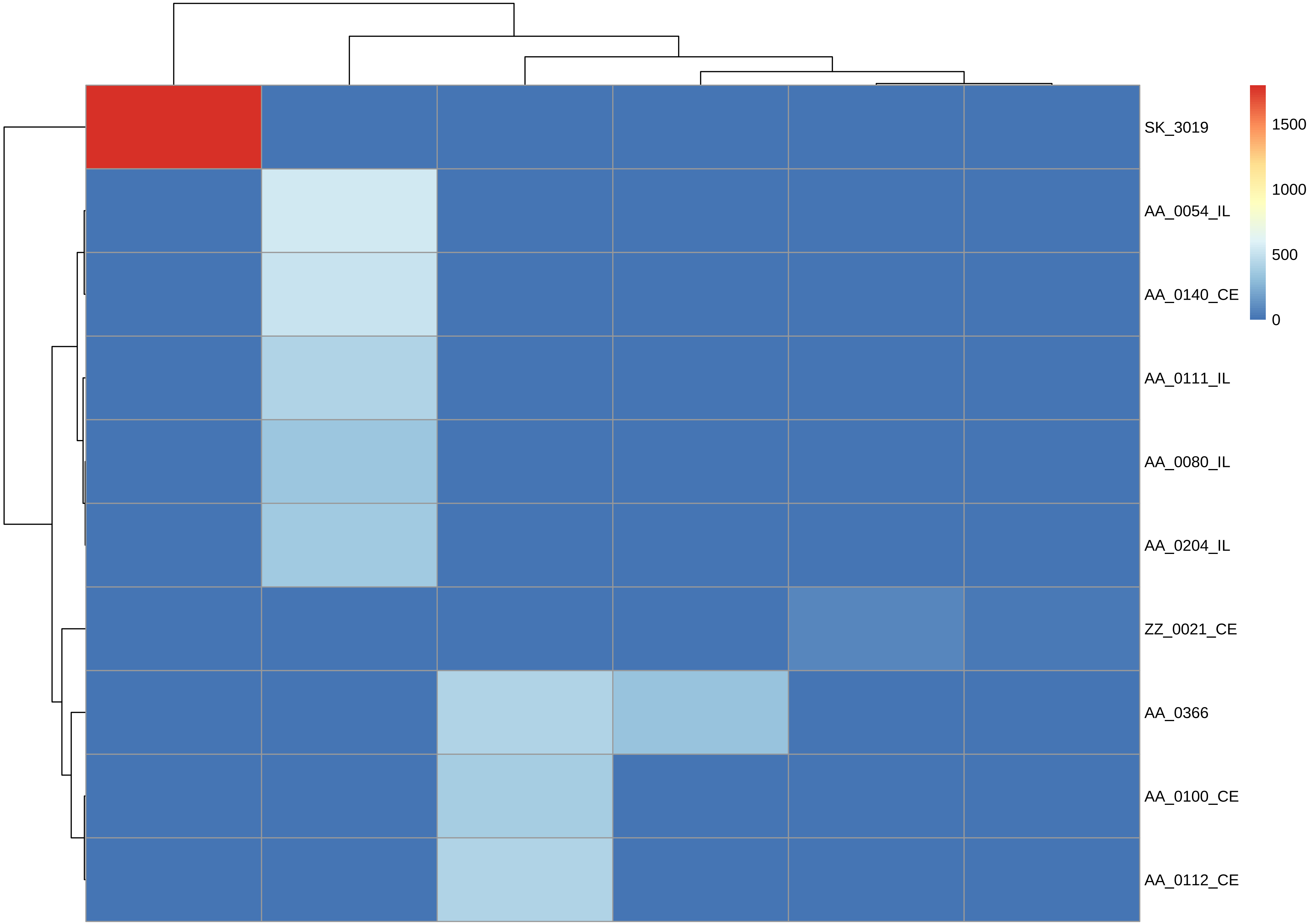
SK_3019
AA_0054_IL
AA_0140_CE
AA_0111_IL
AA_0080_IL
AA_0204_IL
ZZ_0021_CE
AA_0366
AA_0100_CE
AA_0112_CE
1500
1000
500
0

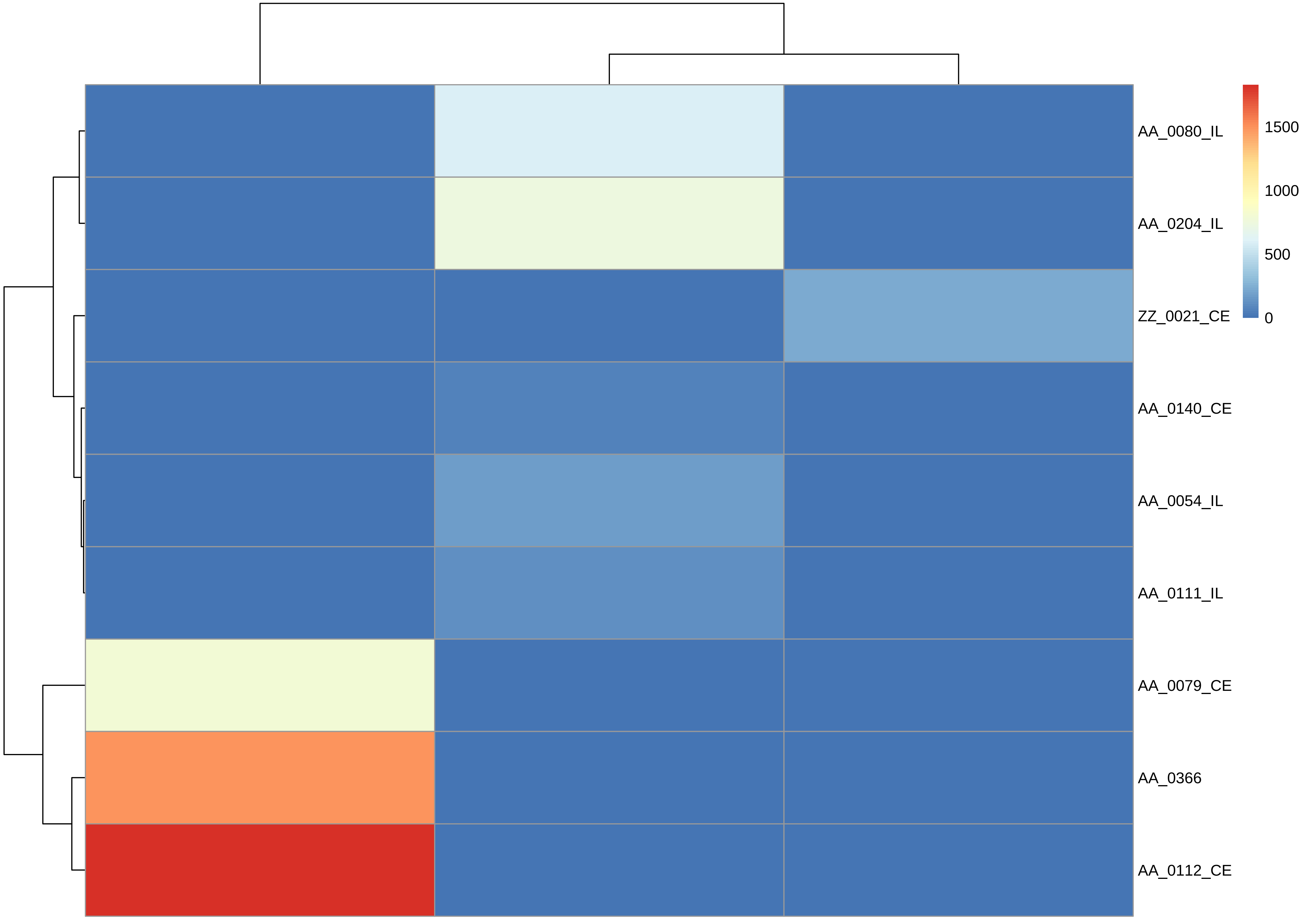
AA_0080_IL
AA_0204_IL
ZZ_0021_CE
AA_0140_CE
AA_0054_IL
AA_0111_IL
AA_0079_CE
AA_0366
AA_0112_CE
1500
1000
500
0

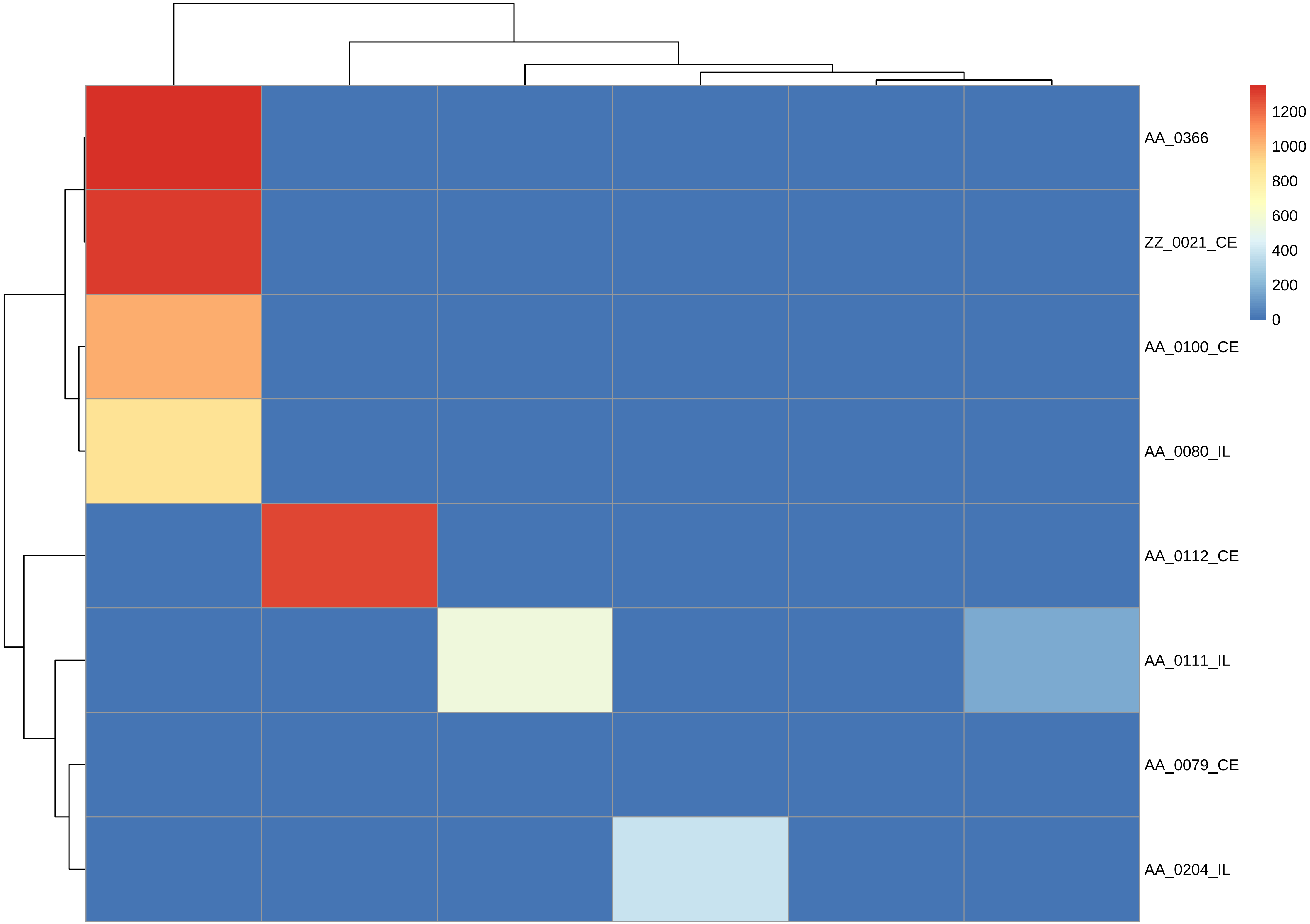

AA_0366
ZZ_0021_CE
AA_0100_CE
AA_0080_IL
AA_0112_CE
AA_0111_IL
AA_0079_CE
AA_0204_IL
1200
1000
800
600
400
200
0

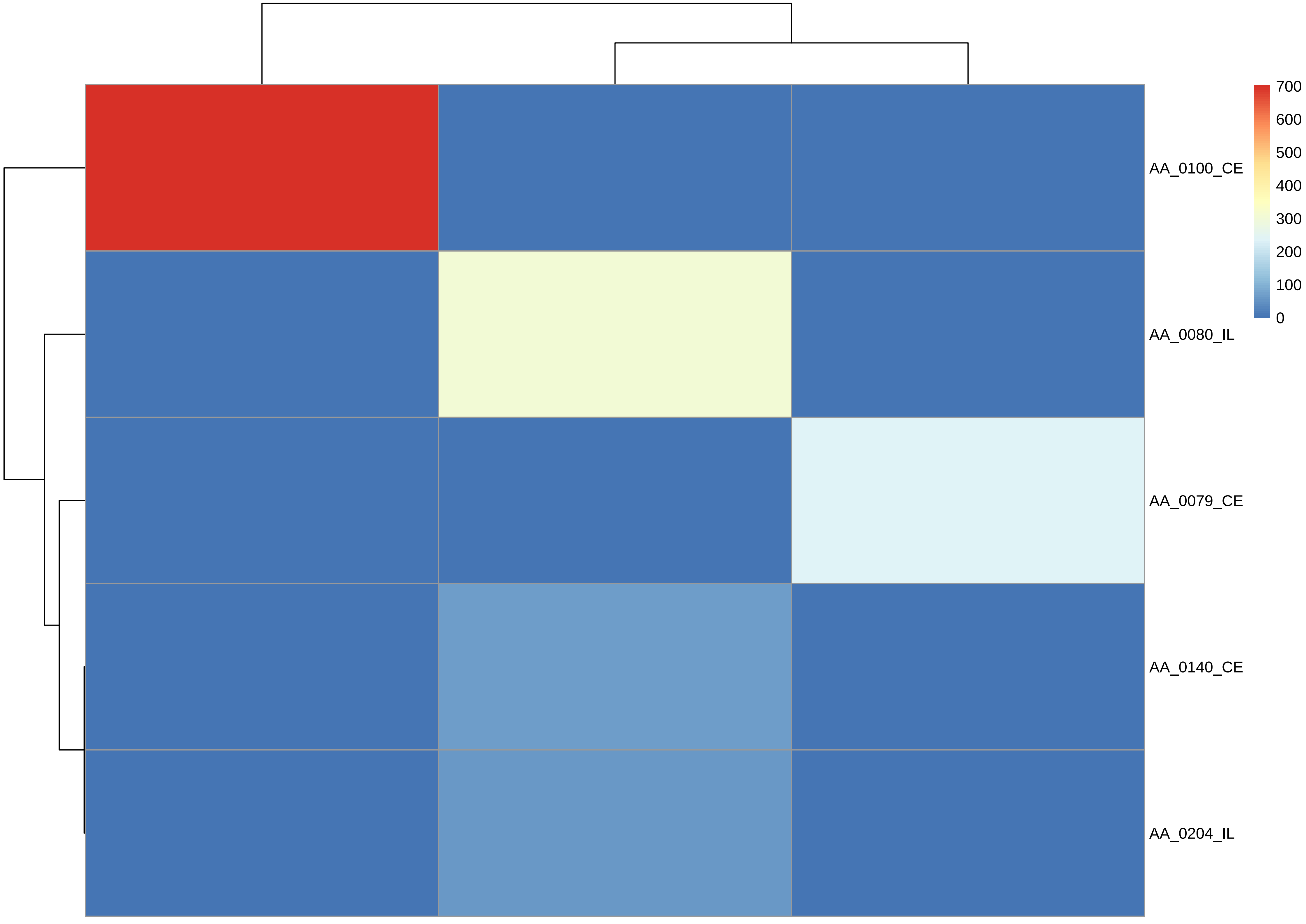
AA_0100_CE
AA_0080_IL
AA_0079_CE
AA_0140_CE
AA_0204_IL
700
600
500
400
300
200
100
0

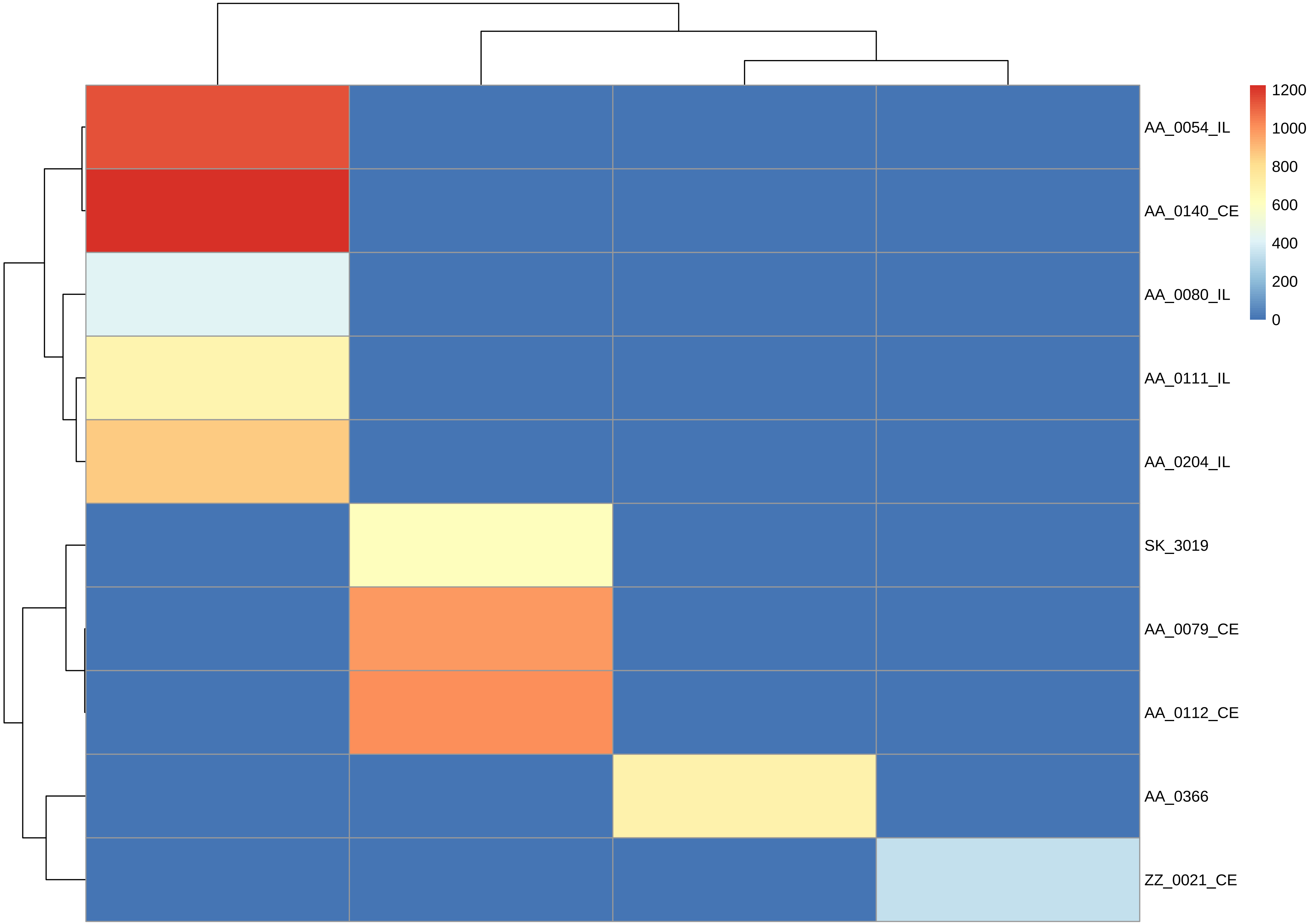

1200
1000
800
600
400
200
0
AA_0054_IL
AA_0140_CE
AA_0080_IL
AA_0111_IL
AA_0204_IL
SK_3019
AA_0079_CE
AA_0112_CE
AA_0366
ZZ_0021_CE

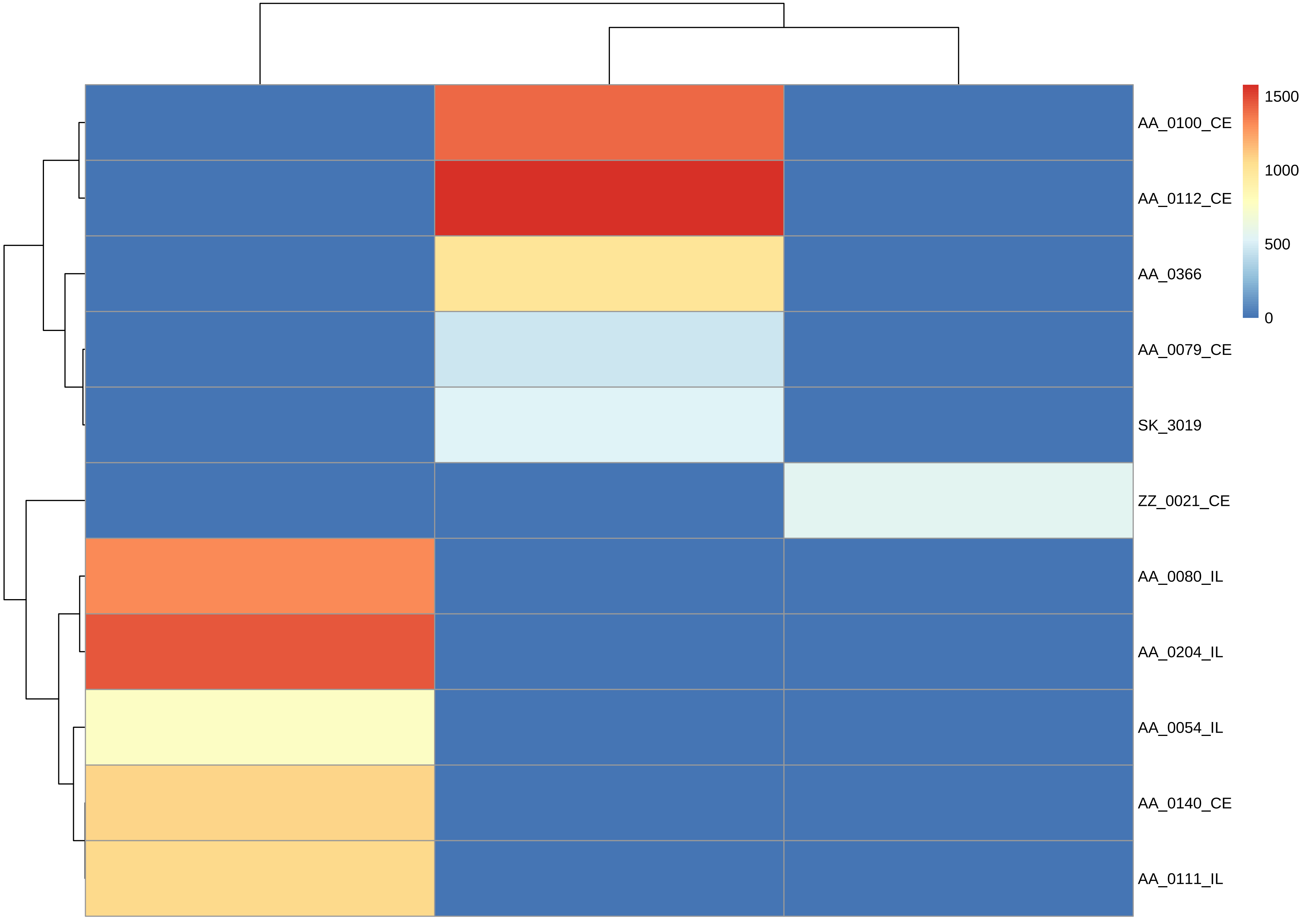
1500
1000
500
0
AA_0100_CE
AA_0112_CE
AA_0366
AA_0079_CE
SK_3019
ZZ_0021_CE
AA_0080_IL
AA_0204_IL
AA_0054_IL
AA_0140_CE
AA_0111_IL

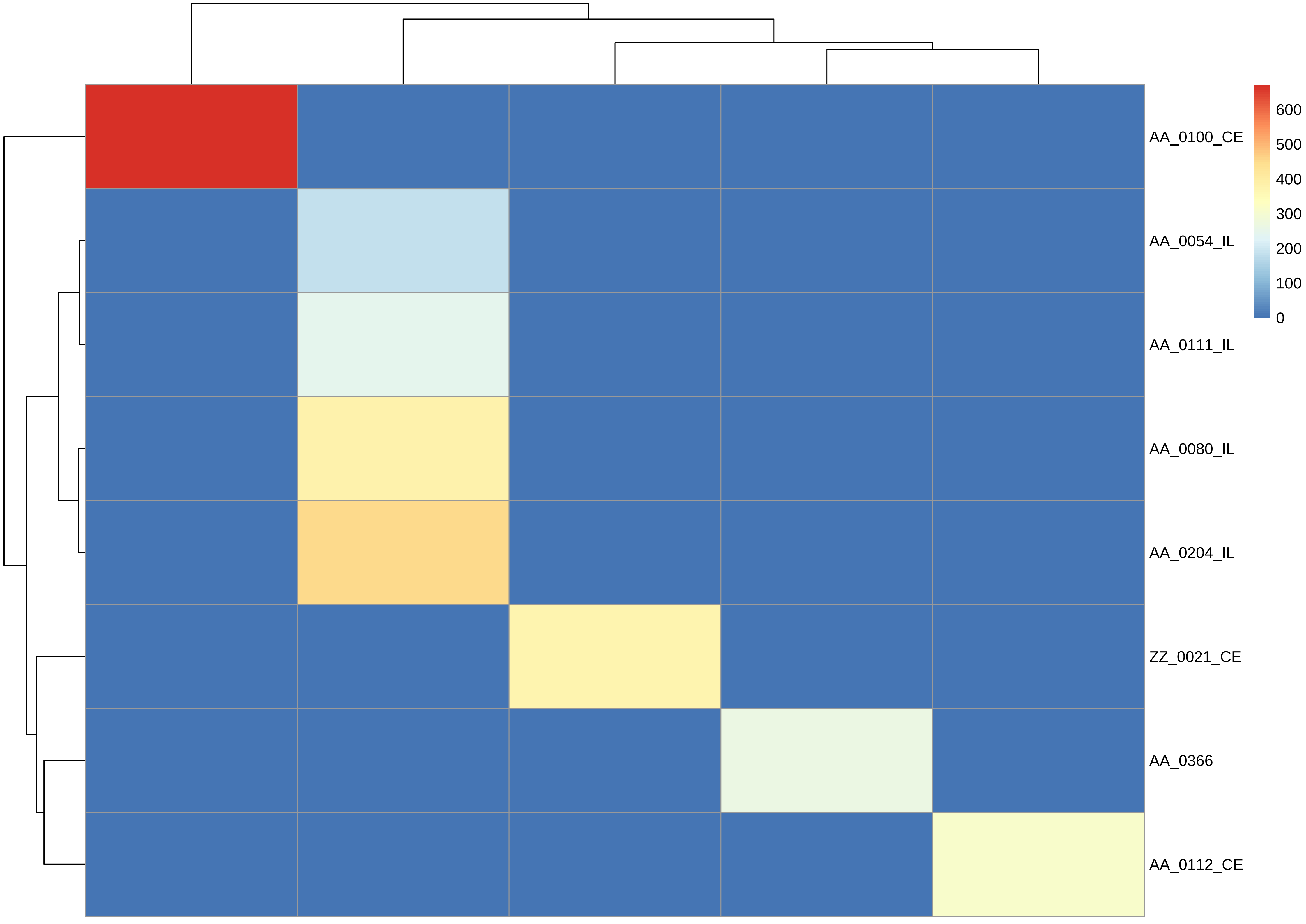

AA_0100_CE
AA_0054_IL
AA_0111_IL
AA_0080_IL
AA_0204_IL
ZZ_0021_CE
AA_0366
AA_0112_CE
600
500
400
300
200
100
0

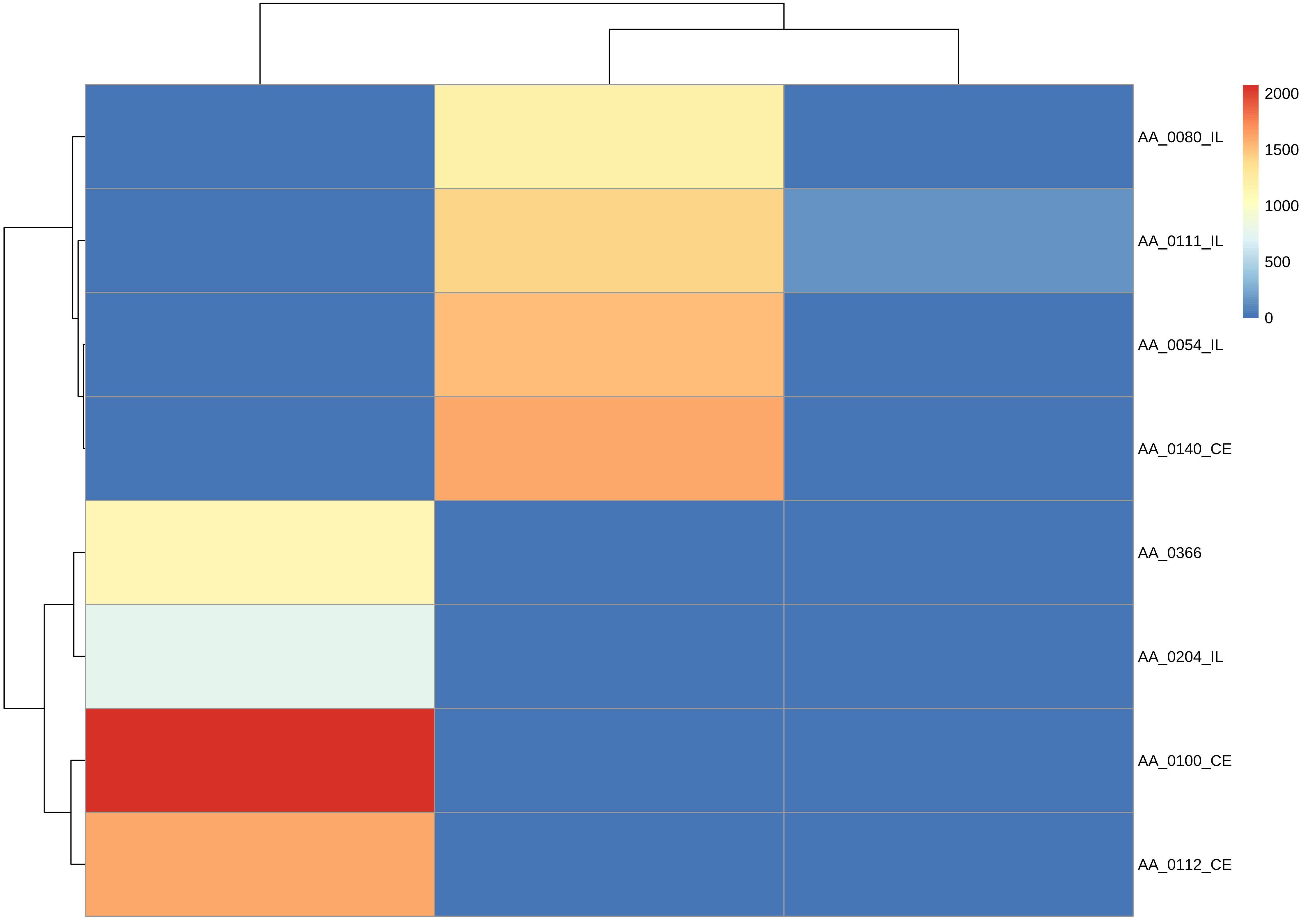
AA_0080_IL
AA_0111_IL
AA_0054_IL
AA_0140_CE
AA_0366
AA_0204_IL
AA_0100_CE
AA_0112_CE
2000
1500
1000
500
0

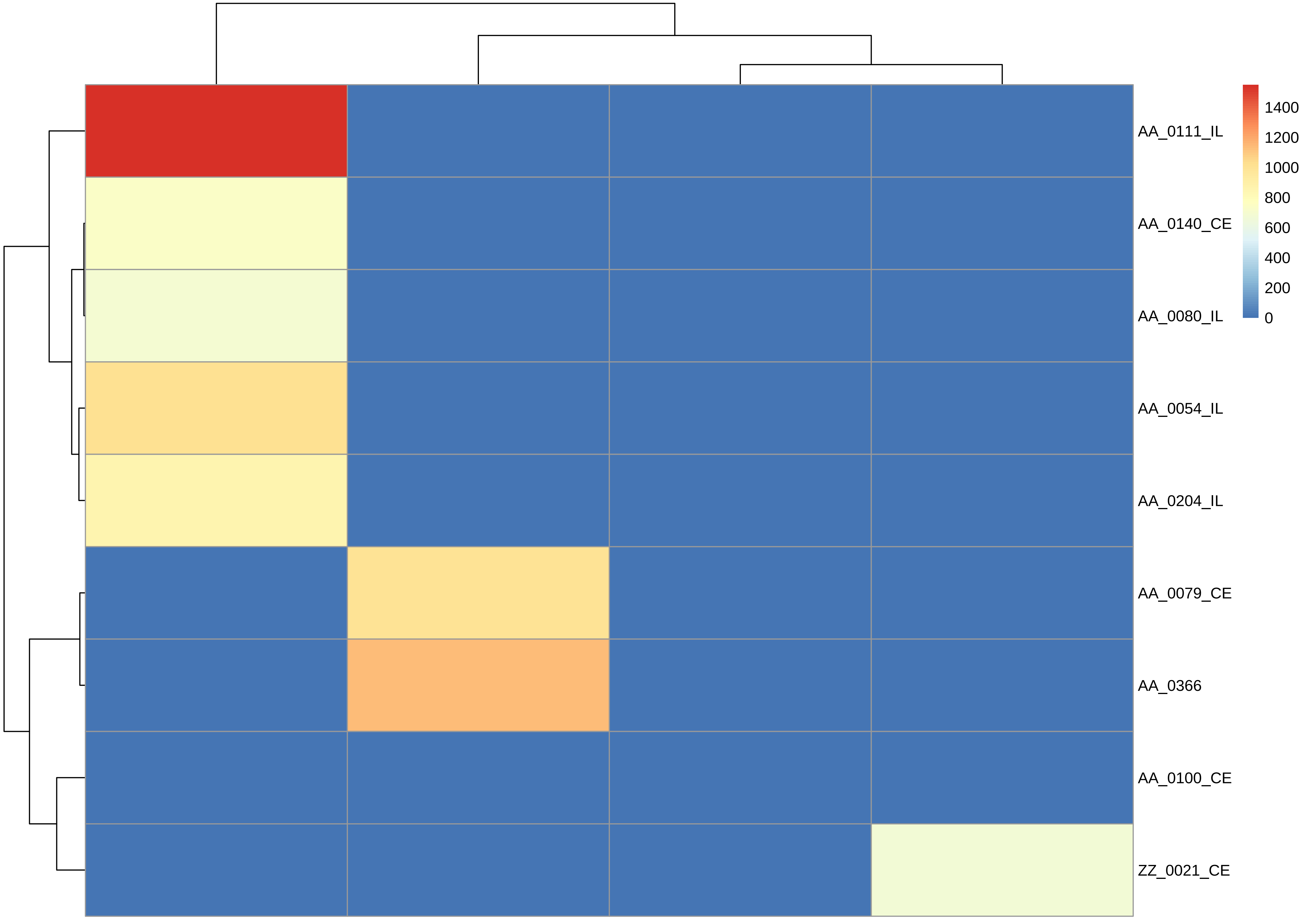

AA_0111_IL
AA_0140_CE
AA_0080_IL
AA_0054_IL
AA_0204_IL
AA_0079_CE
AA_0366
AA_0100_CE
ZZ_0021_CE
1400
1200
1000
800
600
400
200
0

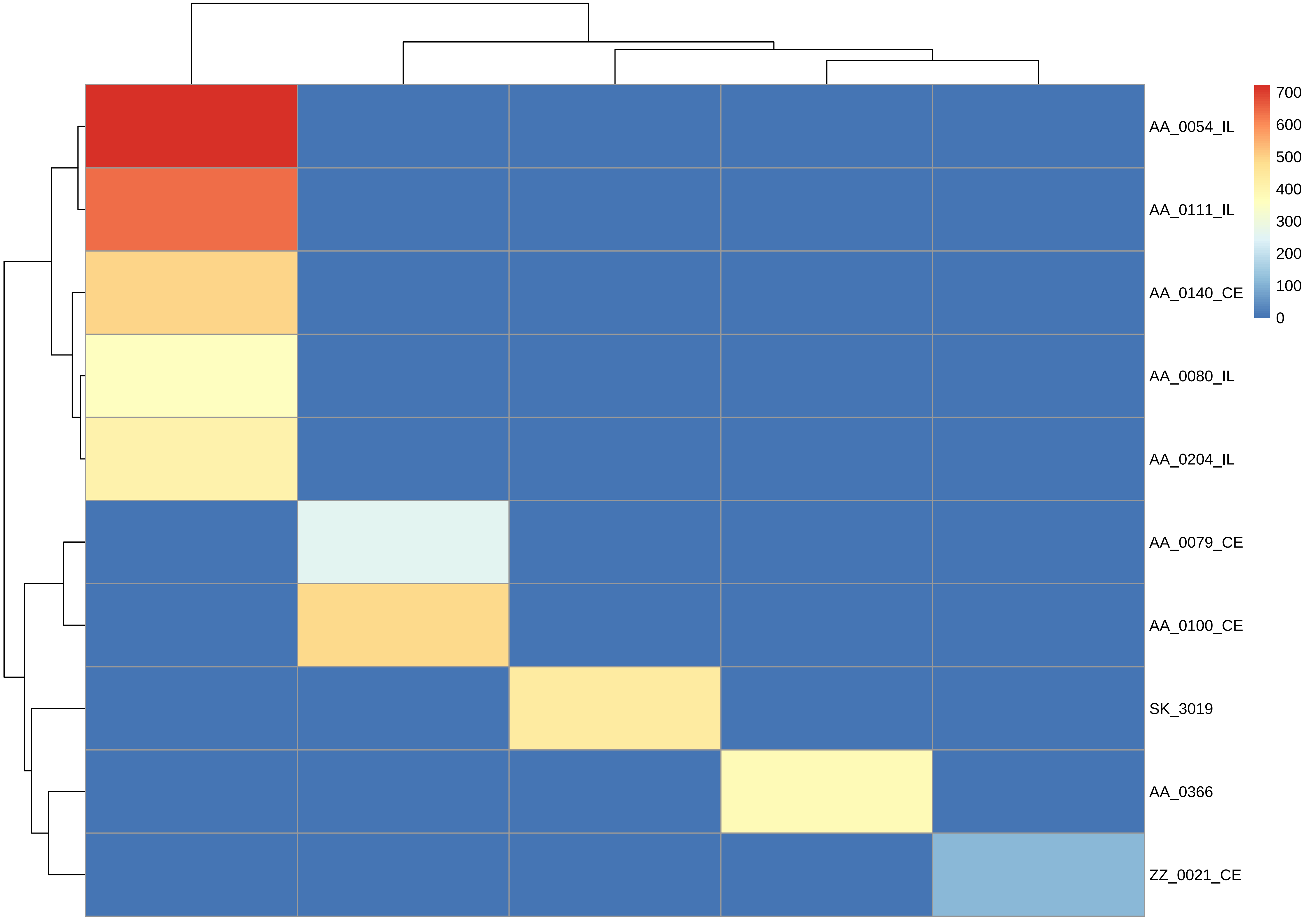

700
600
500
400
300
200
100
0
AA_0054_IL
AA_0111_IL
AA_0140_CE
AA_0080_IL
AA_0204_IL
AA_0079_CE
AA_0100_CE
SK_3019
AA_0366
ZZ_0021_CE

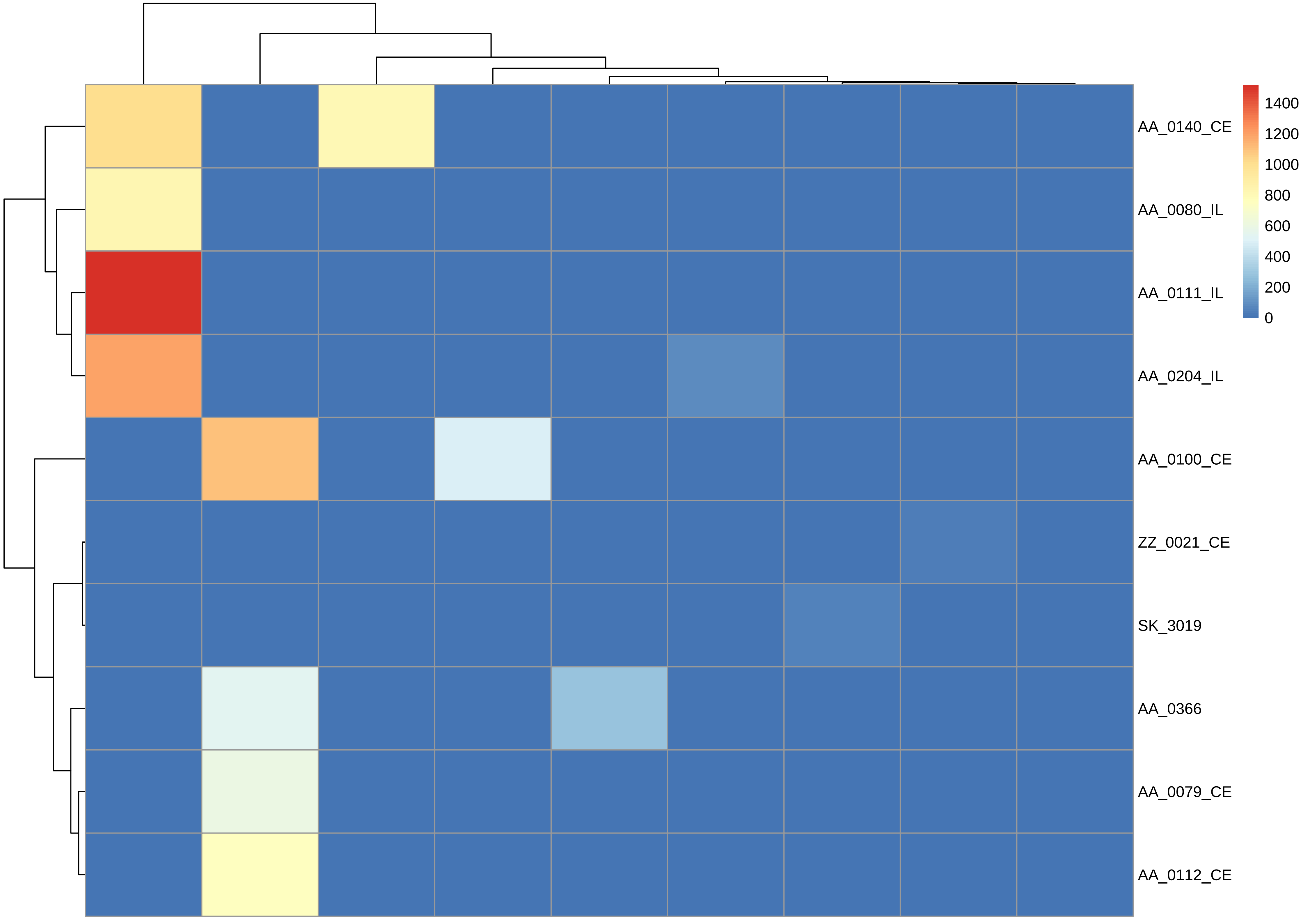

AA_0140_CE
AA_0080_IL
AA_0111_IL
AA_0204_IL
AA_0100_CE
ZZ_0021_CE
SK_3019
AA_0366
AA_0079_CE
AA_0112_CE
1400
1200
1000
800
600
400
200
0

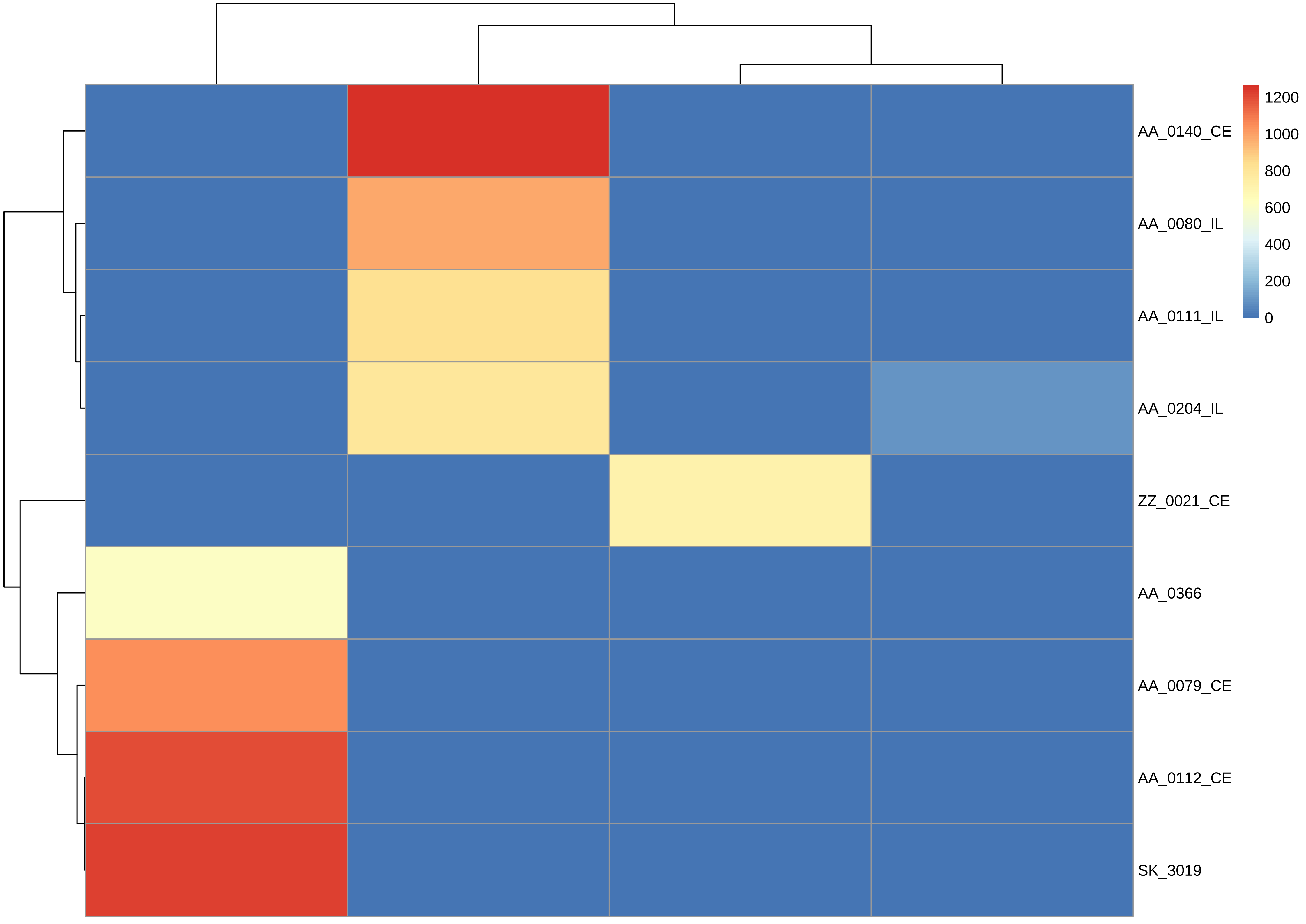
AA_0140_CE
AA_0080_IL
AA_0111_IL
AA_0204_IL
ZZ_0021_CE
AA_0366
AA_0079_CE
AA_0112_CE
SK_3019
1200
1000
800
600
400
200
0
